# Computational prediction resolves thousands of homooligomeric phage protein structures

**DOI:** 10.64898/2026.05.24.727406

**Authors:** Susanna R. Grigson, Natia Geliashvili, Torsten Schubert, George Bouras, Vijini Mallawaarachchi, Marta Bogacz, Ute A. Hellmich, Robert A. Edwards, Bas E. Dutilh

**Affiliations:** Flinders Accelerator for Microbiome Exploration, College of Science and Engineering, Flinders University, Adelaide, SA, 5042, Australia; DOE Joint Genome Institute, Lawrence Berkeley National Laboratory, Berkeley, CA, USA; Institute of Biodiversity, Ecology, and Evolution; Faculty of Biological Sciences, Cluster of Excellence Balance of the Microverse, Friedrich Schiller University Jena, Jena, Germany; Cluster of Excellence Balance of The Microverse, Friedrich Schiller University, Jena, Germany; School of Medicine, College of Health, Adelaide University, Adelaide, SA, 5005, Australia; The Department of Surgery - Otolaryngology Head and Neck Surgery, University of Adelaide and the Basil Hetzel Institute for Translational Health Research, Central Adelaide Local Health Network, Adelaide, SA, 5005, Australia; Institute of Organic Chemistry & Macromolecular Chemistry (IOMC), Cluster of Excellence Balance of the Microverse, Friedrich Schiller University Jena, Jena, Germany; Theoretical Biology and Bioinformatics, Science4Life, Utrecht University, Utrecht, the Netherlands

## Abstract

Bacteriophages (phages) play essential roles in microbial systems, yet most phage proteins remain poorly characterised. Protein tertiary and quaternary structure information contributes valuable information about protein function. As many phage proteins function as homooligomers, complexes that consist of multiple identical subunits, there is great interest in computationally predicting their configurations. Here we present a computational framework, the Phage Homomer Level Estimate and Generation Method (PHLEGM) for inferring homooligomeric states directly from the protein sequence by combining AlphaFold-Multimer modelling with inter-subunit interface quality assessment. We proceeded to experimentally validate two out of nine predicted homooligomers using size exclusion chromatography and complementary hydrodynamic techniques. These efforts confirmed our predictions for a dimer and a trimer, highlighting the value of experimentally benchmarked computational predictions and showing the challenges of heterologous phage protein production. Applied to >22,000 phage protein sequences in the PHROGs database, our approach revealed extensive diversity in phage homooligomeric protein complexes. Benchmarking against protein language model-based predictors on a curated reference set of known phage homooligomers demonstrated superior accuracy of our structure-based method, achieving robust performance in classifying protein homooligomeric states, with the highest accuracy observed for trimers and higher-order complexes. These results highlight the value of computational predictions to decipher the complexities of the vast viral sequence space. All predicted complex structures and functional inferences are made publicly available to support structural and functional studies of phage proteins.

## Introduction

Bacteriophage (phage) genomes encode a wide range of structural, enzymatic, regulatory, and other proteins that drive every stage of the infection cycle, from host recognition and genome delivery to replication, assembly, and lysis. Advances in sequencing technologies^1^, genomic databases^2–5^, and automated annotation tools^6–8^ have rapidly expanded the catalogue of known phage proteins, with recent efforts identifying over 24 million viral genomes and cataloguing nearly 40,000 distinct viral protein families^4^. Despite this growth, approximately 65% of phage proteins remain functionally uncharacterised^8^, limiting our understanding of phage biology, host interactions, and phage biotechnological potential.

Across scales, biology repeatedly converges on symmetry as the shortest path from simple rules to complex, reliable forms^9^. Phages are constrained by their compact genomes yet tasked with building large mechanical machines. To achieve high structural complexity while maintaining genomic economy, phages leverage homooligomerisation, assembling identical protein subunits into homooligomeric complexes to achieve their functional forms^10–12^. A protein’s homooligomeric state refers to the specific number of subunits in these complexes (e.g. dimers, trimers, hexamers). Functional homooligomerisation is essential for many phage proteins: capsid proteins form tri-, penta-, and hexameric units to build icosahedral shells^13,14^; tail fibres mediate host attachment as trimers^15,16^; and enzymes such as endolysins and integrases depend on specific homooligomeric states for their catalytic activity^17,18^. The homooligomeric state of a protein can therefore provide clues to its function. Experimental methods, such as size-exclusion chromatography (SEC), cross-linking mass spectrometry, and high-resolution structural techniques (e.g., cryo-electron microscopy), can resolve homooligomeric states but are resource-intensive and often impractical for phage proteins, which are often small, unstable, or challenging to express^19–21^. As a result, the homooligomeric states of most phage proteins remain undetermined.

Recent breakthroughs in computational structural biology, most notably AlphaFold^22,23^, now enable frequent accurate prediction of three-dimensional protein tertiary structures directly from sequence. When paired with structural comparison tools such as Foldseek^24^, these predicted monomeric structures can be compared against large databases to identify structural homologues with related functions^24,25^. For instance, the phage annotation tool Phold^26^ combines Foldseek with a comprehensive database of predicted phage protein structures to extend functional annotation beyond sequence similarity, enabling annotation of roughly half of the genes in a typical phage genome^26^. Although highly effective, relying solely on tertiary structure overlooks the biological reality that many phage proteins only reach their mature functional forms as complex, higher-order assemblies. To address multi-chain complexes, AlphaFold-Multimer^27^, was developed to facilitate prediction of quaternary protein structures. Recently, this technology was deployed at scale to expand the AlphaFold Protein Structure Database with proteome-wide quaternary predictions^28^ and the Viral AlphaFold Database of monomers and homodimers^29^. However, these large-scale efforts utilise a dimer-only modelling strategy, neglecting higher-order configurations.

While AlphaFold-Multimer’s interface metrics can theoretically deduce assembly stoichiometries, the precise number of subunits in a complex, systematic application remains computationally expensive, particularly for large datasets of phage proteins^30–33^. To bypass this high computational cost, alternative methods including QUaternary state prediction using dEEp learNing (QUEEN)^34^, Seq2Symm^35^, and Stoic^33^ rely on protein language models. While these sequence-based approaches are highly efficient, they have yet to be rigorously benchmarked on phage proteins.

Here, we present a systematic, structure-informed framework for predicting the homooligomeric states of phage proteins from amino acid sequences. Our approach integrates AlphaFold-Multimer modelling across a range of subunit configurations with quantitative interface scoring to identify the most probable homooligomeric states. Applied to over 22,000 structurally representative proteins from the Prokaryotic Virus Remote Homologous Groups (PHROGs) database^36^, our method reveals widespread and diverse multimeric architectures across the phage proteome. The predicted homooligomeric states are consistent with established structure–function relationships and display distinct patterns of enrichment across functional categories. Comparative benchmarking demonstrates that our structure-informed framework outperforms state-of-the-art protein language models, with particularly strong gains for complex higher-order assemblies. All predicted assemblies are publicly available, providing a resource for structure-guided functional annotation and advancing our understanding of phage biology.

## Results

### Curation of representative phage proteins for homooligomeric state prediction

To enable database-wide prediction of homooligomeric states in phage proteins, we curated representatives from the PHROGs database. This resource provides a comprehensive catalogue of 38,880 phage protein families; however, the majority of these, approximately 87%, belong to the ‘unknown’ category, with the remainder distributed across nine specific functional classes^36^. To reduce redundancy while maintaining structural diversity, all-versus-all structural alignments were performed within each PHROG, and the sequence with the highest average similarity (highest cumulative alignment bitscore) to all other members was selected as the representative sequence of the PHROG. Because GPU memory limits AlphaFold-Multimer predictions for large proteins, only representatives shorter than 350 amino acids were retained, comprising 90% of all PHROG representatives (35,083 sequences; Fig. S1A). The excluded sequences were predominantly large structural proteins, including tail length tape measure proteins, virion structural proteins, and tail fibre (Fig. S1B). Each PHROG representative was first modelled as a monomer using AlphaFold2 to assess fold reliability prior to multimeric modelling. Models with a predicted Local Distance Difference Test (pLDDT) score above 70^22^ were retained, yielding 22,545 high-confidence models (Fig. 1A, S1C). Notably, many high-confidence predictions were obtained from shallow multiple sequence alignments, demonstrating that AlphaFold can generate confident folds even with limited sequence homology (Fig. S1C-D).

**Figure 1:**
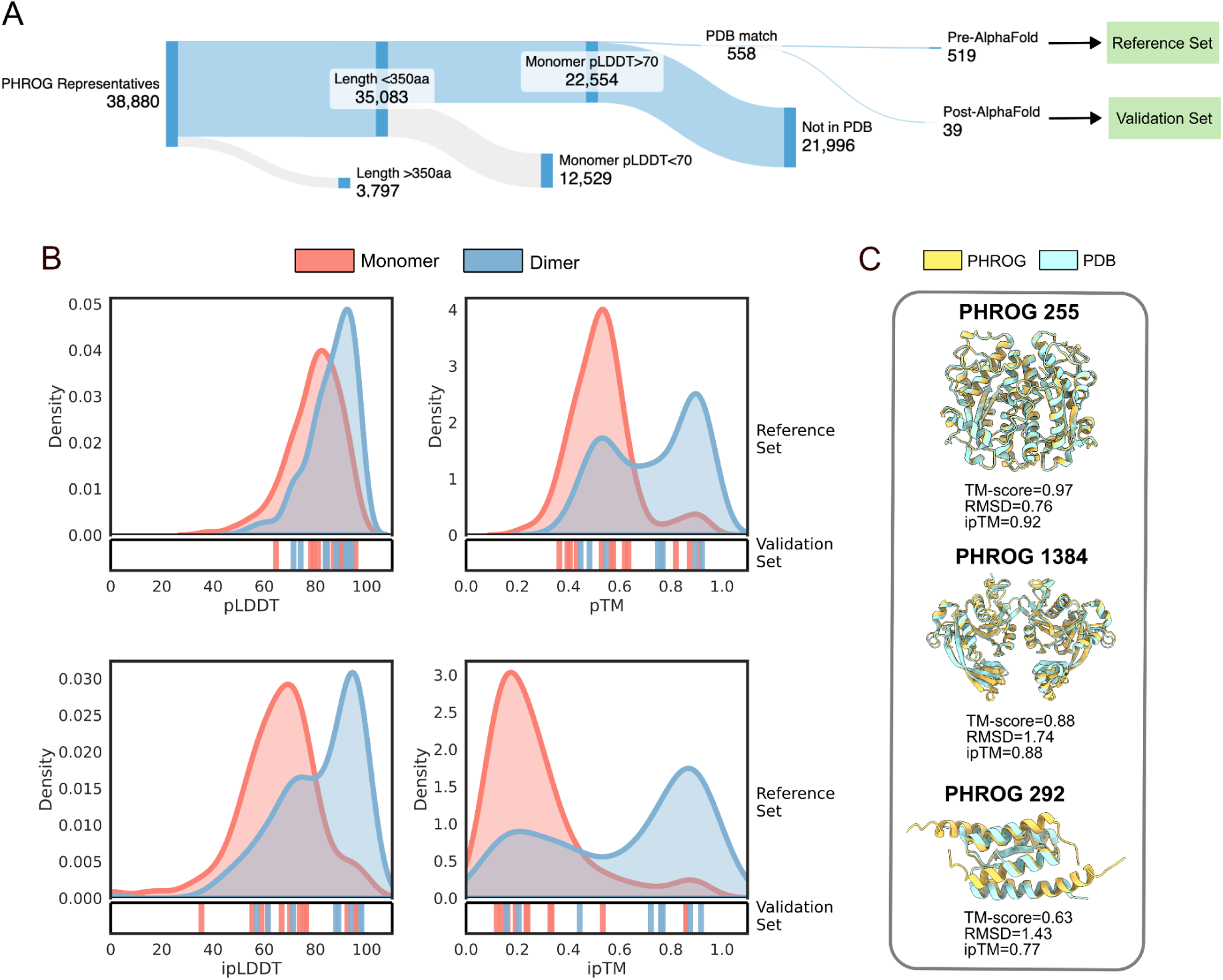
Evaluating AlphaFold-Multimer metrics for predicting biologically relevant interfaces in phage proteins. A. Curation of Prokaryotic Virus Remote Homologous Groups (PHROGs) for homooligomeric state prediction. Diagram generated using SankeyMATIC. B. Density distributions of AlphaFold-Multimer metrics for the reference set of PHROGs with experimental structures in the Protein Data Bank (PDB) crystalised as monomers (n=168) and dimers (n=196). The corresponding scores for monomers (n=12) and dimers (n=8) in the validation set are displayed as rugplots directly beneath each density distribution. C. Structure alignments of AlphaFold-Multimer predictions of PHROG representatives with structures deposited after the Alpha-Multimer training cutoff date: exonuclease PHROG 255 (PDB 8kel), uncharacterised protein PHROG 292 (PDB 8dsb^38^) and Mom-like DNA modification protein PHROG 1384 (PDB 8bv8).

To identify proteins with experimentally validated homooligomeric states, representatives were aligned against the Protein Data Bank (PDB). Homooligomeric states could be assigned to 558 PHROGs based on matching PDB entries with resolved quaternary structures (see Methods). Of these, 519 deposited before the cutoff date of AlphaFold-Multimer v2.3 training (30 September 2021) formed the reference set for threshold optimization and method development. The remaining 39 post-cutoff structures served as an independent test set for evaluating performance on structures unseen during training (Table S1).

### Detecting biologically relevant interfaces in phage proteins with AlphaFold-Multimer

To evaluate AlphaFold-Multimer’s ability to identify biologically relevant interfaces while avoiding false positives, we first benchmarked its predictions against 196 homodimers and 168 monomers from the reference set. Each protein was modelled as a two-chain assembly with five independent predictions. Quality metrics were calculated for each structure, and mean values across the five models were used to assess their capacity to separate genuine interfaces from spurious contacts.

Global structural metrics evaluating the entire complex provided limited discriminative power. The number of inter-subunit contacts and predicted aligned error (PAE), which reflects prediction uncertainty, showed largely overlapping distributions for monomers and dimers (Fig. S2A), while pLDDT offered modest improvement in distinguishing monomers from dimers (Fig. 1B). Metrics capturing interface or complex-level accuracy demonstrated much stronger separation, consistently assigning higher values to dimers than monomers. These included (i) the predicted template modelling score (pTM), which evaluates overall complex accuracy; (ii) interface pLDDT (ipLDDT), which reflects confidence at protein–protein interfaces; (iii) interface pTM (ipTM), which assesses interface structural fidelity; and (iv) the docking confidence score pDockQ2 (31) (Fig. 1B, S2A). Receiver operating characteristic (ROC) and precision–recall (PR) analyses confirmed that ipTM gave the strongest performance for classifying monomers and dimers (AUC = 0.81, AP = 0.85), followed by pTM (AUC = 0.80, AP = 0.84) and pDockQ2 (AUC = 0.79, AP = 0.82) (Fig. S2B-C). Combining multiple metrics via simple machine learning models (e.g., Random Forest, Support Vector Machine) did not enhance predictive performance (Supplementary Note 1), indicating that the different interface scoring methods captured similar information (Fig. S3).

To assess whether interface quality metrics generalise to structures deposited after AlphaFold-Multimer’s training cutoff, we evaluated 20 proteins (12 monomers, 8 dimers) deposited after the training cutoff date (Fig. 1B). Dimers with experimentally validated structures achieved high ipTM scores and closely matched experimental structures. These included PHROG 255 (TM-score = 0.97, ipTM = 0.92), PHROG 1384 (TM-score = 0.88, ipTM = 0.88), and PHROG 292 (TM-score = 0.63, ipTM = 0.77) (Fig. 1C, Table S2). Confirmed monomers typically showed low ipTM values (0.12–0.33, Fig. 1B). We hypothesise that exceptions in these distributions arise from interaction malleability (such as dynamic, redox-dependent oligomerisation), contextual oligomerisation, or method-dependent biases that result in capturing monomeric forms during crystallisation. For example, the *Saccharomyces cerevisiae* riboside hydrolase (PDB 8rih), annotated as a structural homologue of PHROG 8702 (unknown function), was deposited in the PDB as a monomer but achieved a high ipTM score (0.86). This protein forms disulphide-linked dimers under oxidising conditions^37^, suggesting the dimer predicted by the ipTM score is correct.

### Prediction of phage protein homo-oligomeric states with AlphaFold-Multimer

Having established that interface quality metrics can reliably distinguish biologically relevant interfaces from false positives, we next developed a systematic framework to predict homooligomeric state. Because homooligomeric proteins are sometimes misannotated as monomers in the PDB when their larger biological assemblies are not explicitly defined, we manually curated all monomeric entries to exclude cases where the monomeric state lacked corroboration from the literature or conflicted with the known oligomeric state of close homologues (Table S3). Combining these 36 validated monomers with established homooligomeric PDB entries, our final benchmark comprised 196 homodimers, 25 homotrimers, 34 homotetramers, 32 homopentamer–hexamer assemblies, and 20 complexes with seven or more subunits.

AlphaFold-Multimer can predict homooligomeric states by systematically modelling proteins across different subunit configurations^30,32,33,39^. We thus tested whether ipTM scores were maximised when proteins were modelled in their correct homooligomeric form. For example, PHROG 807 (’major tail protein with Ig-like domain’), which has high sequence homology to the hexameric PDB entry 8rk8 (0.96 identity, E value=2.26 × 10^−157^), achieved its highest ipTM (0.75) when modeled as a hexamer (Fig. 2A-B). Based on this principle, we implemented a structure-based prediction framework in which each protein was systematically modelled with 2-10 subunits, with five independent predictions per configuration. To reduce ambiguity between closely related states (e.g., pentamers and hexamers, both plausible for major capsid proteins), stoichiometries were grouped into bins (2-mer, 3-mer, 4-mer, 5–6-mer, and 7+-mer). This binning minimises misclassifications between adjacent states and ensures performance metrics reflect biologically meaningful differences rather than marginal distinctions, for example, 5-mers and 6-mers are grouped because many phage proteins, including capsid proteins, form quasi-equivalent pentameric and hexameric assemblies within the same capsid architecture (e.g. *Bacillus subtilis* dsDNA phage ϕ29)^40^. An ipTM threshold of 0.65 was empirically chosen to balance prediction accuracy (0.894) with dataset coverage (0.603) (Fig. S4A-B).

**Figure 2:**
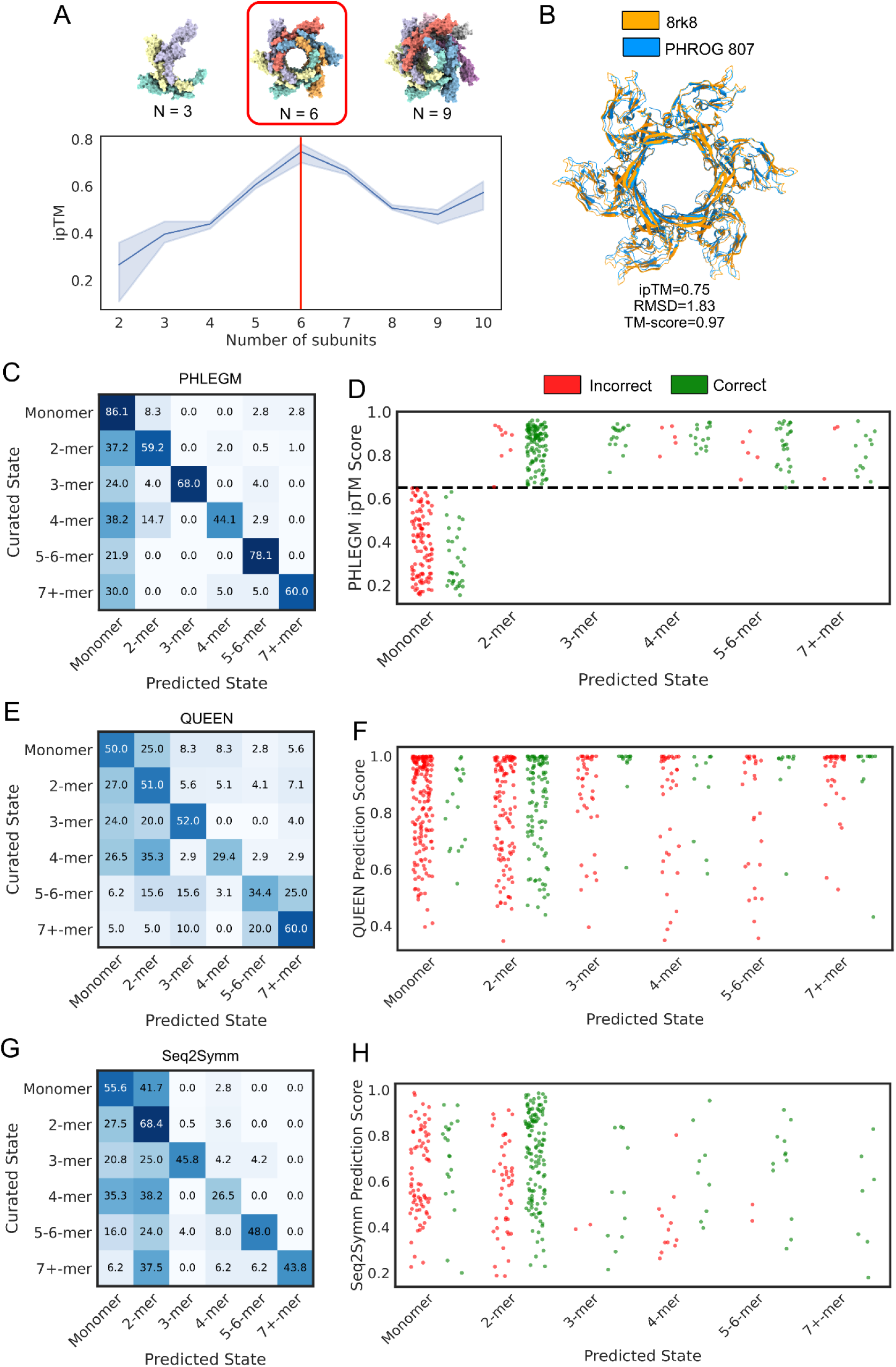
Generating homooligomeric predictions of phage proteins. A. Comparisons of the ipTM score across different stoichiometries for PHROG 807 “major tail protein with Ig-like domain”. The shaded region indicates the standard deviation across five replicates. Structural visualisations are shown for structures with stoichiometries of N=3,6,9. B. Structural alignment between the hexameric AlphaFold-Multimer structure of PHROG 807 and the experimentally determined structure of a sequence homolog (PDB 8rk8). C. Confusion matrix for PHLEGM method showing classification performance across all homooligomeric states presented as row-normalised percentages (true class). D. Stripplot of PHLEGM ipTM scores for each predicted state, demonstrating how prediction confidence varies across homooligomeric classes and correlates with prediction accuracy. The dashed horizontal line at 0.65 indicates the optimal confidence threshold. Note that PHLEGM does not explicitly predict monomers because ipTM requires at least two chains. The ‘Monomer’ column thus represents cases where no multimeric model exceeded ipTM = 0.65, and the plotted values represent the maximum ipTM achieved across the evaluated multimeric states. E. Confusion matrix for the QUEEN method showing predicted versus curated homooligomeric states, presented as row-normalised percentages (true class). F. Stripplot showing QUEEN probability score distributions for each predicted homooligomeric state, with points coloured by prediction correctness. G. Confusion matrix for the Seq2Symm method displaying prediction performance across homooligomeric states, presented as row-normalised percentages (true class). H. Stripplot of Seq2Symm confidence scores for each predicted state, showing the distribution of scores for correct versus incorrect predictions.

This framework, termed Phage Homomer Level Estimate Generation Method (PHLEGM), achieved strong performance for dimers (F1 = 0.72), trimers (0.81), and 5–6-mers (0.81), and moderate accuracy for tetramers (0.56) and 7+-mers (0.69). Tetramers were often misclassified as dimers (14.7%), likely reflecting structural similarity between dimer-of-dimer and tetrameric assemblies. PHLEGM does not explicitly predict monomers because ipTM quantifies inter-chain quality; accordingly, monomers are inferred when no multimeric model exceeds ipTM=0.65, achieving weaker performance (F1=0.36) (Fig. 2C-D).

We benchmarked PHLEGM against two language-model-based approaches: QUEEN^34^ and Seq2Symm^35^, which both utilise ESM2 embeddings^39^. While Seq2Symm correctly classified more true dimers than PHLEGM (68.1% vs 53.2%), this resulted from systematically overpredicting the dimeric state at the expense of 4-mers and 7+-mers (Fig. 2G). Furthermore, both QUEEN and Seq2Symm frequently assigned high confidence to incorrect predictions across all classes (Fig. 2E-H). PHLEGM demonstrated superior confidence calibration (Fig. 2D), ultimately achieving the highest overall performance (QUEEN: accuracy 0.48, F1-macro 0.41; Seq2Symm: accuracy 0.58, F1-macro 0.53; PHLEGM: accuracy 0.63, F1-macro 0.66) (Fig. S5).

To assess generalisability, we evaluated PHLEGM on the validation set of complexes deposited after the cutoff date for AlphaFold-Multimer training, thereby testing on unseen folds and interfaces. Of 39 proteins, 20 were predicted with high confidence (ipTM > 0.65), and 12 were assigned their exact homooligomeric states rather than only the correct bin. In 10 of 11 cases exceeding the ipTM threshold, predicted models closely matched experimental structures, with several aligning to their experimentally derived structures with TM-score above 0.9 and low Root Mean Square Deviation (RMSD), suggesting near-atomic accuracy (Table S4).

### Multistep experimental validation of homooligomeric state predictions

Next, we set out to experimentally validate the computational predictions for nine selected bacteriophage proteins with predicted homooligomeric states ranging from monomers to hexamers (Fig. 3A, Table S5). Hereafter, we refer to the monomer as Px1a, dimers as Px2a-c, trimers as Px3a-c, pentamers as Px5a-b, and a hexamer as Px6a. Proteins were evaluated through a nine-step experimental workflow from gene synthesis to protein characterization (Fig. 3B). First, each of the corresponding synthetic DNA fragments of the nine phage genes were ordered (Integrated DNA Technology, Step I). To introduce restriction sites, we amplified the genes by PCR (Step II) and cloned them in a bacterial expression vector using restriction/ligation cloning (Step IV), allowing for the heterologous production of the respective gene product in *Escherichia coli*. Nine genes were amplified by PCR and produced enough DNA material for cloning. To facilitate single-step purification of the protein after production, we generated a C-terminal streptavidin tag translational fusion to each gene by cloning the streptavidin tag coding sequence in-frame at the 3′ end of the gene sequence. C-terminal tagging was selected based on structural predictions showing it to maximise the tag’s surface accessibility while minimizing interference with native folding and oligomerization. Although computationally predicting the effects of the tag on the protein solubility remains unreliable^41^, structural predictions suggested that the C-termini are exposed and flexible in all selected proteins (Fig. S6). Additionally, we found that the tag did not affect the protein folding or oligomerization.

**Figure 3:**
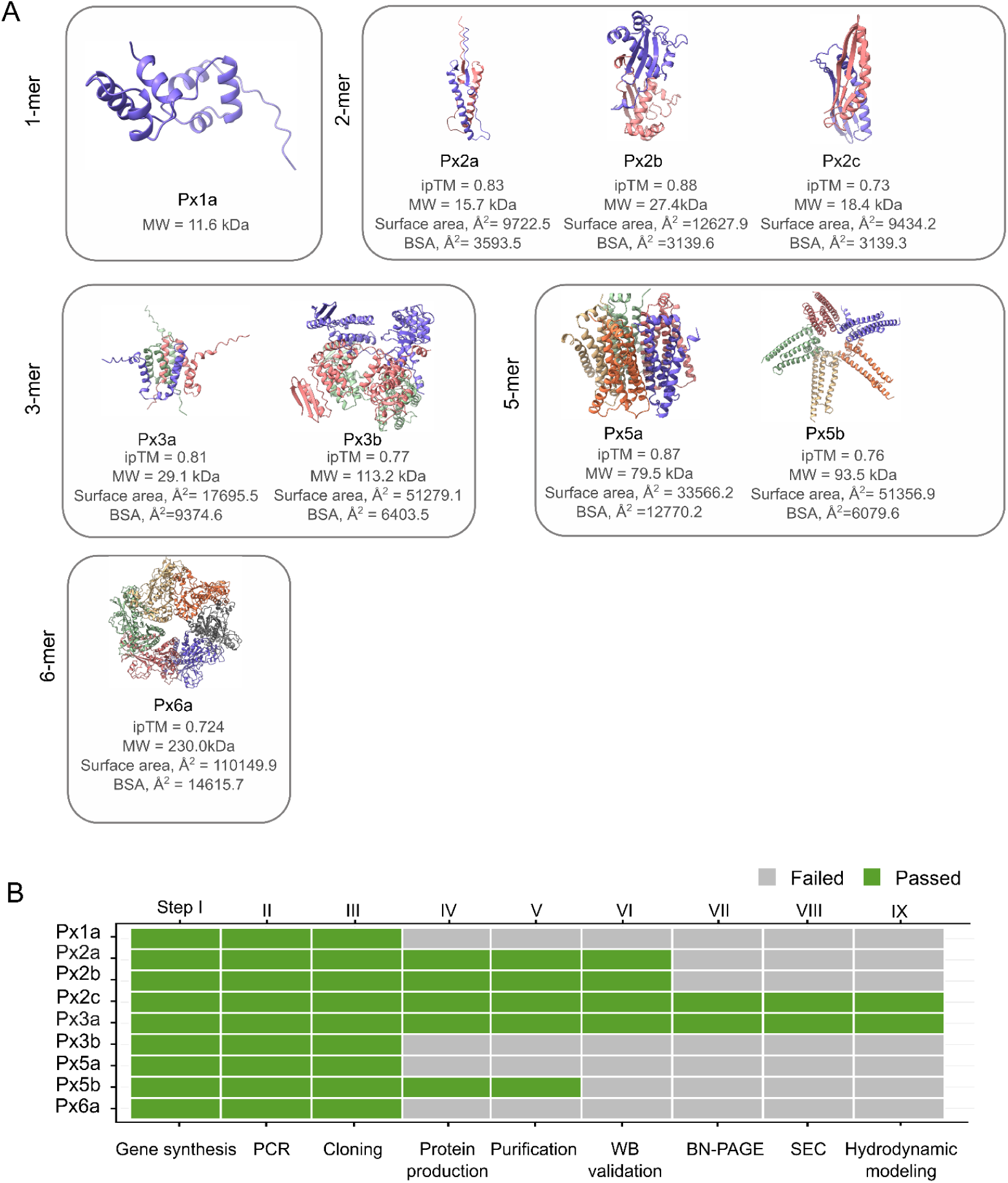
Multi-step screening of experimental candidates with progressive dropout. A. Predicted multimeric structures of nine candidate proteins. Predictions showing homooligomeric assemblies ranging from monomers (1-mer) to hexamers (6-mer) with the molecular weight (M_W_) of 11.6-290.3 kDa. Interface predicted template modeling scores (ipTM) are shown for multimeric structures. For each candidate, the total surface area and predicted buried surface area (BSA) are reported as measures of predicted complex stability. Structures are coloured by chain. Detailed information about the proteins and their function is summarised in Supplementary Table S5. B. Dropout pattern across sequential screening steps. A total of nine candidate proteins were selected for the experimental workflow. Only Px2c and Px3a completed all steps, including size-exclusion chromatography (SEC) and hydrodynamic characterization. WB: Western blot; BN-PAGE: Blue Native polyacrylamide gel electrophoresis.

In Step III, gene expression and protein production of the streptavidin-tagged constructs in *E. coli* were assessed by denaturing sodium dodecyl sulfate polyacrylamide gel electrophoresis (SDS-PAGE) of cell lysates. The successful overproduction of Px2a, Px2b, Px2c, Px3a, and Px5b was apparent from the presence of an additional band in the Coomassie-stained gel at the position of the respective protein monomer (Fig. S7). Three constructs (Px1a, Px3b, and Px6a) showed no detectable expression in the cell lysates, possibly due to the formation of high molecular weight protein aggregates, inclusion bodies that resulted from improper folding of the proteins, or the dependence on phage-specific chaperones that were absent in the heterologous host^42^. One construct (Px5a) appeared to be toxic to the *E. coli* host, as it prevented growth and thus adequate expression of the protein. For purification of the heterologously produced phage proteins, all eight *E. coli* cell lysates were applied to Strep-Tactin affinity chromatography (Step V). Next, the eluates were analysed by SDS-PAGE and Western blot (Step VI). The visualization of the fused *Strep*tag in the Western blot was performed with the Strep-Tactin AP conjugate (Fig. S8A-B). The results of the purification attempt confirmed the observation made with the cell lysates above, as only Px2a, Px2b, Px2c, Px3a, and Px5b were successfully purified. An aberrant protein size band was observed for protein Px5b in the eluate (Fig. S8A-B). We hypothesise that proteolytic degradation could cause Px5b to migrate at around 15 kDa, i.e. smaller than the calculated molecular weight of 20 kDa.

To investigate protein complex formation under native conditions, all eluates from the protein purification step were analysed by Blue Native PAGE (Step VII). While Px3a (Ip = 6.61, calculated using PropKa via PDB2PQR) migrated as a single band, Px2c (Ip = 5.96) displayed two bands. The discrete bands were consistent with stable homooligomeric complexes (Fig. S9). We did not observe band migration for Px2a and Px2b. For protein Px2a, the predicted isoelectric point of the dimer (pI = 9.1) is higher than the BN-PAGE running pH of 7.0, giving the protein a net positive charge under these conditions and likely preventing its migration. In contrast, Px2b has a predicted dimer pI of 3.82, well below the running pH, suggesting that surface charge is unlikely to explain the lack of movement. We therefore hypothesise that Px2b may form aggregates under native conditions, impeding its migration.

Px2c and Px3a were further analysed by size exclusion chromatography (SEC, Step VIII), calibrated with globular protein standards (Fig. S10). Both proteins displayed a single symmetric peak in the SEC chromatogram, representing the majority of the protein fraction. SEC separates proteins according to the hydrodynamic volume, which reflects not only molecular mass but also shape and solvation. A protein’s elution behaviour is governed by its Stokes radius (Rs), the effective hydrodynamic radius determining diffusion through column pores. Rs is a hydrodynamic value indicative of the apparent size of the dynamic hydrated protein^43^. Globular proteins exhibit Stokes radii close to their geometric radii, whereas elongated or asymmetric proteins have larger Stokes radii relative to their mass. Consequently, asymmetric protein complexes elute earlier than spherical proteins of equivalent molecular weight ^44^. Px3a eluted at 10.85 mL (Kav = 0.156), corresponding to an apparent molecular weight of approximately 51.6 kDa based on the calibration curve, i.e. larger than the calculated trimeric mass of 32.2 kDa. Px2c eluted at 11.80 mL (Kav = 0.217), corresponding to an apparent molecular weight of approximately 36.4 kDa (Fig. 4A, B).

**Figure 4:**
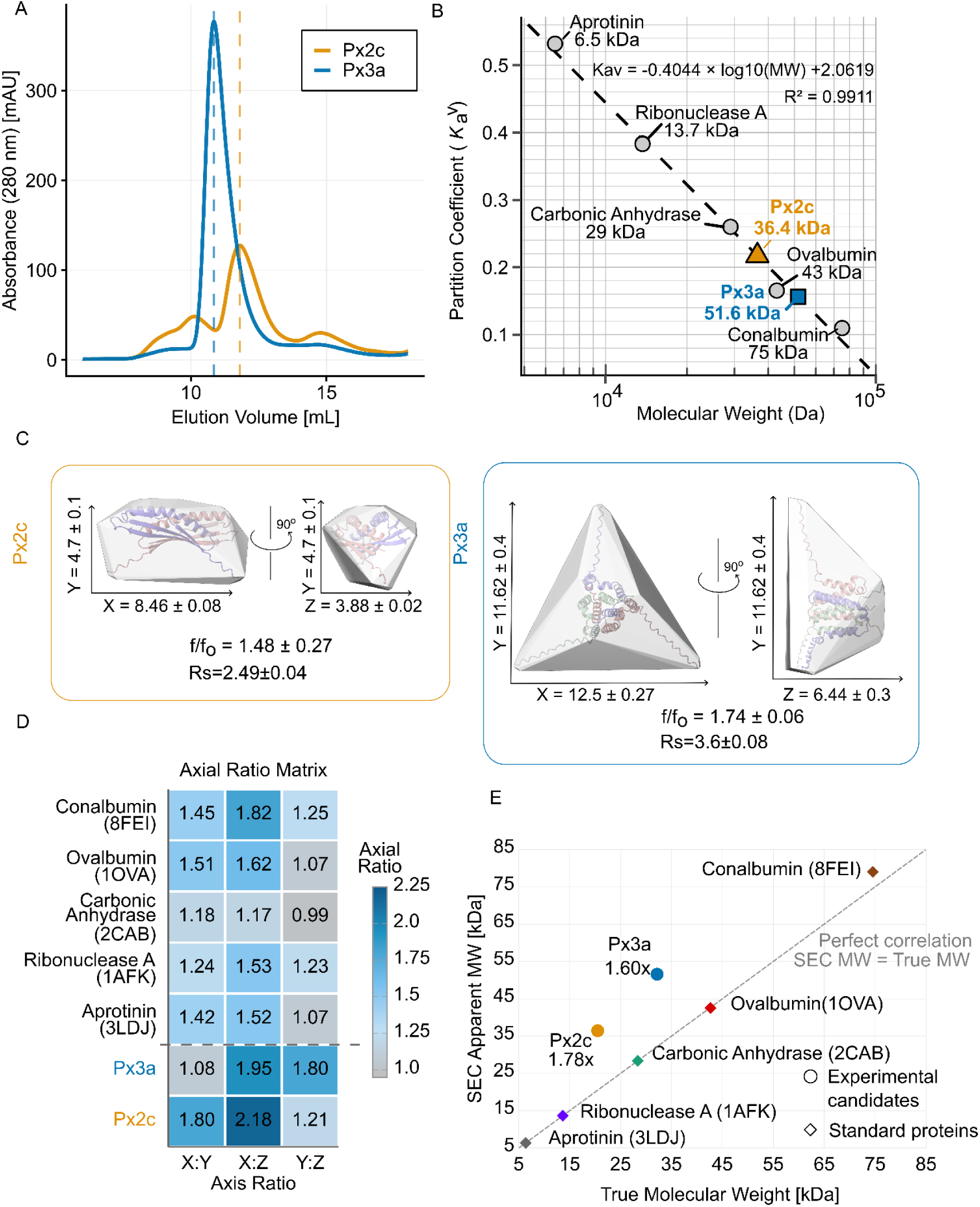
Size-exclusion chromatography (SEC) analysis of apparent molecular weights. Geometric architecture distinguishes Px2c and Px3a chromatographic behavior. A. SEC elution profiles for Px2c and Px3a are monitored at 280 nm. Vertical dashed lines indicate elution volumes used to determine apparent molecular weights from the calibration curve. B. SEC calibration curve using Superdex 75 Increase 10/300 GL column with protein standards: aprotinin (6.5kDa), ribonuclease A (13.7kDa), carbonic anhydrase (29kDa), ovalbumin (43kDa), and conalbumin (75kDa). C. Protein oligomer structures with convex hull representations, HullRad and SOMO calculated dimensions, hydrodynamic radius (Rs) and frictional ratios (f/f₀). Px2c as a dimer exhibits prolate (rugby ball) geometry with pronounced elongation along the X-axis (f/f₀ = 1.49±0.27, Rs = 2.49±0.04 nm, 8.46 × 4.7 × 3.88 nm). Px3a trimer shows flattening oblate-shaped (disc-like) geometry with near-equal X and Y dimensions but compressed Z-axis (f/f₀ = 1.74±0.06, Rs = 3.6±0.08 nm, 12.5 × 11.62 × 6.44 nm). D. Axial ratio matrix heatmap showing the degree of elongation and compression relative to spherical geometry. Ratios equal to 1 indicate isotropic dimensions and spherical geometry, ratios >1 indicate elongation and prolate structure, and ratios <1 indicate compression and oblate structure. Px2c exhibits consistent elongation across all ratios, characteristic of prolate geometry ^45^, while Px3a displays selective compression with high X:Z and Y:Z ratios but near-unity X:Y ratio, characteristic of oblate geometry. E. SEC apparent molecular weight versus true molecular weight. Reference proteins fall on or near the identity line, while our candidate oligomers Px2c and Px3a exhibit SEC apparent molecular weight overestimations of 1.78-fold and 1.60-fold, respectively, in line with their calculated dimer and trimer molecular weights.

To resolve the discrepancy between calculated and SEC-derived molecular weights, we calculated hydrodynamic parameters. Hydrodynamic parameters are used to gain information on the structure of proteins and their assemblies. Such parameters can be computed from “dry” protein structures by performing hydrodynamic modelling (HM) (Step IX). We calculated hydrodynamic parameters of our predicted multimeric structures with two computational methods with different underlying algorithms. The UltraScan SOlution MOdeler (US-SOMO) is a suite of computer programs centred on hydrodynamic modelling and simulation. US-SOMO integrates four different bead modelling tools. A bead is the simplest geometrical body - a sphere of defined radii, that collectively produces detailed representations of biomolecules^46–48^. Additionally, HullRad was employed as an independent method for convex hull analysis (Figure 4C). HullRad uses a convex hull to model the hydrodynamic volume of the protein. The convex hull is mathematically defined as the smallest convex envelope that contains a set of points. The convex hull of a three-dimensional molecular model encapsulates both the overall volume of the molecule as well as its shape asymmetry and surface roughness. These are characteristics that influence hydrodynamic diffusion ^49^. Both tools were applied to Px2c and Px3a, alongside all reference proteins with well-characterised shapes. First, we ran Monte Carlo simulations with 10⁶ random walks at pH 8.0 and 8°C, which provided hydrodynamic parameters for both our oligomers under the experimental conditions of the SEC.

Analysis of the Px2c dimer (True M_W_ 20.48 kDa) based on SOMO bead model analysis with ZENO surface-based calculations revealed a hydrodynamic radius (R_s_) of 2.49±0.04 nm and a frictional ratio (f/f₀) of 1.49±0.27, indicating that the dimer has a moderately asymmetric geometry. ZENO surface-based calculations also showed the radius of gyration (Rg = 2.24 nm), which evaluates the compactness or overall size of a protein molecule. Rg quantifies the distribution of mass around the center of mass of the protein and provides insights into its structural properties and dynamics. The elongated architecture, measuring 8.46 × 4.7 × 3.88 nm with axial ratios of 1.80 (X:Y), 2.18 (X:Z) and 1.21(Y:Z), suggests an elongated prolate ellipsoid. The dimer eluted at 11.80 mL with an apparent molecular weight of 36.4 kDa, a 1.78-fold overestimation of its true mass. This rate of overestimation is explained by the volume the protein occupies in solution and is calculated also by SOMO as the hydrated molecular volume from *vbar* (Partial specific volumes) and equals 36 kDa.

Px3a formed a trimer in solution with hydrodynamic properties indicative of a highly asymmetric, oblate architecture (Fig 4C). Bead-model analysis of the Px3a trimer (32.2 kDa; 364 beads) yielded a hydrodynamic radius (R_s_) of 3.6±0.08 nm. Two independent measures of the frictional ratio (f/f₀) yielded 1.74±0.06, indicating good agreement between methods. The radius of gyration of Px3a trimer is 3.05 nm. Shape modelling estimated dimensions of 12.5 × 11.62 × 6.44 nm, with nearly identical X and Y axes (axial ratio X:Y = 1.08) and a larger X:Z ratio of 1.95, confirming a flattening oblate geometry. Consistent with these hydrodynamic properties, size-exclusion chromatography showed that the Px3a trimer eluted earlier at 10.85 mL with an apparent molecular mass of 51.6 kDa, representing a 1.60-fold overestimation relative to its true mass. This rate of overestimation is explained by SOMO hydrated molecular volume calculations from vbar and equals 56 kDa. Px2c and Px3a displayed markedly distinct molecular geometries confirmed by both analytical methods. The close agreement between HullRad and SOMO-based frictional ratios (Px2c: 1.49±0.27; Px3a: 1.74±0.06) validates both computational approaches and confirms the reliability of our size-shape determinations.

Our results suggest that the apparent M_W_ discrepancies stem from the non-spherical protein shapes rather than experimental artifacts or sample heterogeneity. The M_W_ overestimation factors for Px2c and Px3a (1.78-fold and 1.60-fold, respectively) correlate well with the frictional ratios (1.49±0.27; 1.74±0.06, respectively), consistent with SEC apparent masses reflecting the effective hydrodynamic volumes of elongated homooligomeric protein complexes. These results indicate that the two homooligomers adopt different stable geometries in solution, a prolate dimer and an oblate trimer, and that their shape-dependent chromatographic behaviour can be explained by boundary element hydrodynamic modelling ^50,46^. This pattern demonstrates that SEC matrices are sensitive to the specific geometric distribution of mass prolate versus oblate architecture in ways that complement but extend beyond solution-phase frictional measurements. The distinction between Px2c prolate and Px3a oblate geometries manifests in their differential SEC behaviour, providing experimental evidence that geometry is a critical parameter for chromatographic characterization alongside traditional hydrodynamic metrics.

Although only two out of nine proteins successfully completed the 9-step experimental pipeline, the results obtained for Px2c and Px3a support their predicted oligomeric states, a dimer and trimer, respectively. Such attrition is common in structural and biophysical characterisation pipelines, and the successful characterisation of even a subset of targets provides meaningful validation of the computational predictions.

### Prediction and structural clustering of homooligomeric states across the phage proteome

We next applied PHLEGM to all 22,554 representative proteins in the PHROG database^36^ to predict their most probable homooligomeric states. Because our earlier evaluations demonstrated that PHLEGM is unreliable for predicting monomeric configurations, monomers were excluded from this analysis. Homooligomeric states were confidently predicted for 6,902 proteins (30.6%, ipTM > 0.65). The remainder either failed to exceed the threshold for any stoichiometry or showed comparable scores across multiple states, precluding unambiguous prediction.

Functional annotations were available for 1,125 of the confidently predicted representatives (16.3%), revealing distinct enrichment patterns across PHROG categories (Fig. S11A, Fig. S12). Tail fibre proteins were generally trimeric (31/38; Fig. 5A), and terminase small subunits were consistently classified as 7+-mers (37/38), in agreement with their established nonameric organisation (9-mers, Figure 5B)^51,52^. Dimers predominated among nucleic acid metabolism, transcription regulation, and morons (Fig. S11A). Among the 5,777 PHROGs lacking functional annotation, 3,567 were predicted as dimers. Because dimeric assemblies were strongly associated with regulatory and metabolic roles in our annotated set, we infer that these largely correspond to non-structural proteins (Fig. S11A).

**Figure 5:**
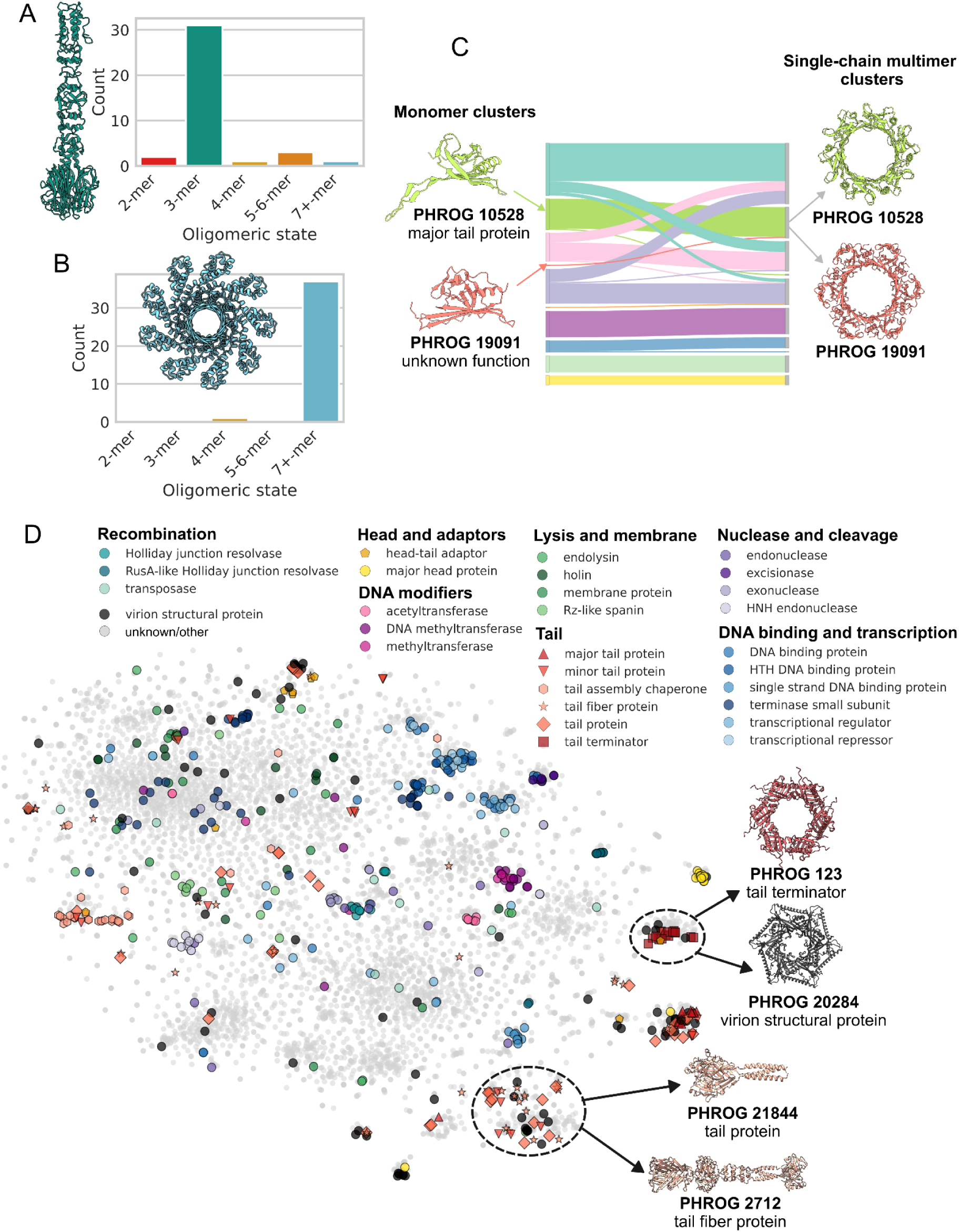
Predicted homooligomeric states and structural clustering of PHROGs. A. Predicted homooligomeric states of PHROGs annotated as ‘tail fiber protein’ (n=38). Representative shown of PHROG 24210 folded as a 3-mer. B. Predicted homooligomeric states of PHROGs annotated as ‘terminase small subunit’ (n=38). Representative shown of PHROG 308 folded as a 9-mer. C. Correspondence between Foldseek monomer-based and multimer-based clustering. The Sankey diagram shows the mapping of the eight largest clusters predicted from monomeric structures (left) and single-chain multimeric assemblies (right). Selected clusters are illustrated with AlphaFold-Multimer models, including a major tail protein (PHROG 10528) and a previously uncharacterised protein (PHROG 19091). D. t-SNE projection of multimeric PHROG representative structures coloured by functional annotation. The 30 most frequently occurring annotations are shown and all others grouped as ‘other’/unknown.

To assess whether homooligomeric state information improves functional annotation, the 6,902 confidently predicted assemblies were clustered with Foldseek using either monomeric structures or “single-chain multimer” representations in which subunit coordinates were concatenated into a single chain. This concatenation bypassed the overly strict constraints imposed by standard multi-chain clustering with Foldseek-Multimer which resulted in 96% (6,248/6,902) of assemblies being classified as singletons. By collapsing the subunits into a single chain, Foldseek performed broader shape-based comparisons reducing the singleton rate to 66% (4,542/6,902; Table S7; Fig. S11A-B). Comparing monomeric and single-chain multimer clustering produced 723 and 697 non-singleton clusters, respectively, with moderate agreement (adjusted Rand index = 0.507; Fig. S11C, Table S7). While 227 clusters grouped the same sets of proteins regardless of whether monomeric or multimeric representations were used (Fig. S11A-B), systematic differences emerged. For example, proteins that clustered together as monomers were sometimes split when quaternary context was introduced, reflecting interface-dependent rearrangements. Conversely, some multimers formed coherent clusters despite their monomers not clustering, indicating that homooligomeric state higher-order assembly can reveal relationships obscured at the monomer level. For instance, PHROG 19091, unannotated and a singleton in the monomeric analysis, formed a hexameric ring that clustered with PHROG 10528, a major tail protein, in the multimeric analysis, suggesting a comparable structural role (Fig. 5C).

Having grouped the 6,902 multimeric assemblies into discrete clusters, we next explored the broader structural relationships among these complexes. We derived symmetric pairwise distances from the Foldseek-Multimer template modelling scores (see Methods) and visualised these structural similarities using t-distributed stochastic neighbour embedding (t-SNE) (Fig. 5D). Complexes annotated as DNA methyltransferases, HNH endonucleases, and tail assembly chaperones formed distinct clusters, likely reflecting stringent functional constraints. For example, DNA-modifying enzymes frequently assemble into homodimeric or high-order even numbered oligomers to provide the two-fold symmetry required to position active sites on opposite strands of the target DNA^53,54^, while tail assembly chaperones must assemble in defined stoichiometries to scaffold virion morphogenesis^55^. The structural organisation visible in the t-SNE map facilitates refinement of phage protein annotations.

Proteins annotated broadly as “virion structural protein” frequently shared structural space with functionally characterised proteins, such as tail terminator, major tail, and tail fibre proteins. For example, hexameric PHROG 20284, annotated as “virion structural protein,” showed structural similarity to tail terminator proteins, suggesting analogous function. Likewise, PHROG 21844 (“tail protein“; from *Staphylococcus* and *Lactococcus* phages) similarly resembled tail fibre proteins, although its architecture appeared to be more compact than the canonical tail fibre protein PHROG 2712 (from *Pseudomonas* phages).

Beyond refining broad functional descriptions, this assembly-level perspective provides a novel pathway for characterising completely unannotated proteins. While standard monomeric comparisons only evaluate individual chains, clustering based on assembled structures allowed us to evaluate proteins that were completely orphaned at the monomer level, meaning they failed to group with any known proteins and remained as isolated singletons or clustered exclusively with other uncharacterised sequences. This approach successfully rescued 37 previously unannotated PHROGs, mapping them into characterised multimer clusters and assigning putative roles, such as virion structural components and tail proteins (Fig. S11E, Supplementary Table S8).

## Discussion

Determining the homooligomeric states of phage proteins is essential for understanding their roles in genome packaging, capsid stabilisation, host recognition, and cell wall penetration. Yet, the homooligomeric states of most phage proteins remain unknown, owing to the scarcity of experimentally resolved multimeric structures and the lack of functional annotation for many viral genes. Here, we present the first benchmark to date for predicting homooligomeric states in phages, using AlphaFold-Multimer to model thousands of structurally representative sequences from the widely used PHROGs annotation system^36^.

Our results demonstrate that AlphaFold’s neural network has implicitly learned features associated with homooligomerisation, allowing it to reconstruct biologically relevant quaternary structures even without experimental templates. The ipTM score serves as an indicator of interface validity, with higher values consistently associated with well-packed, chemically plausible assemblies^56^. While AlphaFold-Multimer does not recover all biologically relevant interfaces, resulting in false negatives, our benchmarking confirms that its predicted interfaces are generally high-quality and accurately represent known functional homooligomers. Indeed, the subset of low-scoring experimental dimers in our set may reflect crystal packing artifacts rather than true biological interactions, while high-scoring monomers could represent true biological dimers that were misannotated or forced into monomeric states by specific experimental conditions. Consequently, in instances where historical experimental structures conflict with high-confidence multimeric predictions, our models provide highly plausible hypotheses for the native biological state that warrant further experimental re-evaluation.

Protein language models have advanced phage protein function prediction^57–61^, yet their utility for quaternary structure inference remains limited. We showed that predictions using ESM2 embeddings^39^ with QUEEN^37^ did not reliably resolve homooligomeric states (Fig. 2). Fine-tuning ESM2 embeddings, as in Seq2Symm^35^, through a modified language model head improves performance but remains insufficient for accurate prediction on our curated reference set of phage proteins with known oligomeric states (Fig. 2). This may reflect the unique compositional biases and limited homology of phage proteins compared to the predominantly cellular proteins used in language model training. Although these two language model-based approaches, QUEEN and Seq2Symm, are far more computationally efficient than PHLEGM, complete AlphaFold-Multimer modelling remains necessary for robust homooligomer prediction. Specifically, our evaluation reveals that a non-monomeric prediction by PHLEGM is highly trustworthy offering a reliable indicator of true higher-order assembly. While PHLEGM’s monomer predictions warrant more cautious interpretation, acting as a default category for challenging assemblies, this still represents a significant advantage over current language-model approaches, which remain unreliable across all oligomeric states. To enable researchers to bypass this computational expense, we provide a complete set of AlphaFold-Multimer folds for PHROG representatives, including the predicted multimeric models, as a publicly available resource (https://linsalrob.github.io/PHROG_structures). Future improvements to quaternary state prediction may leverage the architectural advances of next-generation models like AlphaFold3^23^ or its open-source counterparts, such as Boltz^62,63^. Alternatively, performance may be boosted by adopting different tokenisation strategies for protein language models or by leveraging AlphaFold2’s internal representations, rather than its final structural outputs, which were recently used to distinguish between dimeric, trimeric, and tetrameric coiled-coil domains^32^.

To assess the reliability of PHLEGM predictions in practice, we experimentally validated homooligomeric states for a subset of nine phage proteins. Experiments are important to benchmark computational predictions, establish confidence in large-scale structure prediction pipelines, and verify that predicted assemblies represent biologically relevant states. However, experimental validation proved challenging: five out of nine candidate proteins could be expressed and further purified for multimeric characterisation . The remaining four failed at earlier stages of the 9-step pipeline, possibly due to the formation of inclusion bodies, cytotoxicity, or dependence on phage-specific chaperones that were absent in the expression host. Of the five successfully purified proteins, only Px3a and Px2c produced stable homooligomeric complexes amenable to chromatographic analysis, while one was lost due to possible proteolysis, and two formed high-molecular-weight aggregates incompatible with native gel separation.

Native gel electrophoresis, the standard method for multimer characterization, requires proteins to maintain their assembled state during hours-long electrophoresis. This may be complicated by elongated geometries, which migrate aberrantly due to shape-dependent sieving effects. Shape effects similarly impacted SEC analyses. Since SEC separates proteins by hydrodynamic volume rather than mass, elongated molecules elute earlier than compact ones of equal weight, artificially inflating apparent molecular weights by 1.78-fold for Px2c and 1.60-fold for Px3a. To understand these shape effects, we used two independent computational methods: HullRad convex hull analysis and SOMO bead modelling with ZENO Monte Carlo calculations. The methods agreed remarkably well. This convergence validates both the structural models and our geometric assignments, revealing that Px2c had a prolate (rugby ball-like) geometry, while Px3a has an oblate (disc-like) geometry with near-equal X and Y dimensions but compressed Z-axis Despite Px2c having lower frictional ratios than Px3a (f/f₀ = 1.49±0.27 vs 1.74±0.06) indicating less hydrodynamic resistance, it displayed greater SEC overestimation (1.78-fold vs. 1.60-fold). This demonstrates that SEC matrices interact differently with rods and discs, a distinction lost in spherically-averaged parameters like f/f₀. While characterization methods commonly assume that proteins are roughly spherical, many proteins and protein complexes are rod- or disc-shaped, or have extended structures. SEC elution profiles deviating from expected behaviour based on molecular weight commonly reflect the shape of the protein. Elongated prolate proteins present a larger hydrodynamic cross-section relative to their molecular weight, causing earlier elution and systematic overestimation of apparent molecular weight when calibrated against globular standards. Oblate proteins, by contrast, are flattened perpendicular to their longest axis, reducing their effective hydrodynamic radius relative to their mass and producing a distinct deviating elution behaviour. The degree of deviation from expected elution is therefore not simply a function of molecular weight or oligomeric state, but is directly influenced by the three-dimensional shape of the hydrated protein volume and how it interacts with the porous polymer network of the SEC matrix. Dimensionality analysis based on structural modelling can predict which proteins will exhibit shape dependent deviation behavior and account for these effects.

Our clustering analyses underscore the value of incorporating quaternary structure into large-scale comparative studies. While some proteins cluster consistently in their monomeric and multimeric forms, others diverge when their homooligomeric state is taken into account, or converge despite distinct monomeric folds. We hypothesise that these shifts reflect functional specialisation shaped by interface geometry, symmetry, and higher-order assembly, revealing relationships that remain hidden at the monomer level^30^. Structural clustering has been successfully applied to transfer annotations between proteins with similar monomeric folds^30,64,65^. Given that clustering outcomes appear to be influenced by homooligomeric state, these algorithms should be enhanced to consider quaternary structure explicitly. Notably, our analysis revealed that nucleic acid metabolism proteins are overwhelmingly dimeric, potentially allowing DNA to engage symmetrically with both subunits, a configuration that may facilitate progressivity or cooperative binding. Such functional constraints likely drive the strong preference for specific oligomeric states within protein families. Multimer-informed clustering, therefore, represents a powerful strategy for function annotation, grouping proteins by assembly structure even when their monomers differ, and helping resolve ambiguous or uncharacterised cases.

Predicted homooligomeric states provide a foundation for computationally reconstructing phage components and, ultimately, whole virions. Application of the PHELGM procedure to tail fibre proteins from *Enterococcus* phages containing depolymerase domains demonstrates this potential, yielding multimeric structures that clarified their structural organisation and putative role in host interaction^66^. Extending this strategy to capsid subunits, baseplates, and tail tubes could contribute to predicting virion organisation *in silico*, with potential applications in synthetic biology. For example, computational models could guide modular phage redesign, including the retargeting of host specificity through engineered receptor-binding proteins^67^. As prediction accuracy continues to improve, computational assembly may ultimately enable the design and construction of functional phages entirely *in silico*.

## Conclusion

This study presents the largest systematic framework to date for predicting and classifying phage protein homooligomers directly from sequence, combining AlphaFold-Multimer modelling, interface-based scoring, and multimer-informed clustering. Applied to more than 25,000 representative proteins, our approach revealed extensive diversity in homooligomeric states and identified clear structural patterns associated with essential viral functions.

Compared with recent protein language model–based methods, this framework demonstrated superior accuracy in predicting oligomeric states for proteins from the PHROGs database. To experimentally validate these computational predictions, we developed and applied a nine step biochemical workflow to nine selected phage proteins spanning a range of predicted oligomeric states. Two proteins, Px2c and Px3a completed the full pipeline, and their oligomeric states were confirmed by size-exclusion chromatography and hydrodynamic modelling. The differences in their chromatographic behaviour were explained by their molecular geometries. Beyond providing a resource for experimental validation, these predictions lay the groundwork for computationally reconstructing viral architecture and for uncovering the structural principles that govern phage protein function.

## Methods

### Data preparation

#### PHROG representative selection

To generate representative multimeric protein structures for each PHROG^36^, we first identified a representative monomer within each PHROG. All non-redundant PHROG proteins (440,050 sequences) were predicted in their monomeric form using ColabFold (version v1.5.5) with default parameters^68^. Within each PHROG (38,880), we performed all-versus-all structural alignments using Foldseek (version 6cfb8805be28925a81f194e4b616ad5616f2ed9a)^24^ with the “--exhaustive_search” flag to quantify the structural similarity among members. The most representative protein, defined as the protein within each PHROG with the highest cumulative alignment bitscore to all other PHROG group members, was selected as the representative and was made publicly available via the Phage Protein Structure Gallery https://linsalrob.github.io/PHROG_structures. This approach ensured that the chosen monomer captured the structural core of the homologous group, providing a reliable basis for downstream multimeric modelling. The quality of the multiple-sequence alignments (MSAs) used to generate protein structures was evaluated using Neffy (version 0.1.1)^69^ and the number of sequences and the number of effective sequences in each alignment.

To ensure computational efficiency, a sequence length cutoff of 350 amino acids was applied to representative PHROGs, resulting in 35,083 PHROG representatives. This threshold reflects the practical limitations of AlphaFold-Multimer, which exhibits reduced performance and increased memory demands when modelling longer sequences, particularly for multimers.

#### Establishing ground truth homooligomeric states

We annotated the PHROG representative sequences with proteins deposited to the Worldwide Protein Data Bank (wwPDB)^70^ to determine which PHROGs have experimental structures in the wwPDB and whether they appeared in the AlphaFold-Multimer training dataset (deposited before 30 September 2021). The amino acid sequences of all protein structures available in the wwPDB (925,682 sequences, accessed 16 March 2025) were downloaded via the public FTP repository. Sequence similarity searches between PHROG representatives and the PDB protein sequences were performed using MMseqs2 (version 12.113e3) ^71^ with the parameters “--cov-mode 1 --max-seqs 925682 -e 1e-10 --min-seq-id 0.3 -c 0.7”. The “--max-seqs” parameter was set to the total number of PDB sequences to ensure that all potential matches were returned, rather than the default limit of 300. Using this approach, 1,186 PHROG representative sequences were identified as having at least one match to a PDB-deposited protein.

For each matched PDB entry, the deposition date and predicted homooligomeric state were retrieved using the RCSB PDB RESTful API (https://data.rcsb.org) via a custom Python script. Of the 1,186 PHROG representative sequences with matches to PDB entries, 1,088 had at least one associated structure deposited prior to the AlphaFold-Multimer training cutoff date of 30 September 2021, while 98 matched structures were deposited after this date. Among the 1,186 matched representatives, 301 were associated only with monomeric structures in the PDB. However, because proteins may crystallise or be experimentally resolved as monomers even when they form higher-order assemblies in vivo, we manually reviewed the associated literature where available to identify biologically relevant homooligomeric states (Table S2). For non-monomeric homooligomeric complexes, ground truth homooligomeric states were determined as the most frequent homooligomeric state of their PDB hits. PHROG representatives with hits to more than three different homooligomeric states, or with a most common state with a frequency less than 0.65 were excluded from homooligomeric state prediction (122 representatives).

### Homooligomeric state prediction

#### Language Model-based Homooligomeric State Prediction

Predictions of the homooligomeric state were made using Seq2Symm (commit eb83e19)^35^ and QUEEN (commit f96f3b5)^34^.

#### Multimeric structure generation

For each representative PHROG, protein structures were generated using ColabFold (version 1.5.5)^68^. For each sequence, a MSA was generated using colabfold_search, using MMseqs2 (version 15.6f452) against the ColabFold reference database. MSA depth showed a moderate correlation with pLDDT (Pearson’s r = 0.500, Spearman’s r = 0.522). These correlations support the use of a pLDDT threshold to filter for confidently predicted structures: proteins with a mean pLDDT greater than 70 (25,513 structures) were used for homooligomeric state prediction. This threshold is consistent with the criteria used by the AlphaFold Protein Structure Database^72^ to assign ‘high’ and ‘very high’ structures, ensuring that structures are reliably predicted by AlphaFold.

For each representative protein, the original monomeric MSA was replicated and modified to generate nine new MSAs, each corresponding to a different multimeric state ranging from 2 to 10 identical subunits (indicated in the FASTA headers). For each of these modified MSAs, protein complex structures were predicted with colabfold_batch using the alphafold2_multimer_v3 model with 5 structures and 3 recycles without subsequent relaxation. Structures were generated using AMD Instinct MI250X GPUs and 256 GB RAM provided by the Pawsey Supercomputing Research Centre (Perth, Australia).

#### Confidence metric-based evaluation

For each PHROG representative, structural quality metrics were calculated from the corresponding .pdb and .json output files using a custom Python script. We evaluated several confidence and interface-based metrics, including pLDDT^73^, pTM, PAE and pDockQ2 ^74^. These metrics were calculated using code from FoldDock^74^ and biopython (version 1.85). ROC curves and PR curves, ROC-AUC and AP, as well as classification metrics including F1-score, precision, and recall, were computed using scikit-learn (version 1.4.1).

Machine learning approaches were implemented using scikit-learn to compare the performance of individual metrics with that of combined feature sets for monomer/dimer classification. Four models were evaluated using 5-fold stratified cross-validation:

- **Logistic Regression**: A linear classifier with L2 regularisation (max_iter = 1000) used to establish baseline performance.
- **Random Forest**: An ensemble method (n_estimators = 100) capable of ranking feature importance and capturing non-linear relationships.
- **Gradient Boosting**: A sequential ensemble method that improves classification performance through iterative refinement.
- **Support Vector Machine (SVM)**: A kernel-based classifier applied to standardised features to effectively handle high-dimensional data.

Each model was trained on two feature sets: (1) the ipTM score alone, and (2) a combination of metrics including ipTM, pTM, pDockQ2, pLDDT, and interface pLDDT. Model performance was assessed using the area under the ROC curve and average precision.

#### Structural replicates sensitivity analysis

A subsampling analysis was performed to determine the optimal number of structural replicates required for reliable classification. For each protein and for each sample size ranging from one to five structures, the specified number of replicates was randomly selected from the five available models. Both mean and maximum ipTM scores were calculated for the chosen replicates. The area under the ROC curve (AUC-ROC) was then computed for each sample size to evaluate how classification performance scaled with the number of structural replicates.

### Structural evaluation

#### Comparison with experimental structures

We downloaded experimentally derived structures from the PDB, and converted those in .cif format to .pdb format using pdb-tools (version 2.5.0)^75^. Predicted multimeric structures were compared to their corresponding experimental structures using US-align (version 20241201)^76^ to assess structural similarity. Pairwise alignments were performed in multimer mode (-mm 1) with chain-level comparison enabled (-ter 1), and TM-scores were reported using normalisation by the length of the experimental structure (TM2). Root-mean-square deviation (RMSD), sequence identity, and alignment lengths were also extracted. Protein structure visualisations were generated in ChimeraX^77^.

#### Clustering predicted assemblies

Prior to structural search, all structural chains were collapsed into a single chain per model using pdb-tools (version 2.5.0)^75^. Each PDB file was tidied, reassigned to chain “A”, and renumbered from residue 1 to remove gaps or overlaps.

To determine the optimal clustering strategy, the proteins in their predicted homooligomeric state were clustered using FoldSeek Easy-Cluster^24^ on the collapsed single-chain models and FoldSeek Easy-Multimer Cluster^56^ on the full multi-chain assemblies with default parameters. Because direct clustering of full homooligomeric assemblies with Foldseek-Multimer imposed overly high specificity, we adopted the single-chain multimer strategy for our primary analysis.

To evaluate whether homooligomeric state information improves functional annotation over traditional monomeric analysis, the corresponding monomeric structures were also clustered with FoldSeek easy-cluster under default settings. These monomeric and multimeric clustering results were then evaluated against one another using the Adjusted Rand Index (ARI) implemented in scikit-learn, providing a quantitative measure of similarity between clustering outcomes.

### Visualisation

A t-distributed stochastic neighbour embedding (t-SNE) was generated to visualise the structural relationships among multimeric protein clusters. For each predicted assembly, pairwise structural similarity was quantified using both the query-template TM score (qTM) and the template-query TM-score (tTM) generated by Foldseek-Multimer. Because TM-scores are not inherently symmetric, the two values were averaged for each protein pair to yield a single symmetric similarity score ranging from 0 (no similarity) to 1 (identical structures). This procedure ensured that structural similarity was measured consistently regardless of which protein was designated as the query. The resulting similarity scores were compiled into a symmetric matrix and converted into a dissimilarity measure by calculating 1-average (qTM, tTM) for each protein pair. This dissimilarity matrix was then used as input to the scikit-learn implementation of t-SNE^78^ to project the high-dimensional structural relationships into two dimensions for visualisation.

A Sankey diagram of the protein clusters was generated using SankeyMATIC (https://sankeymatic.com/). Upset plots were generated using upsetplot (version 0.9.0).

## Experimental Validation

### Gene synthesis and cloning

Nucleotide sequences encoding phage proteins (UniProt IDs: Q38482, AXMXLFPO, P68658, Q38479, Q9T1X4, WHBPZJEF_CDS_0030, WEU69744, P03759, Q38673, and G9M973) were codon-optimised for expression in *E. coli* BL21 using the Integrated DNA Technologies (IDT) Codon Optimization Tool. Optimised gene sequences were synthesised as DNA fragments. Gene fragments were amplified by PCR using Q5 DNA polymerase (New England Biolabs) with 0.1 pmol synthetic DNA fragment as template and primers incorporating *Nco*I and *Xho*I restriction sites (Table S6). PCR amplification used an annealing temperature of 70°C. PCR products were analysed by electrophoresis on 0.8% agarose gels at 100 V for 1 hour, and DNA bands of expected size were excised and extracted. Both, PCR products and plasmid pASK-IBA63C (IBA Lifesciences, Germany), were digested with *Nco*I and *Xho*I restriction enzymes (New England Biolabs) at 37°C for 1 hour. Plasmid digestion included rSAP (recombinant shrimp alkaline phosphatase, New England Biolabs) to prevent plasmid re-ligation. Reactions were heat-inactivated at 80°C for 10 minutes, and digested fragments were analysed and purified by agarose gel electrophoresis followed by gel extraction.Digested inserts and plasmid were ligated using the Quick Ligation Kit (New England Biolabs) at room temperature for 1 hour, followed by ethanol precipitation with 1/10 volume of 3 M sodium acetate (pH 5.2) and 3 volumes of 100% ethanol overnight at -80°C. DNA was recovered by centrifugation at 14,000 × g for 15 minutes at 4°C, washed with 70% ethanol, air-dried, and resuspended in 30 μL nuclease-free water. Ligation products (10 μL) were transformed into chemically competent *E. coli* DH5α cells prepared according to Hanahan^79^ with modifications described by Inoue et al.^80^, and plated on LB agar containing 100 μg/mL ampicillin. Individual colonies were selected for plasmid isolation using the ZymoPURE Plasmid Miniprep Kit and analysed by *Nco*I / *Xho*I restriction and agarose gel electrophoresis to confirm successful cloning. Plasmid constructs were verified by Sanger sequencing using the primers listed in Table S6.

### Phage protein expression and analysis

Verified plasmid constructs were transformed into electrocompetent *E. coli* BL21(DE3) cells using optimised electroporation conditions: 12.5 kV/cm field strength (electroporator ECM 399, BTX Inc.), 200 Ω resistance, 100 ng/mL DNA, and 10¹⁰ cells/mL ^81 82^. The pASK-IBA63c vector (IBA Lifesciences, Germany) utilises a tetracycline-inducible expression system, providing tight regulation to prevent leaky expression and allowing controlled protein production upon addition of anhydrotetracycline (ATc). Transformed *E. coli* BL21(DE3) were grown in LB medium containing 100 μg/mL ampicillin at 37°C until reaching OD₆₀₀ = 0.5. The gene expression was induced with 50 ng/mL ATc for 3 hours at 30°C. Cells were harvested by centrifugation at 5,000 × g for 10 minutes at 4°C and resuspended in lysis buffer containing 50 mM Tris-HCl (pH 7.5), 150 mM NaCl, 10% glycerol, protease inhibitor cocktail(Roche,cOmplete, EDTA-free Protease Inhibitor Cocktail Tablets provided in EASYpack), and 0.25 mg/mL lysozyme (Carl Roth GmbH). Cell lysis was performed by sonication using 3 cycles of 45 seconds separated by 1 min off at 30% amplitude, with samples kept on ice throughout ^83^. Cell lysates were clarified by centrifugation at 12,000 × g for 15 minutes at 4°C, and total protein concentration in the soluble fraction was determined using the RotiNanoQuant assay (Carl Roth, Germany) according to manufacturer’s instructions.Protein production and purity were analysed by SDS-PAGE using 4-20% gradient polyacrylamide gels (8 cm × 8 cm × 1 mm; SERVAGel™ TG PRiME™ 4-20%, separation range 6-200 kDa). Protein samples (5 μg) were mixed 1:1 with Laemmli sample buffer containing 5% 2-mercaptoethanol, heated at 95°C for 5 minutes, and centrifuged briefly before loading. Gels were run in Laemmli running buffer at 50 mA for approximately 45 minutes at room temperature until adequate separation was achieved. Molecular weights of protein complexes were estimated using PageRuler™ Unstained Broad Range Protein Ladder (Thermo Fisher Scientific) and the SERVA Triple Color Protein-Standard II<u>I (SERVA)</u>. Proteins were visualised using SERVA Blue R Coomassie staining, followed by destaining in solution containing 20% ethanol, 5% acetic acid, and 1% glycerol.

### Protein purification using Strep-Tactin

*E. coli* lysate containing the recombinant Strep-tagged phage proteins were used directly for affinity purification with MagStrep “Strep-Tactin® XT” magnetic beads (IBA Lifesciences). All steps were carried out at 4 °C unless otherwise stated. Briefly, MagStrep “Strep-Tactin® XT” beads were equilibrated with wash buffer (100 mM Tris-HCl, 150 mM NaCl, 1 mM EDTA, pH 8.0) and then incubated with the clarified lysate for 30 min with gentle end-over-end mixing to allow binding of the tagged protein. After incubation, beads were captured using a magnetic rack and washed three times with wash buffer to remove unbound material.Bound protein was eluted under native conditions with biotin-containing elution buffer (100 mM Tris-HCl, 150 mM NaCl, 1 mM EDTA, pH 8.0, supplemented with 50 mM biotin) for 10 min at room temperature with gentle agitation. The supernatant containing the purified protein was collected by magnetic beads. Protein purity and yield were assessed by SDS-PAGE and protein concentration determination.To prepare the protein fractions analysed via size exclusion chromatography, affinity column chromatography was performed using Strep-Tactin® Superflow (high capacity, IBA Lifesciences) with *cell* lysates containing Px2c or Px3a, according to the manufacturer’s instructions.

### Blue native PAGE

Native protein complexes were analyzed by Blue Native (BN) PAGE using 4-20% polyacrylamide gradient gels (8 cm × 8 cm × 1 mm; SERVA*Gel*™ TG PRiME™ 4-20%). Purified protein samples were mixed with detergent-free sample buffer and 5 μg protein were loaded per lane. The SERVA Native Marker Liquid Mix for BN-PAGE was applied to calibrate the BN-PAGE. Electrophoresis was performed using light blue cathode buffer (50 mM Tricine, 15 mM Bis-Tris, pH 7.0, 0.002-0.05% Coomassie Brilliant Blue G-250) and Anode Buffer(50nM BisTris-HCl pH 7.0) at 50 V for 10 minutes, followed by 200 V until the dye front reached the gel edge. The Coomassie-containing cathode buffer was replaced with Coomassie-free cathode buffer after half of the electrophoresis run.

### Western blotting

Separated proteins were visualised by SERVA Blue R Coomassie staining. For immunoblot analysis, duplicate gels were equilibrated in transfer buffer (25 mM Tris, 192 mM glycine, 20% methanol) and proteins were transferred to PVDF membranes using wet blotting at 15 V, 80 mA for 60 minutes at 4°C. PVDF membranes were briefly activated in 100% methanol. Membranes were blocked in PBST (140 mM NaCl, 10 mM KCl, 6.4 mM Na₂HPO₄, 2 mM KH₂PO₄, 0.05% Tween 20) containing 3% skim milk powder for 1 hour at room temperature.Blocked membranes were incubated with Strep-Tactin AP conjugate (1:4000 dilution) for 60 minutes at room temperature, followed by washing with PBST. Protein detection was performed using NBT/BCIP substrate solution (Thermo Fisher Scientific) according to manufacturer’s instructions.

### Size exclusion chromatography

Analytical size exclusion chromatography was performed using a Superdex 75 Increase 10/300 GL column (GE Healthcare) equilibrated in Buffer W (100 mM Tris-HCl 150 mM NaCl, 1 mM EDTA pH 8.0,) at a flow rate of 0.650 mL/min and a temperature of 8°C. The column void volume (V₀) was determined using Blue Dextran 2000 (1 mg/mL) in 50 mM Bis-Tris, 50 mM NaCl, pH 6.5. Protein samples were applied in 900 µL volumes: Px3a at 3.3 mg/mL (3 mg total) and Px2c at 2.8 mg/mL (2.5 mg total). Elution was monitored by absorbance at 280 nm. The column was calibrated using two protein standard mixtures in Buffer W: Mix A (ribonuclease A, 13.7 kDa; carbonic anhydrase, 29 kDa; conalbumin, 75 kDa) and Mix B (aprotinin, 6.5 kDa; ribonuclease A, 13.7 kDa; ovalbumin, 43 kDa).

Standards were applied at 3 mg/mL except ovalbumin (4 mg/mL). Elution volumes (Vₑ) were measured from the point of sample application to peak maximum. Partition coefficients (Kₐᵥ) were calculated as Kₐᵥ = (Vₑ - V₀)/(V1 - V₀), where V1 is the total bed volume. A calibration curve was generated by linear regression of log(M_W_) versus Kₐᵥ, and apparent molecular weights of sample proteins were determined from this standard curve.

### Hydrodynamic calculations

Hydrodynamic properties were calculated from AlphaFold-predicted trimeric (Px3a) and dimeric (Px2c) structures using ZENO (Monte Carlo boundary element method) implemented in the UltraScan SOMO suite ^47^. Four different bead modelling methods were employed to calculate hydrodynamic parameters: the SoMo bead model, the SoMo overlap bead model, the AtoB grid-based bead model, and the van der Waals (vdW) overlap bead model. Hydrodynamic calculations were performed using the ZENO method with 1,000,000 Monte Carlo walks per structure, at pH 8.0 and 8°C to match experimental SEC conditions, using Buffer_W (100 mM Tris-HCl 150 mM NaCl, 1 mM EDTA pH 8.0) solvent properties (viscosity 1.72 cP, density 1.022 g/mL). Partial specific volumes (vbar) of 0.740 cm³/g and 0.732 cm³/g were used for Px2c and Px3a, respectively. All calculations were performed using default US-SOMO parameters unless otherwise stated.

### HullRad convex hull analysis

Protein shape analysis and frictional ratio calculations were performed using HullRad ^49^. HullRad computes the convex hull—the minimal bounding surface encompassing all protein atoms from three-dimensional structural coordinates. Crystal structures for reference proteins (PDB: 3LDJ; 1OVA; 1AFK; 2CAB; 8FEI) and homology models for experimental proteins (Px2c, Px3a) were used as input. HullRad calculates the frictional ratio (f/f₀) by approximating the convex hull as a tri-axial ellipsoid and applying Perrin’s equations. The frictional ratio represents the protein’s translational frictional coefficient relative to a sphere of equivalent molecular weight. Principal axis dimensions (X, Y, Z) were determined from the convex hull geometry, representing maximum extents along orthogonal directions. Axial ratios were calculated as pairwise ratios of principal dimensions (X:Y, X:Z, Y:Z).

## Acknowledgements

This work was supported by resources and services from Flinders University using the DeepThought High Performance Cluster (https://doi.org/10.25957/FLINDERS.HPC.DEEPTHOUGHT)^84^, the Phoenix HPC at Adelaide University, and the National Computational Infrastructure (NCI) and the Pawsey Supercomputing Research Centre (computing support was provided by a National Computational Merit Allocation Scheme grant to R.A.E.), which are supported by the Australian Government. We would like to thank Sarah Beecroft for her assistance in operating ColabFold at scale at Pawsey, especially for containerising ColabFold for use on Setonix’s AMD GPUs. We also acknowledge Michael Heinzinger and Sandra Studenik for useful discussions and Tiyasa Adhikary for her assistance in protein purification attempts. N.G., T.S., and B.E.D. were supported by the European Research Council (ERC) Consolidator grant 865694: DiversiPHI, the Deutsche Forschungsgemeinschaft (DFG, German Research Foundation) under Germany’s Excellence Strategy – EXC 2051 – via the Cluster of Excellence ‘Balance of the Microverse’ (Project-ID 390713860, to U.A.H., B.E.D.), and the Alexander von Humboldt Foundation in the context of an Alexander von Humboldt-Professorship founded by the German Federal Ministry of Education and Research. R.A.E. was supported by an award from the NIH NIDDK RC2DK116713 and awards from the Australian Research Council DP220102915, DP250103825, and FL250100019. S.R.G was supported by an Australian Government Research Training Program Scholarship and the Flinders University Overseas Travelling Fellowship.

## Author Contributions Statement

B.E.D, R.A.E, S.R.G and N.G conceived the project. S.R.G, V.M and G.B performed the bioinformatic analysis and protein structure predictions. N.G, T.S, and M.S performed the experimental work. S.R.G and N.G wrote the paper. B.E.D, R.A.E and U.A.H supervised the project. All authors read and approved the final manuscript.

## Competing Interests Statement

The authors declare no competing interests.

## Data Availability

Protein structures are publicly available at the Phage Protein Structural Gallery (https://linsalrob.github.io/PHROG_structures/).

## Code Availability

Phage Homomer Level Estimate and Generation Method (PHLEGM) is available at https://github.com/susiegriggo/phlegm.

## Notes

### Competing Interest Statement

The authors have declared no competing interest.

## References

1. Rihtman, B., Meaden, S., Clokie, M. R. J., Koskella, B. & Millard, A. D. Assessing Illumina technology for the high-throughput sequencing of bacteriophage genomes. PeerJ 4, e2055 (2016).

2. Cook, R. et al. INfrastructure for a PHAge REference Database: Identification of Large-Scale Biases in the Current Collection of Cultured Phage Genomes. PHAGE 2, 214–223 (2021).

3. Saghaei, S. et al. VirJenDB: a FAIR (meta)data and bioinformatics platform for all viruses. Nucleic Acids Res. gkaf1224 (2025) doi:10.1093/nar/gkaf1224.

4. Fiamenghi, M. B. et al. Meta-virus resource (MetaVR): expanding the frontiers of viral diversity with 24 million uncultivated virus genomes. Nucleic Acids Res. (2025) doi:10.1093/nar/gkaf1283.

5. Nishijima, S., Fullam, A., Schmidt, T. S. B., Kuhn, M. & Bork, P. VIRE: a metagenome-derived, planetary-scale virome resource with environmental context. Nucleic Acids Res. (2025) doi:10.1093/nar/gkaf1225.

6. Bouras, G. et al. Pharokka: a fast scalable bacteriophage annotation tool. Bioinformatics 39, (2023).

7. McNair, K., Zhou, C., Dinsdale, E. A., Souza, B. & Edwards, R. A. PHANOTATE: a novel approach to gene identification in phage genomes. Bioinformatics 35, 4537–4542 (2019).

8. Grigson, S. R., Bouras, G., Dutilh, B. E., Olson, R. D. & Edwards, R. A. Computational function prediction of bacteria and phage proteins. Microbiol. Mol. Biol. Rev. e0002225 (2025) doi:10.1128/mmbr.00022-25.

9. García-Bellido, A. Symmetries throughout organic evolution. Proc. Natl. Acad. Sci. U. S. A. 93, 14229–14232 (1996).

10. Pang, T., Savva, C. G., Fleming, K. G., Struck, D. K. & Young, R. Structure of the lethal phage pinhole. Proc. Natl. Acad. Sci. U. S. A. 106, 18966–18971 (2009).

11. Pillai, S. S. & Jain, V. Elucidating the role of mycobacteriophage D29-encoded Gp36 in DNA binding and phage gene expression regulation. Nucleic Acids Res. 53, (2025).

12. Sun, L. et al. Icosahedral bacteriophage ΦX174 forms a tail for DNA transport during infection. Nature 505, 432–435 (2014).

13. Podgorski, J. M. et al. Stabilization mechanism accommodating genome length variation in evolutionarily related viral capsids. Nat. Commun. 16, 3145 (2025).

14. Wikoff, W. R. et al. Topologically linked protein rings in the bacteriophage HK97 capsid. Science 289, 2129–2133 (2000).

15. Bartual, S. G. et al. Structure of the bacteriophage T4 long tail fiber receptor-binding tip. Proc. Natl. Acad. Sci. U. S. A. 107, 20287–20292 (2010).

16. Klein-Sousa, V., Roa-Eguiara, A., Kielkopf, C. S., Sofos, N. & Taylor, N. M. I. RBPseg: Toward a complete phage tail fiber structure atlas. Sci. Adv. 11, eadv0870 (2025).

17. Skaar, K. et al. Crystal structure of the bacteriophage P2 integrase catalytic domain. FEBS Lett. 589, 3556–3563 (2015).

18. Gupta, K., Sharp, R., Yuan, J. B., Li, H. & Van Duyne, G. D. Coiled-coil interactions mediate serine integrase directionality. Nucleic Acids Res. 45, 7339–7353 (2017).

19. Li, F. et al. High-resolution cryo-EM structure of the Pseudomonas bacteriophage E217. Nat. Commun. 14, 4052 (2023).

20. Bayfield, O. W. et al. Structural atlas of a human gut crassvirus. Nature 617, 409–416 (2023).

21. Bongirwar, V. & Mokhade, A. S. Different methods, techniques and their limitations in protein structure prediction: A review. Prog. Biophys. Mol. Biol. 173, 72–82 (2022).

22. Jumper, J. et al. Highly accurate protein structure prediction with AlphaFold. Nature 596, 583–589 (2021).

23. Abramson, J. et al. Accurate structure prediction of biomolecular interactions with AlphaFold 3. Nature 630, 493–500 (2024).

24. van Kempen, M. et al. Fast and accurate protein structure search with Foldseek. Nat. Biotechnol. 42, 243–246 (2024).

25. Holm, L. Dali server: structural unification of protein families. Nucleic Acids Res. 50, W210–5 (2022).

26. Bouras, G. et al. Protein structure-informed bacteriophage genome annotation with Phold. Nucleic Acids Res. 54, (2026).

27. Evans, R., et al. Protein complex prediction with AlphaFold-Multimer. bioRxiv (2021) doi:10.1101/2021.10.04.463034.

28. Han, Y., et al. AlphaFold Database expands to proteome-scale quaternary structures. bioRxiv (2026) doi:10.64898/2026.03.27.714458.

29. Odai, R. et al. The Viral AlphaFold Database of monomers and homodimers reveals conserved protein folds in viruses of bacteria, archaea, and eukaryotes. Sci. Adv. 11, eadz8560 (2025).

30. Schweke, H. et al. An atlas of protein homo-oligomerization across domains of life. Cell (2024) doi:10.1016/j.cell.2024.01.022.

31. Lin, Y., Wallis, C. & Corry, B. AlphaFold can be used to predict the oligomeric states of proteins. bioRxiv (2025) doi:10.1101/2025.03.10.642518.

32. Madaj, R., Martinez-Goikoetxea, M., Kaminski, K., Ludwiczak, J. & Dunin-Horkawicz, S. Applicability of AlphaFold2 in the modeling of dimeric, trimeric, and tetrameric coiled-coil domains. Bioinformatics (2024).

33. Shor, B. & Schneidman-Duhovny, D. CombFold: predicting structures of large protein assemblies using a combinatorial assembly algorithm and AlphaFold2. Nat. Methods 21, 477–487 (2024).

34. Avraham, O., Tsaban, T., Ben-Aharon, Z., Tsaban, L. & Schueler-Furman, O. Protein language models can capture protein quaternary state. BMC Bioinformatics 24, 433 (2023).

35. Kshirsagar, M. et al. Rapid and accurate prediction of protein homo-oligomer symmetry using Seq2Symm. Nat. Commun. 16, 2017 (2025).

36. Terzian, P., et al. PHROG: families of prokaryotic virus proteins clustered using remote homology. NAR Genom Bioinform 3, lqab067 (2021).

37. Carriles, A. A. et al. Structure-function insights into the dual role in nucleobase and nicotinamide metabolism and a possible use in cancer gene therapy of the URH1p riboside hydrolase. Int. J. Mol. Sci. 25, 7032 (2024).

38. Bad Phages in Good Bacteria: Role of the Mysterious orf63 of Lambda and Shiga Toxin-Converting Phi 24 B Bacteriophages.

39. Lin, Z. et al. Evolutionary-scale prediction of atomic-level protein structure with a language model. Science 379, 1123–1130 (2023).

40. Tao, Y. et al. Assembly of a tailed bacterial virus and its genome release studied in three dimensions. in Selected Papers of Michael G Rossmann with Commentaries 280–286 (WORLD SCIENTIFIC, 2014). doi:10.1142/9789814513357_0032.

41. Gräslund, S. et al. The use of systematic N- and C-terminal deletions to promote production and structural studies of recombinant proteins. Protein Expr. Purif. 58, 210–221 (2008).

42. Zhang, K. et al. Systemic expression, purification, and initial structural characterization of bacteriophage T4 proteins without known structure homologs. Front. Microbiol. 12, 674415 (2021).

43. Verde, V. L., Dominici, P. & Astegno, A. Determination of Hydrodynamic Radius of Proteins by Size Exclusion Chromatography. Bio Protoc. 7, e2230 (2017).

44. Burgess, R. R. A brief practical review of size exclusion chromatography: Rules of thumb, limitations, and troubleshooting. Protein Expr. Purif. 150, 81–85 (2018).

45. Landreh, M. et al. Predicting the shapes of protein complexes through collision cross section measurements and database searches. Anal. Chem. 92, 12297–12303 (2020).

46. Rocco, M. & Byron, O. Computing translational diffusion and sedimentation coefficients: an evaluation of experimental data and programs. Eur. Biophys. J. 44, 417–431 (2015).

47. Brookes, E., Demeler, B., Rosano, C. & Rocco, M. The implementation of SOMO (SOlution MOdeller) in the UltraScan analytical ultracentrifugation data analysis suite: enhanced capabilities allow the reliable hydrodynamic modeling of virtually any kind of biomacromolecule. Eur. Biophys. J. 39, 423–435 (2010).

48. Brookes, E. & Rocco, M. Recent advances in the UltraScan SOlution MOdeller (US-SOMO) hydrodynamic and small-angle scattering data analysis and simulation suite. Eur. Biophys. J. 47, 855–864 (2018).

49. Fleming, P. J. & Fleming, K. G. HullRad: Fast Calculations of Folded and Disordered Protein and Nucleic Acid Hydrodynamic Properties. Biophys. J. 114, 856–869 (2 2018).

50. Kang, E.-H., Mansfield, M. L. & Douglas, J. F. Numerical path integration technique for the calculation of transport properties of proteins. Phys. Rev. E Stat. Nonlin. Soft Matter Phys. 69, 031918 (2004).

51. Roy, A., Bhardwaj, A. & Cingolani, G. Crystallization of the nonameric small terminase subunit of bacteriophage P22. Acta Crystallogr. Sect. F Struct. Biol. Cryst. Commun. 67, 104–110 (2011).

52. Niazi, M. et al. Biophysical analysis of Pseudomonas-phage PaP3 small terminase suggests a mechanism for sequence-specific DNA-binding by lateral interdigitation. Nucleic Acids Res. 48, 11721–11736 (2020).

53. Silva, G. H. & Belfort, M. Analysis of the LAGLIDADG interface of the monomeric homing endonuclease I-DmoI. Nucleic Acids Res. 32, 3156–3168 (2004).

54. Chandrashekaran, S., Manjunatha, U. H. & Nagaraja, V. KpnI restriction endonuclease and methyltransferase exhibit contrasting mode of sequence recognition. Nucleic Acids Res. 32, 3148–3155 (2004).

55. Wei, Z.-L. et al. Structural insights into the chaperone-assisted assembly of a simplified tail fiber of the myocyanophage Pam3. Viruses 14, 2260 (2022).

56. Kim, W. et al. Rapid and sensitive protein complex alignment with Foldseek-Multimer. Nat. Methods 22, 469–472 (2025).

57. Guan, J. et al. GOPhage: protein function annotation for bacteriophages by integrating the genomic context. Brief. Bioinform. 26, (2024).

58. Boulay, A., Leprince, A., Enault, F., Rousseau, E. & Galiez, C. Empathi: Embedding-based phage protein annotation tool by hierarchical assignment. bioRxiv 2024.12.31.630607 (2025) doi:10.1101/2024.12.31.630607.

59. Flamholz, Z. N., Biller, S. J. & Kelly, L. Large language models improve annotation of viral proteins. Res. Sq. (2023) doi:10.21203/rs.3.rs-2852098/v1.

60. Grigson, S. R., et al. Synteny-aware functional annotation of bacteriophage genomes with Phynteny. bioRxiv (2025) doi:10.1101/2025.07.28.667340.

61. Bouras, G., et al. Protein structure informed bacteriophage genome annotation with phold. bioRxiv (2025) doi:10.1101/2025.08.05.668817.

62. Passaro, S., et al. Boltz-2: Towards accurate and efficient binding affinity prediction. bioRxivorg (2025) doi:10.1101/2025.06.14.659707.

63. Wohlwend, J., et al. Boltz-1 democratizing biomolecular interaction modeling. bioRxivorg (2025) doi:10.1101/2024.11.19.624167.

64. Guo, X. & He, Z.-G. Pangenome-scale annotation of mycobacteriophages for dissecting phage-host interactions based on a sequence clustering and structural homology analysis strategy. mSystems e0050825 (2025) doi:10.1128/msystems.00508-25.

65. Gnanasekar, P., Gambhir, S., Kinatukara, P. & Bhardwaj, A. Functional (re)annotation of Mycobacteroides abscessus proteome using integrative sequence and AI-based structural approaches. Curr. Res. Struct. Biol. 100172 (2025) doi:10.1016/j.crstbi.2025.100172.

66. Awad, M. et al. LicD-mediated cell wall decoration governs phage sensitivity in Enterococcus faecalis clinical isolates. Microbiol. Res. 302, 128341 (2026).

67. Novy, N., Huss, P., Evert, S., Romero, P. A. & Raman, S. Multiobjective learning and design of bacteriophage specificity. bioRxivorg (2025) doi:10.1101/2025.05.19.654895.

68. Mirdita, M. et al. ColabFold: making protein folding accessible to all. Nat. Methods 19, 679–682 (2022).

69. Haghani, M., Bhattacharya, D. & Murali, T. M. NEFFy: A versatile tool for computing the Number of Effective Sequences. Bioinformatics (2025) doi:10.1093/bioinformatics/btaf222.

70. Behzadi, P. & Gajdács, M. Worldwide Protein Data Bank (wwPDB): A virtual treasure for research in biotechnology. Eur. J. Microbiol. Immunol. (Bp.) 11, 77–86 (2022).

71. Steinegger, M. & Söding, J. MMseqs2 enables sensitive protein sequence searching for the analysis of massive data sets. Nat. Biotechnol. 35, 1026–1028 (2017).

72. Varadi, M. et al. AlphaFold Protein Structure Database: massively expanding the structural coverage of protein-sequence space with high-accuracy models. Nucleic Acids Res. 50, D439–D444 (2022).

73. Mariani, V., Biasini, M., Barbato, A. & Schwede, T. lDDT: a local superposition-free score for comparing protein structures and models using distance difference tests. Bioinformatics 29, 2722–2728 (2013).

74. Bryant, P., Pozzati, G. & Elofsson, A. Improved prediction of protein-protein interactions using AlphaFold2. Nat. Commun. 13, 1265 (2022).

75. Rodrigues, J. P. G. L. M., Teixeira, J. M. C., Trellet, M. & Bonvin, A. M. J. J. Pdb-tools: A swiss army knife for molecular structures. bioRxiv (2018) doi:10.1101/483305.

76. Zhang, C., Shine, M., Pyle, A. M. & Zhang, Y. US-align: universal structure alignments of proteins, nucleic acids, and macromolecular complexes. Nat. Methods 19, 1109–1115 (2022).

77. Pettersen, E. F. et al. UCSF ChimeraX: Structure visualization for researchers, educators, and developers. Protein Sci. 30, 70–82 (2021).

78. Van der Maaten, L. & Hinton, G. Visualizing data using t-SNE. J. Mach. Learn. Res. 9, (2008).

79. Hanahan, D. Studies on transformation of Escherichia coli with plasmids. J. Mol. Biol. 166, 557–580 (1983).

80. Inoue, H., Nojima, H. & Okayama, H. High efficiency transformation of Escherichia coli with plasmids. Gene 96, 23–28 (1990).

81. Dower, W. J., Miller, J. F. & Ragsdale, C. W. High efficiency transformation of E. coli by high voltage electroporation. Nucleic Acids Res. 16, 6127–6145 (1988).

82. Welch, M. et al. Design Parameters to Control Synthetic Gene Expression in Escherichia coli. PLoS One 4, e7002 (9 2009).

83. Kwon, Y.-C. & Jewett, M. C. High-throughput preparation methods of crude extract for robust cell-free protein synthesis. Sci. Rep. 5, 8663 (2015).

84. Deepthought HPC. Research @ Flinders at https://researchnow.flinders.edu.au/en/equipments/deepthought-hpc/

