## Supplementary Information for "Computational prediction resolves thousands of homooligomeric phage protein structures"

### Contents

| | |
|---|---|
| Supplementary Notes | 2 |
| Supplementary Figures | 3 |
| Supplementary Tables | 15 |
| References | 23 |

### Supplementary Notes

#### Supplementary Note 1: Evaluation of metric combinations and model aggregation strategies

To assess whether combining multiple metrics enhanced classification, we trained logistic regression, Random forest, gradient boosting, and support vector machine models using either ipTM alone or the full set of metrics (Fig. S2D). Utilising ipTM alone yielded better performance than combining the full set of metrics. This likely reflects the strong correlations among complex- and interface-level measures such as pTM, ipTM, ipLDDT, and pDockQ2 (Fig. S3), suggesting that incorporating additional, redundant metrics introduces noise rather than improving predictive signal. We examined the effect of aggregating multiple AlphaFold-Multimer predictions by comparing mean versus maximum scores. Due to high computational requirements, we limited our analysis to a maximum of five generated models per complex (Fig. S2D). Mean scores consistently outperformed maxima, with ipTM reaching optimal performance when averaging five models (AUC = 0.81, AP = 0.85). Performance gains plateaued after three models, providing confidence that generating and evaluating more than five predictions would offer minimal benefit while incurring prohibitive computational costs.

### Supplementary Figures

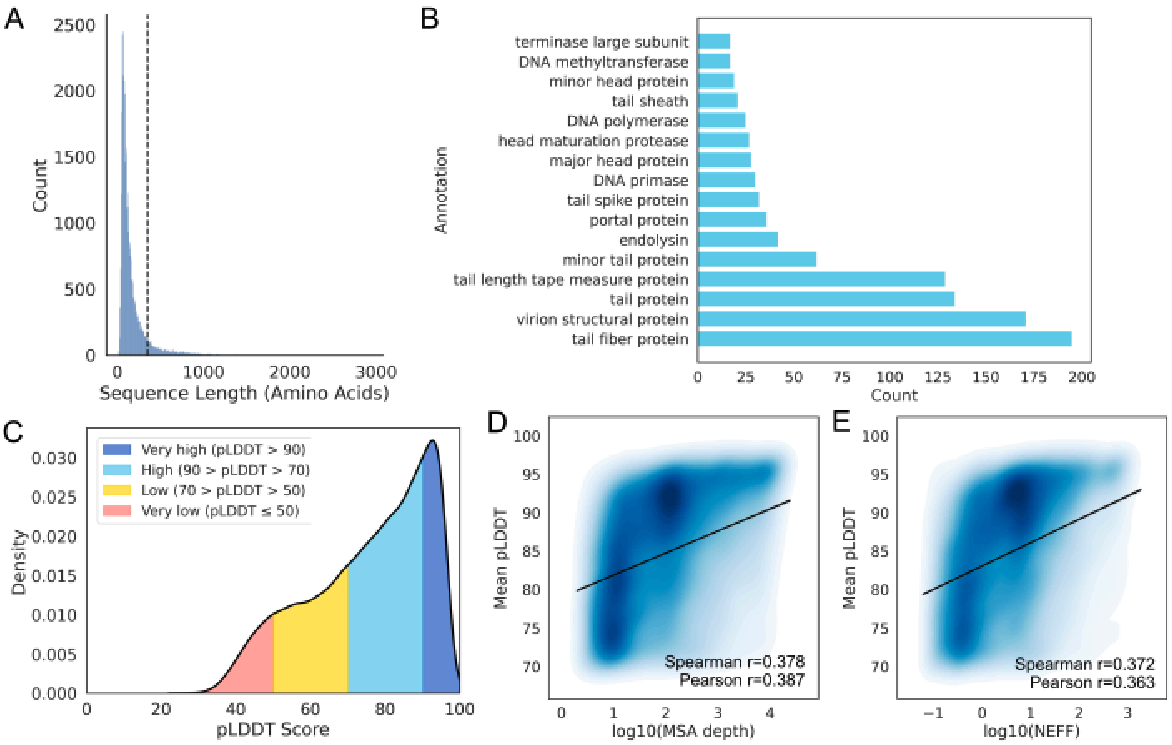


**Supplementary Figure 1: Filtering PHROG representatives and assessing AlphaFold2 confidence**

A. Distribution of PHROG representative sequence lengths, with the 350 amino acid threshold (dashed line) used to exclude sequences too long to fold with available GPU memory.
B. Functional annotations of PHROGs excluded due to exceeding this length threshold, highlighting an overrepresentation of large structural proteins such as tail fibers, tape measure proteins, and virion structural components. Annotations are shown which occurred at least 15 times in the excluded sequences.
C. Distribution of predicted mean pLDDT scores for AlphaFold2-generated structures of PHROG representative proteins.
D. Relationship between MSA depth and AlphaFold confidence (mean pLDDT). Density contours indicate the distribution of data points, and regression lines are overlaid. Pearson correlation: $r = 0.363$, $p < 1e–10$; Spearman correlation : $r = 0.372$, $p < 1e–10$.
E. Relationship between NEFF and AlphaFold confidence (mean pLDDT). Density contours indicate the distribution of data points, and regression lines are overlaid. Pearson correlation: r = 0.378, p < 1e–10; Spearman correlation: r = 0.387, p < 1e–10.

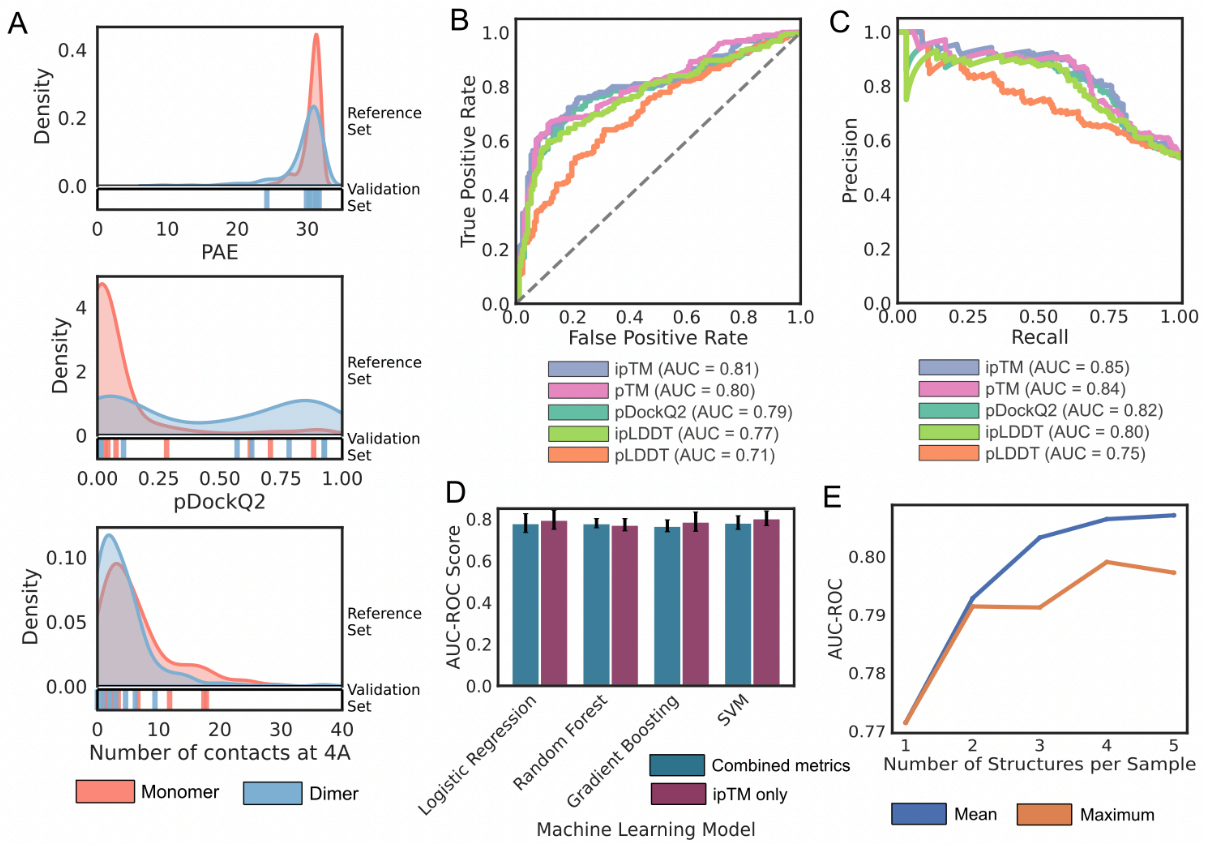


**Supplementary Figure 2: Evaluation of AlphaFold-Multimer metrics and structure aggregation strategies for predicting phage protein interfaces.**

A. Distributions of the PAE and the number of contacts within 4 Å for experimentally validated monomers (n=168) and dimers (n=196).
B. ROC curves assessing the discriminative power of AlphaFold-derived metrics taken as an average of five independent structures.
C. PR curves assessing the discriminative power of AlphaFold-derived metrics taken as an average of five independent structures.
D. Performance of machine learning models (AUC) trained on either ipTM alone or a combination of all metrics. Error bars show standard deviation of AUC-ROC across 5-fold cross-validation.
E. Effect of aggregating predictions across multiple independently generated structures on classification performance (AUC-ROC).

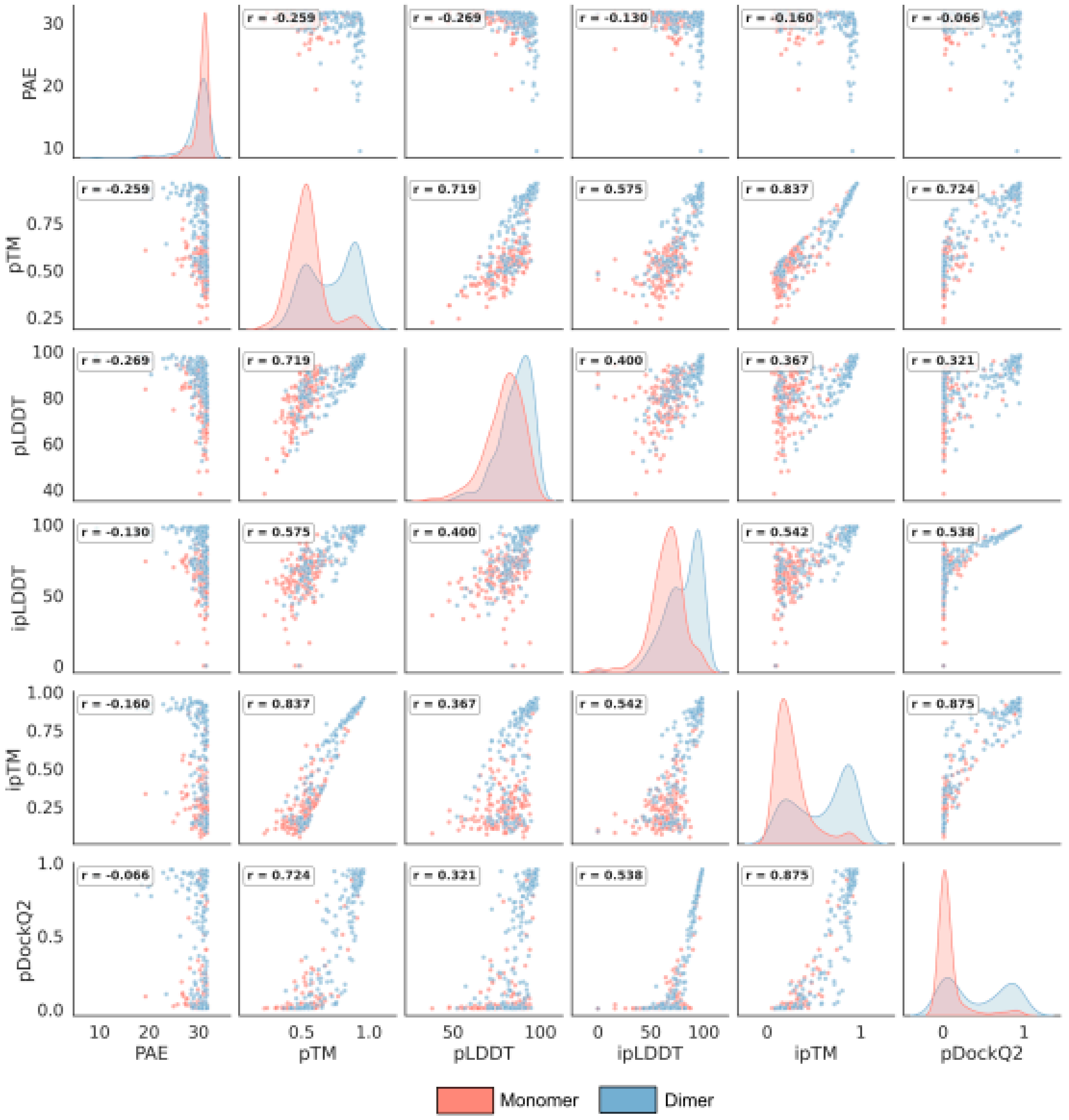


**Supplementary Figure 3: Relationship between structural metrics for monomer and dimer states.** Pearson correlation coefficients (r) are displayed for each plot. Kernel density estimates are shown along the diagonal, comparing the metrics.

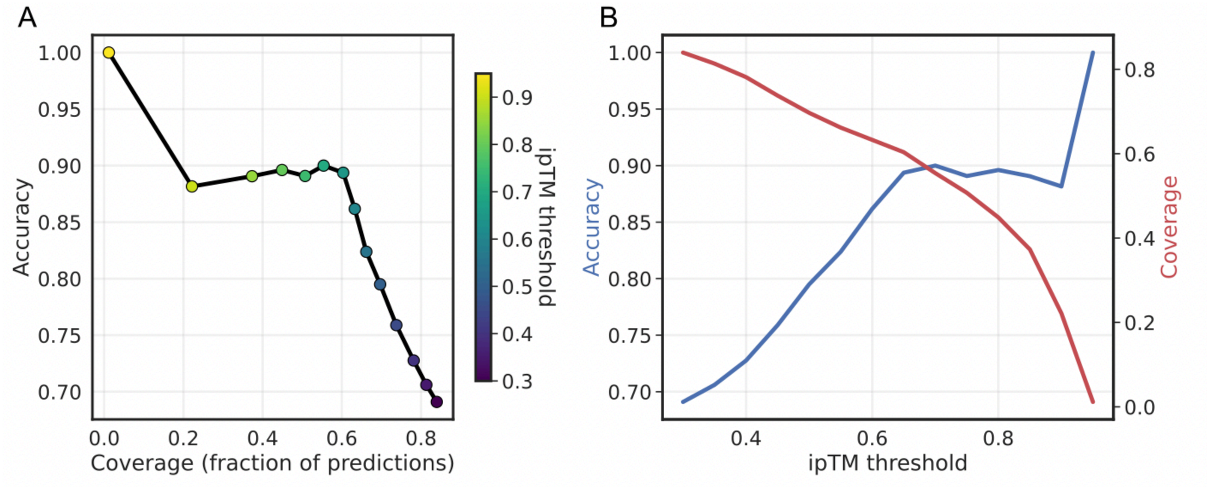


**Supplementary Figure 4: PHLEGM threshold analysis for oligomeric state prediction. A:** Accuracy-coverage trade-off plot showing the relationship between prediction accuracy and dataset coverage as a function of ipTM confidence threshold. Each point represents a different threshold value (0.3-1.0, indicated by colour gradient), with the connecting line showing the trade-off trajectory. **B:** Dual-axis plot displaying accuracy and coverage as functions of the ipTM threshold.

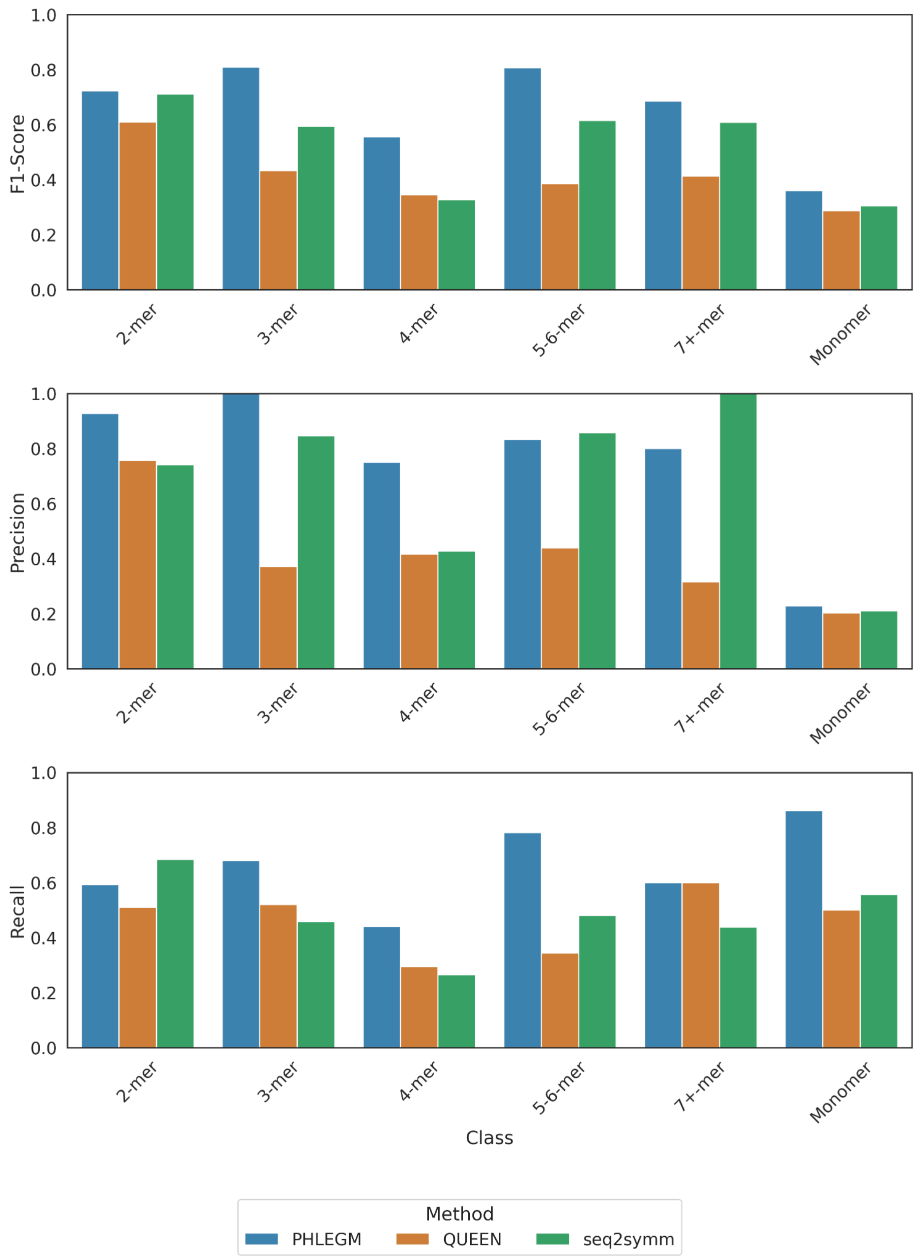


**Supplementary Figure 5: Per-class performance comparison across oligomerisation prediction methods.** Performance metrics for PHLEGM, QUEEN, and seq2symm methods evaluated across different oligomeric state classes (Monomer, 2-mer, 3-mer, 4-mer, 5-6-mer, and 7+-mer). **A:** F1-score comparison by class, showing the harmonic mean of precision and recall for each method across oligomeric states. **B:** Precision comparison by class, indicating the fraction of correct predictions among all positive predictions for each oligomeric state. **C:** Recall comparison by class, representing the fraction of actual positive cases correctly identified by each method. Error bars represent standard error across the discovery dataset.

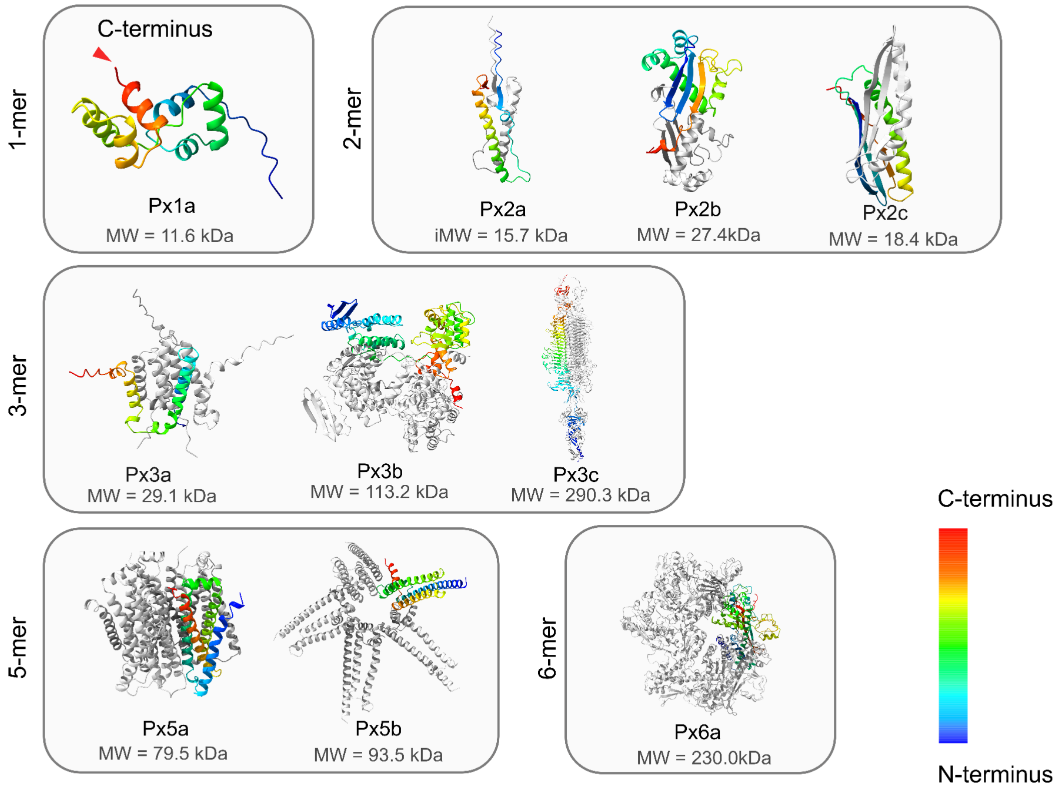


**Supplementary Figure 6.** Structural predictions of Px protein candidates grouped by oligomeric assembly (1-mer to 6-mer). Molecular weights are indicated for each construct. Rainbow coloring from N-terminus (blue) to C-terminus (red) is used to visualize monomer positions within each structure. The C-terminus was chosen for Strep-tag fusion based on its accessible localization, which minimizes interference with protein folding and oligomerization. The tag did not affect the protein folding or induced oligomerization.

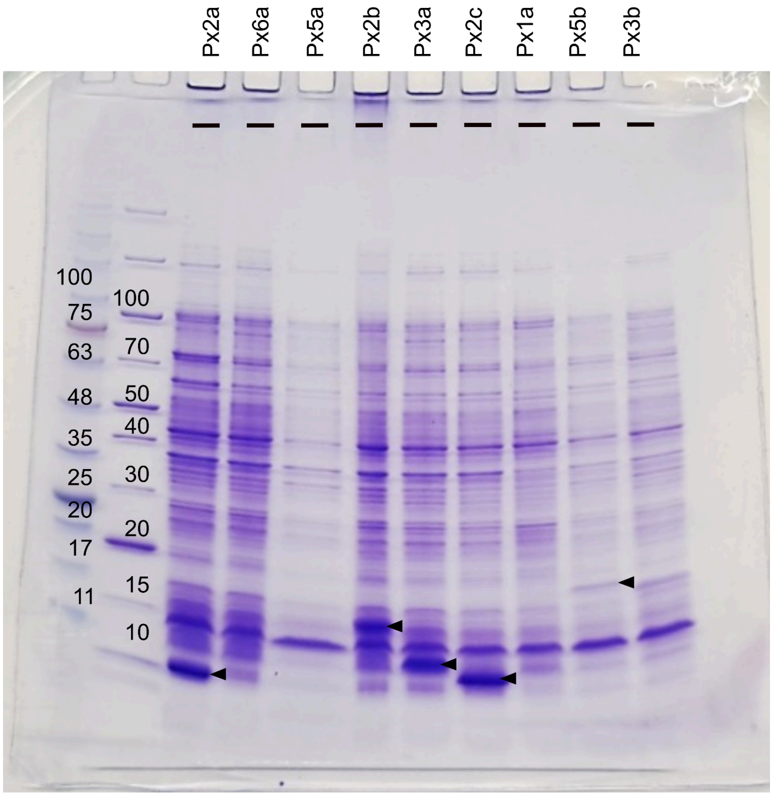


**Supplementary Figure 7.** SDS-PAGE analysis of candidate protein expression in *E. coli*. Coomassie-stained 4-20% SDS-PAGE gel showing expression screening results for nine candidate proteins Px2a, Px6a, Px5a, Px2b, Px3a, Px2c, Px1a, Px5b, and Px3b (lanes 1-9; 5 µg protein loaded per lane). Molecular weight markers are shown in the leftmost lane (kDa). Predicted molecular weights of successfully expressed proteins are to their respective lanes indicated with (8.9 kDa, 14.8 kDa, 10.7 kDa, 10.26 kDa and 19.7 kDa) black arrowheads.

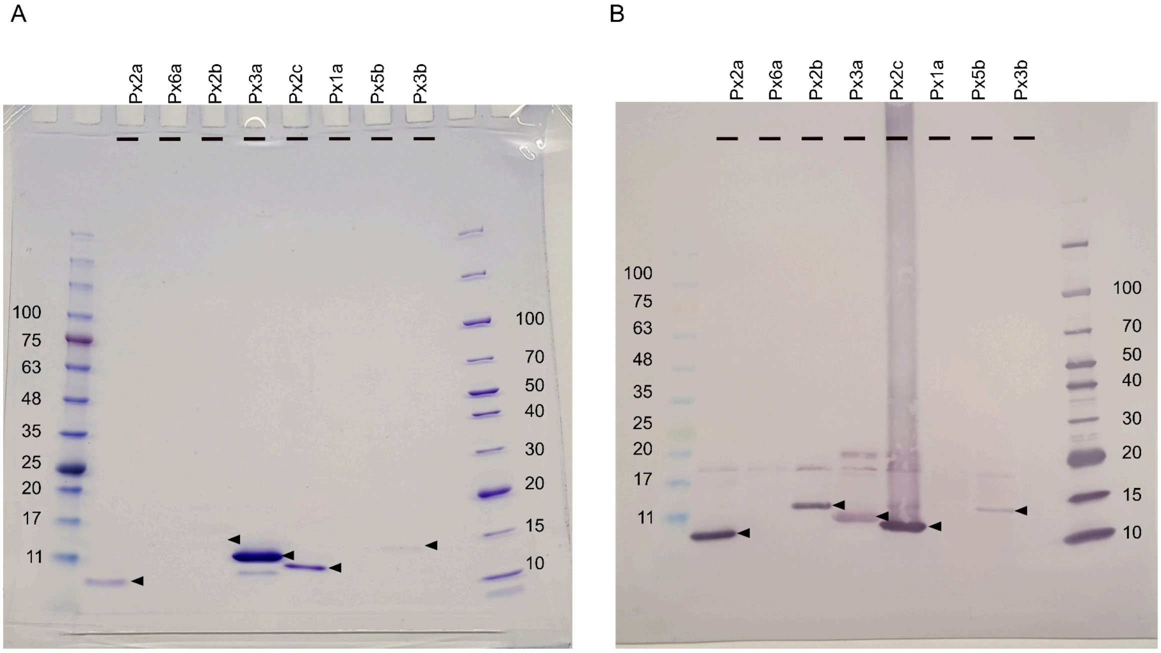


**Supplementary Figure 8.** SDS-PAGE analysis of purified proteins under denaturing conditions followed by anti-Strep-tag Western blot. Samples were separated on a 4-20% Tris-Glycine gradient gel and visualized by Coomassie staining (A) and anti-Strep-tag Western blot (B). Black arrowheads indicate the monomeric forms of the purified proteins migrating at their calculated molecular weights (~10-20 kDa range). Molecular weight markers (kDa) are shown on both sides.

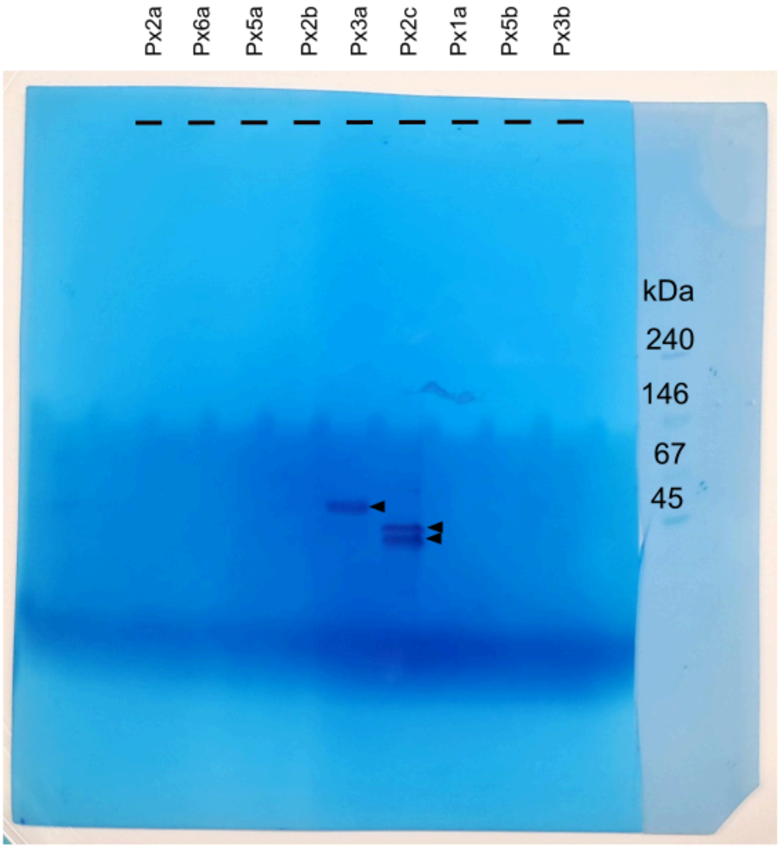


**Supplementary Figure 9.** Native oligomeric state analysis of purified proteins by Blue Native PAGE followed by anti-Strep-tag Western blot. Samples were separated on 4-20% Tris-Glycine gradient gels under native conditions (0.05% Coomassie in cathode buffer) and visualized by Coomassie staining and anti-Strep-tag Western blot. Black arrowheads indicate Px3a migrating at ~45 kDa and Px2c at a smaller than 45 kDa apparent molecular weight. The two arrowheads label two native states of protein Px2c showing different migration behavior. Molecular weight markers (kDa) are shown.

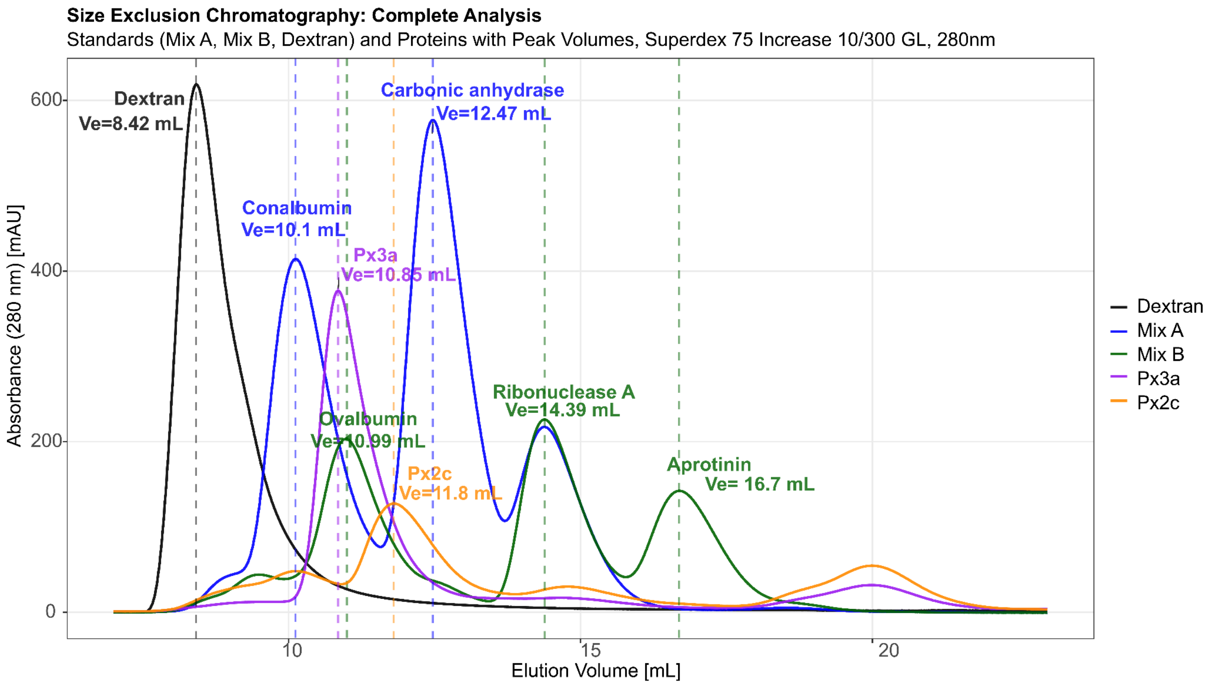


**Supplementary Figure 10.** Size exclusion chromatography analysis of purified proteins. Analytical SEC was performed on a Superdex 75 Increase 10/300 GL column equilibrated in Buffer W (100 mM Tris-HCl, 150 mM NaCl, 1 mM EDTA, pH 8.0) at 0.650 mL/min. Protein standard mixtures Mix A (ribonuclease A, 13.7 kDa; carbonic anhydrase, 29 kDa; conalbumin, 75 kDa) and Mix B (aprotinin, 6.5 kDa; ribonuclease A, 13.7 kDa; ovalbumin, 43 kDa) were used for calibration (gray traces). Sample proteins Px3a (blue, 3.3 mg/mL) and Px2c (orange, 2.8 mg/mL) were applied in 900 µL volumes. Elution was monitored by absorbance at 280 nm. The elution profiles showed apparent molecular weights for both proteins under native conditions, used to generate the calibration curve for molecular weight determination.

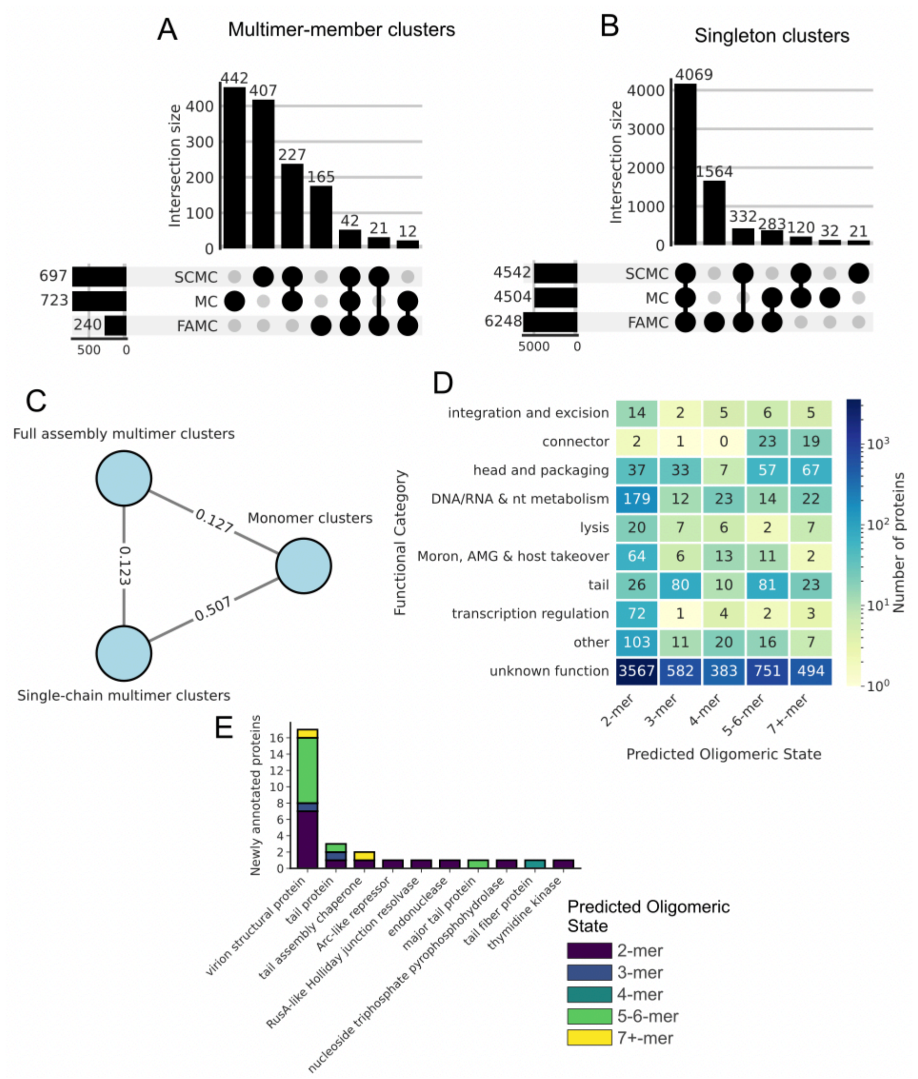


**Supplementary Figure 11:** Clustering of homooligomeric phage proteins.

A. UpSet plot showing the intersection of multi-member clusters that were identical across three clustering methods: monomer clusters (MC), single-chain multimer clusters (SCMC), and full assembly multimer clusters (FAMC).
B. UpSet plot of singleton clusters highlighting the extent to which unique proteins were consistently recovered by each clustering approach in (A).
C. Agreement between clustering strategies visualised as a network of adjusted Rand indices (ARI) comparing the three methods.
D. Heatmap displaying the distribution of predicted homooligomeric states across functional categories for PHROG representatives with known (n=1,125) and unknown (n=5,777) functions. Predictions were generated using PHLEGM with an ipTM threshold of 0.65, with protein counts annotated within each cell and colored on a logarithmic scale.
E. Functional reassignment of previously uncharacterised proteins. The stacked bar chart shows the top putative functional categories assigned to proteins that were structurally orphaned (singletons or clustered exclusively with unknowns) in the monomeric analysis but successfully mapped to annotated clusters using the SCMC representation. Bars are subdivided by the predicted oligomeric state of the newly annotated proteins. The sixteen most reassigned PHROG annotations are shown, all annotations shown in Supplementary Table 11.

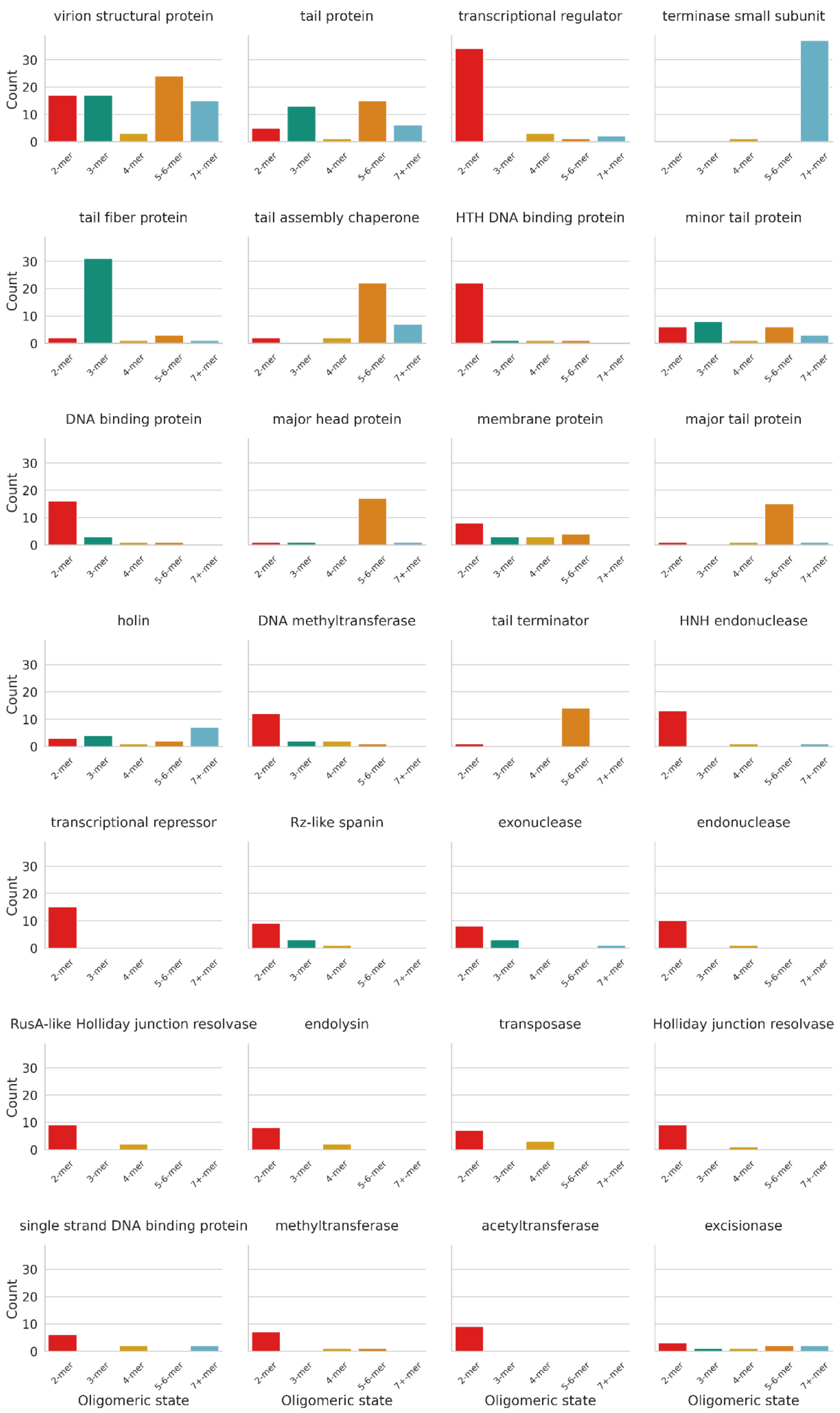


**Supplementary Figure 12:** Predicted oligomeric state distributions for the 28 most frequent protein annotations in the PHROG dataset. Each panel shows the count of proteins assigned to one of five oligomeric state categories (2-mer, 3-mer, 4-mer, 5–6-mer, and 7+-mer) based on AlphaFold-Multimer predictions. Note that monomeric states are not explicitly predicted as they could not be reliably predicted (Fig. 3D).

### Supplementary Tables

**Supplementary Table 1:** Distribution of homooligomeric states in the reference and validation sets. The Reference Set comprises structures deposited before the AlphaFold-Multimer v2.3 training cutoff (30 September 2021) and the validation set comprises structures deposited after this date.

| Oligomeric State | Reference Set (n) | Validation Set (n) |
|---|---|---|
| Monomer | 168 | 12 |
| Dimer | 196 | 8 |
| Trimer | 25 | 2 |
| Tetramer | 34 | 3 |
| 5-6-mers | 32 | 9 |
| 7+-mers | 20 | 5 |
| **Total** | **519** | **39** |

**Supplementary Table 2:** ipTM scores of PHROG representatives with dimeric and monomeric PDB hits deposited after the AlphaFold-Multimer training cutoff date.

| PHROG | PDB hit | PDB oligomeric state | ipTM | Notes | Reference |
|---|---|---|---|---|---|
| phrog_255 | 8kel | Homo 2-mer | 0.918 | | No publication |
| phrog_1384 | 8bv8 | Homo 2-mer | 0.876 | | No publication |
| phrog_8702 | 8rih | Monomer | 0.862 | Predominantly monomeric but forms disulfide-linked dimers via cysteine oxidation as evidenced by analytical ultracentrifugation and SDS-PAGE | (1) |
| phrog_38247 | 7u5q | Monomer | 0.86 | | No publication |
| phrog_292 | 8dsb | Homo 2-mer | 0.768 | | (2) |
| phrog_755 | 7t26 | Monomer | 0.762 | Monomer in the crystal structure and has activity in purified form. Related phosphoesterases sometimes multimerise so is possible that it may oligomerise. | (3) |
| phrog_30232 | 7xmw | Homo 2-mer | 0.762 | Dimer in crystal structure and size exclusion chromatography. | (4) |
| phrog_1856 | 7xl7 | Homo 2-mer | 0.722 | | No publication |
| phrog_21966 | 7pvc | Monomer | 0.534 | Monomer in the crystal structure. No complementary solution experiments showing the oligomeric state. | (5) |
| phrog_12654 | 8twq | Homo 2-mer | 0.444 | Dimer using gel filtration chromatography | (6) |
| phrog_1017 | 8bau | Monomer | 0.334 | Crystallised as a monomer | (7) |
| phrog_2860 | 7q4s | Monomer | 0.33 | Cyrstallised as a monomer. | (8) |
| phrog_22474 | 8tw1 | Monomer | 0.24 | Crystallised as a monomer. | (9) |
| phrog_8786 | 8zey | Monomer | 0.236 | Crystallised as a monomer. Size exclusion chromatography shows a monomer. | (10) |
| phrog_17304 | 8ikr | Homo 2-mer | 0.206 | Crystallised as dimer. No evidence it is a dimer in solution. | (11) |
| phrog_14334 | 7wkp | Monomer | 0.194 | Forms a heterodimer. | (12) |
| phrog_866 | 8cx8 | Monomer | 0.162 | Forms a dimer in asymmetric unit. Uploaded as a monomer. | (13) |
| phrog_6364 | 8z2m | Homo 2-mer | 0.158 | Asymmetric unit shows a dimer. No evidence it is a dimer in solution. | (14) |
| phrog_998 | 8tw1 | Monomer | 0.14 | Crystallised as a monomer. | (9) |
| phrog_47 | 8dwj | Monomer | 0.12 | Crystallised as a monomer. | (15) |

**Supplementary Table 3:** Manual curation of oligomeric states for PHROG representatives deposited in the PDB as monomers. For each entry, the corresponding reference, PDB identifier(s), and deposited oligomeric state are listed. The curated oligomeric state reflects evidence from the primary literature or structural context suggesting the biologically relevant assembly. Notes provide clarification where publications explicitly determine, contradict, or fail to mention the oligomeric state.

Refer to Excel Spreadsheet 'Supplementary_Tables.xlsx'

**Supplementary Table 4:** Comparison of PHLEGM-predicted oligomeric assemblies with experimentally resolved protein structures from the Protein Data Bank (PDB). TM-scores are normalised by the length of the reference structure. Comparisons were not performed where the stoichiometry of the PHLEGM-predicted assembly did not match that of the corresponding PDB entry. Where multiple PDB structures were available, the structure selected for alignment is indicated in bold (lowest alignment e-value). Notes highlight cases of discrepancies between deposited and biologically relevant oligomeric states.

Refer to Excel Spreadsheet ‘Supplementary_Tables.xlsx’

**Supplementary Table 5.** Candidate bacteriophage proteins for oligomeric state validation

| No. | Protein ID | Gene ID | Length (aa) | Monomer MW (kDa)[a] | Monomer MW Streptag | Database Accession | Predicted Function | Predicted Oligomeric State | Multimer MW (kDa)[b] | Multimer MW (kDa) Streptag |
|---|---|---|---|---|---|---|---|---|---|---|
| 1 | Px1a | Q38482 | 104 | 11.59 | 12.63359 | Gene ID: 2636278 | Transcriptional regulator | Monomer | 11.6 | 12.63359 |
| 2 | Px2a | AXMXLF-CDS _0004 | 72 | 7.846 | 8.8864 | GenBank: WYM31395.1 | RZ-like spanin | Dimer | 15.7 | 17.7728 |
| 3 | Px2b | P68658 | 122 | 13.78 | 14.82053 | NCBI: WP_000065374.1 | Unknown (Tro phenotype) | Dimer | 27.6 | 29.64106 |
| 4 | Px2c | Q38479 | 93 | 9.217 | 10.25779 | NCBI: NC_000929.1 | hypothetical protein (gp7) | Dimer | 18.4 | 20.51558 |
| 5 | Px3a | Q9T1X4 | 99 | 9.686 | 10.7262 | NCBI: AAA32405.1 | hypothetical protein (gp15) | Trimer | 29.1 | 32.1786 |
| 6 | Px3b | WHBPZJ-CDS_0030 | 348 | 37.73 | 38.77734 | NCBI: WP_035366093.1 | Integrase/Recombinase | Trimer | 113.2 | 116.332 |
| 7 | Px3c | WEU69744 | 879 | 96.77 | 97.81171 | WEU69744.1 | Tailspike protein | Trimer | 290.3 | 293.4351 |
| 8 | Px5a | P03759 | 143 | 15.96 | 17.0073 | UniProtKB: P03759 | RexB exclusion protein | Pentamer | 79.8 | 85.0365 |
| 9 | Px5b | Q38674 | 169 | 18.7 | 19.74148 | NCBI: WP_015975200.1 | Superinfection exclusion | Pentamer | 93.5 | 98.7074 |
| 10 | Px6a | G9M97 | 417 | 46.06 | 47.10234 | UniProtKB: G9M973.1 | Major capsid protein | Pentamer/Hexamer | 230.3 | 235.5117 |

[a] Monomer molecular weights calculated using ExPASy ProtParam from amino acid sequences.

[b] Multimer molecular weights calculated from predicted oligomeric states (excluding affinity tags).

[c] Expression performed in *E. coli* BL21(DE3) with Strep-tag II

**Supplementary Table 6.** Primers used in this study. Restriction sites highlighted in bold: NcoI (CCATGG) in red, XhoI (CTCGAG) in blue, AfeI (AGCGCT) in green. F, forward primer; R, reverse primer.

| # | Primer Name | Sequence (5'->3') |
|---|---|---|
| 1 | F_AXMXLFPO_NcoI | GATCCCATGGTGCTTCTCACGTCTTG |
| 2 | R_AXMXLFPO_XhoI | GATCCTCGAGGTTTTTCATCCGAACTTTAG |
| 3 | F_G9M973_NcoI | GATCCCATGGCGGAGAAATCCACTAAA |
| 4 | R_G9M973_XhoI | GATCCTCGAGTTCTGTGCGAGGCGT |
| 5 | F_P03759_NcoI | GATCCCATGGGGAATCGCATCATGCC |
| 6 | R_p03759_XhoI | GATCCTCGAGCTCCTGCTCGCGAG |
| 7 | F_P68658_NcoI | GATCCCATGGCTAACATTAAGAAGTATATCATCGATTATG |
| 8 | R_P68658_XhoI | GATCCTCGAGCGCCGCCTTAATTGTC |
| 9 | F_Q9T1X4_NcoI | GATCCCATGGAGAACAATGTTCAACC |
| 10 | R_Q9T1X4_XhoI | GATCCTCGAGTTGCCGCTCGTGG |
| 11 | F_Q38479_NcoI | GATCCCATGGCAAAGGTTATTATTGAGAT |
| 12 | R_Q38479_XhoI | GATCCTCGAGGTGACAAGTCTTTTCGACC |
| 13 | F_Q38482_NcoI | GATCCCATGGTGACGGACATGAA |
| 14 | R_Q38482_XhoI | GATCCTCGAGATACCCCGAGTAGACAATA |
| 15 | F_Q38674_NcoI | GATCCCATGGCCGACAGTATGTCCTAT |
| 16 | R_Q38674_XhoI | GATCCTCGAGTTTAATAGTCTTGATGAATTCCTTATTG |
| 17 | F_WHBPZJEF_CDS_0030_NcoI | GATCCCATGGCTAGTATTATTAAACGTG |
| 18 | R_WHBPZJEF_CDS_0030_XhoI | GATCCTCGAGCCCTAATTTCTTAGCCAAATC |
| 19 | F_WEU69744_NcoI | GATCCCATGGATTTCACCCAAGAGGA |
| 20 | R_WEU69744_AfeI | GATCAGCGCTAACGATCACAACCTTTGAA |

**Supplementary Table 7**: Comparison of clustering outcomes for PHROG representatives using Foldseek with different strategies. Monomeric structures were clustered with Foldseek easy-cluster (monomers), predicted multimeric assemblies were clustered with Foldseek easy-cluster (multimers), and quaternary structures were clustered directly using Foldseek easy-multimer-cluster. Clustering was performed with default parameters.

| Clustering Method | Number of singletons | Number of clusters | Mean cluster size | Max size |
|---|---|---|---|---|
| Foldseek easy-cluster (monomers) | 4504 | 723 | 3.317 | 44 |
| Foldseek easy-cluster (multimers) | 4542 | 697 | 3.385 | 52 |
| Foldseek easy-multimer-cluster | 6248 | 240 | 2.725 | 15 |

**Supplementary Table 8:** Multimer-guided functional annotations for 506 previously uncharacterised PHROGs. List of proteins that were structurally orphaned at the monomer level but successfully mapped to characterised clusters using single-chain multimer representations

Refer to Excel Spreadsheet ‘Supplementary_Tables.xlsx’

#### References


1. Carriles AA, Muzzolini L, Minici C, Tornaghi P, Patrone M, Degano M. 2024. Structure-function insights into the dual role in nucleobase and nicotinamide metabolism and a possible use in cancer gene therapy of the URH1p riboside hydrolase. Int J Mol Sci 25:7032.

2. Dydecka A, Bloch S, Rizvi A, Perez S, Nejman-Falenczyk B, Topka G, Gasior T, Necel A, Wegrzyn G, Donaldson LW, Wegrzyn A. 2017. Bad phages in good bacteria: Role of the mysterious orf63 of λ and Shiga toxin-converting Φ24B bacteriophages. Front Microbiol 8:1618.

3. Hobbs SJ, Wein T, Lu A, Morehouse BR, Schnabel J, Leavitt A, Yirmiya E, Sorek R, Kranzusch PJ. 2022. Phage anti-CBASS and anti-Pycsar nucleases subvert bacterial immunity. Nature 605:522–526.

4. Song G, Li X, Wang Z, Dong C, Xie X, Yan X. 2022. Structure of AcrVIA2 and its binding mechanism to CRISPR-Cas13a. Biochem Biophys Res Commun 612:84–90.

5. Torres Cabán CC, Yang M, Lai C, Yang L, Subach FV, Smith BO, Piatkevich KD, Boyden ES. 2022. Tuning the sensitivity of genetically encoded fluorescent potassium indicators through structure-guided and genome mining strategies. ACS Sens 7:1336–1346.

6. Adams MC, Schiltz CJ, Sun J, Hosford CJ, Johnson VM, Pan H, Borbat PP, Freed JH, Thomason LC, Court C, Court DL, Chappie JS. 2024. The crystal structure of bacteriophage λ RexA provides novel insights into the DNA binding properties of Rex-like phage exclusion proteins. Nucleic Acids Res 52:4659–4675.

7. Schuller M, Raggiaschi R, Mikolcevic P, Rack JGM, Ariza A, Zhang Y, Ledermann R, Tang C, Mikoc A, Ahel I. 2023. Molecular basis for the reversible ADP-ribosylation of

guanosine bases. Mol Cell 83:2303–2315.e6.

8. Vázquez R, Seoane-Blanco M, Rivero-Buceta V, Ruiz S, van Raaij MJ, García P. 2022. Monomodular Pseudomonas aeruginosa phage JG004 lysozyme (Pae87) contains a bacterial surface-active antimicrobial peptide-like region and a possible substrate-binding subdomain. Acta Crystallogr D Struct Biol 78:435–454.

9. Oechslin F, Zhu X, Morency C, Somerville V, Shi R, Moineau S. 2024. Fermentation practices select for thermostable endolysins in phages. Mol Biol Evol 41.

10. Kim DY, Han JH, Lee SY, Ha HJ, Park HH. 2024. Novel structure of the anti-CRISPR protein AcrIE3 and its implication on the CRISPR-Cas inhibition. Biochem Biophys Res Commun 722:150164.

11. Wang H-J, Hernández-Rocamora VM, Kuo C-I, Hsieh K-Y, Lee S-H, Ho M-R, Tu Z, Vollmer W, Chang C-I. 2023. Structural basis for the hydrolytic activity of the transpeptidase-like protein DpaA to detach Braun's lipoprotein from peptidoglycan. MBio 14:e0137923.

12. Zhang M, Peng R, Peng Q, Liu S, Li Z, Zhang Y, Song H, Yang J, Xing X, Wang P, Qi J, Gao GF. 2023. Mechanistic insights into DNA binding and cleavage by a compact type I-F CRISPR-Cas system in bacteriophage. Proc Natl Acad Sci U S A 120:e2215098120.

13. Rudolph MJ, Tsymbal AM, Dutta A, Davis SA, Algava B, Roberge JY, Tumer NE, Li X-P. 2024. Fragment screening to identify inhibitors targeting ribosome binding of Shiga toxin 2. ACS Infect Dis 10:2814–2825.

14. Journal Article Structural insights into the biosynthetic mechanism of Nα-GlyT and 5-NmdU hypermodifications of DNA.

15. Feng X, Spiering MM, de Luna Almeida Santos R, Benkovic SJ, Li H. 2023. Structural basis of the T4 bacteriophage primosome assembly and primer synthesis. Nat Commun

14:4396.

16. Loschi L, Brokx SJ, Hills TL, Zhang G, Bertero MG, Lovering AL, Weiner JH, Strynadka NCJ. 2004. Structural and biochemical identification of a novel bacterial oxidoreductase. J Biol Chem 279:50391–50400.

17. Pääkkönen J, Hakulinen N, Andberg M, Koivula A, Rouvinen J. 2022. Three-dimensional structure of xylonolactonase from Caulobacter crescentus: A mononuclear iron enzyme of the 6-bladed β-propeller hydrolase family. Protein Sci 31:371–383.

18. Talagas A, Fontaine L, Ledesma-García L, Mignolet J, Li de la Sierra-Gallay I, Lazar N, Aumont-Nicaise M, Federle MJ, Prehna G, Hols P, Nessler S. 2017. Correction: Structural insights into streptococcal competence regulation by the cell-to-cell communication system ComRS. PLoS Pathog 13:e1006208.

19. Mashburn-Warren L, Goodman SD, Federle MJ, Prehna G. 2018. The conserved mosaic prophage protein paratox inhibits the natural competence regulator ComR in Streptococcus. Sci Rep 8:16535.

20. Kim I, Koo J, An SY, Hong S, Ka D, Kim E-H, Bae E, Suh J-Y. 2020. Structural and mechanistic insights into the CRISPR inhibition of AcrIF7. Nucleic Acids Res 48:9959–9968.

21. Paulin KA, Cortez D, Eichman BF. 2022. The SOS response-associated peptidase (SRAP) domain of YedK catalyzes ring opening of abasic sites and reversal of its DNA-protein cross-link. J Biol Chem 298:102307.

22. Thompson PS, Amidon KM, Mohni KN, Cortez D, Eichman BF. 2019. Protection of abasic sites during DNA replication by a stable thiazolidine protein-DNA cross-link. Nat Struct Mol Biol 26:613–618.

23. Wang N, Bao H, Chen L, Liu Y, Li Y, Wu B, Huang H. 2019. Molecular basis of abasic site sensing in single-stranded DNA by the SRAP domain of E. coli yedK. Nucleic Acids Res 47:10388–10399.

24. Teller DC, Behnke CA, Pappan K, Shen Z, Reese JC, Reeck GR, Stenkamp RE. 2014. The structure of rice weevil pectin methylesterase. Acta Crystallogr F Struct Biol Commun 70:1480–1484.

25. Eklöf JM, Tan T-C, Divne C, Brumer H. 2009. The crystal structure of the outer membrane lipoprotein YbhC from Escherichia coli sheds new light on the phylogeny of carbohydrate esterase family 8. Proteins 76:1029–1036.

26. Crystal structure of YbaK protein from Haemophilus influenzae (HI1434) at 1.8 Å resolution: Functional implications.

27. Yang Y, Yue Y, Song N, Li C, Yuan Z, Wang Y, Ma Y, Li H, Zhang F, Wang W, Jia H, Li P, Li X, Wang Q, Ding Z, Dong H, Gu L, Li B. 2020. The YdiU domain modulates bacterial stress signaling through Mn2+-dependent UMPylation. Cell Rep 32:108161.

28. Tomchick DR, Sreelatha A, Yee SS, Lopez VA, Park BC, Kinch LN, Pilch S, Servage KA, Zhang J, Jiou J, Karasiewicz-Urbańska M, Łobocka M, Grishin NV, Orth K, Kucharczyk R, Pawłowski K, Tagliabracci VS. 2019. Protein AMPylation by an evolutionarily conserved pseudokinase. Acta Crystallogr A Found Adv 75:a314–a314.

29. Chung J, Goo E, Yu S, Choi O, Lee J, Kim J, Kim H, Igarashi J, Suga H, Moon JS, Hwang I, Rhee S. 2011. Small-molecule inhibitor binding to an N-acyl-homoserine lactone synthase. Proc Natl Acad Sci U S A 108:12089–12094.

30. Kerr ID, Sivakolundu S, Li Z, Buchsbaum JC, Knox LA, Kriwacki R, White SW. 2007. Crystallographic and NMR analyses of UvsW and UvsW.1 from bacteriophage T4. J Biol Chem 282:34392–34400.

31. Zhang L, Xu D, Huang Y, Zhu X, Rui M, Wan T, Zheng X, Shen Y, Chen X, Ma K, Gong Y. 2017. Structural and functional characterization of deep-sea thermophilic bacteriophage GVE2 HNH endonuclease. Sci Rep 7:42542.

32. Ho CK, Wang LK, Lima CD, Shuman S. 2004. Structure and mechanism of RNA ligase. Structure 12:327–339.

33. Nandakumar J, Shuman S, Lima CD. 2006. RNA ligase structures reveal the basis for RNA specificity and conformational changes that drive ligation forward. Cell 127:71–84.

34. Dessau M, Goldhill D, McBride RC, Turner PE, Modis Y. 2012. Correction: Selective pressure causes an RNA virus to trade reproductive fitness for increased structural and thermal stability of a viral enzyme. PLoS Genet 8.

35. Salgado PS, Makeyev EV, Butcher SJ, Bamford DH, Stuart DI, Grimes JM. 2004. The structural basis for RNA specificity and Ca2+ inhibition of an RNA-dependent RNA polymerase. Structure 12:307–316.

36. Ilca SL, Kotecha A, Sun X, Poranen MM, Stuart DI, Huiskonen JT. 2015. Localized reconstruction of subunits from electron cryomicroscopy images of macromolecular complexes. Nat Commun 6:8843.

37. Wright S, Poranen MM, Bamford DH, Stuart DI, Grimes JM. 2012. Noncatalytic ions direct the RNA-dependent RNA polymerase of bacterial double-stranded RNA virus ϕ6 from de novo initiation to elongation. J Virol 86:2837–2849.

38. Rissanen I, Grimes JM, Pawlowski A, Mäntynen S, Harlos K, Bamford JKH, Stuart DI. 2013. Bacteriophage P23-77 capsid protein structures reveal the archetype of an ancient branch from a major virus lineage. Structure 21:718–726.

39. Khayat R, Tang L, Larson ET, Lawrence CM, Young M, Johnson JE. 2005. Structure of an archaeal virus capsid protein reveals a common ancestry to eukaryotic and bacterial

viruses. Proc Natl Acad Sci U S A 102:18944–18949.

40. Darling PJ, Holt JM, Ackers GK. 2000. Coupled energetics of lambda cro repressor self-assembly and site-specific DNA operator binding II: cooperative interactions of cro dimers. J Mol Biol 302:625–638.

41. Jana R, Hazbun TR, Mollah AK, Mossing MC. 1997. A folded monomeric intermediate in the formation of lambda Cro dimer-DNA complexes. J Mol Biol 273:402–416.

42. Maluf NK, Yang Q, Catalano CE. 2005. Self-association properties of the bacteriophage lambda terminase holoenzyme: implications for the DNA packaging motor. J Mol Biol 347:523–542.

43. Koller-Eichhorn R, Marquardt T, Gail R, Wittinghofer A, Kostrewa D, Kutay U, Kambach C. 2007. Human OLA1 defines an ATPase subfamily in the Obg family of GTP-binding proteins. J Biol Chem 282:19928–19937.

44. Cheung M-Y, Li X, Ku Y-S, Chen Z, Lam H-M. 2022. Co-crystalization reveals the interaction between AtYchF1 and ppGpp. Front Mol Biosci 9:1061350.

45. Teplyakov A, Obmolova G, Chu SY, Toedt J, Eisenstein E, Howard AJ, Gilliland GL. 2003. Crystal structure of the YchF protein reveals binding sites for GTP and nucleic acid. J Bacteriol 185:4031–4037.

46. Qin L, Fokine A, O’Donnell E, Rao VB, Rossmann MG. 2010. Structure of the small outer capsid protein, Soc: a clamp for stabilizing capsids of T4-like phages. J Mol Biol 395:728–741.

47. He X, Byrd AK, Yun M-K, Pemble CW 4th, Harrison D, Yeruva L, Dahl C, Kreuzer KN, Raney KD, White SW. 2012. The T4 phage SF1B helicase Dda is structurally optimized to perform DNA strand separation. Structure 20:1189–1200.

48. Kim Y, Skarina T, Beasley S, Laskowski R, Arrowsmith C, Joachimiak A, Edwards A,

Savchenko A. 2002. Crystal structure of Escherichia coli EC1530, a glyoxylate induced protein YgbM. Proteins 48:427–430.

49. Gui W-J, Qu Q-H, Chen Y-Y, Wang M, Zhang X-E, Bi L-J, Jiang T. 2011. Crystal structure of YdaL, a stand-alone small MutS-related protein from Escherichia coli. J Struct Biol 174:282–289.

50. Zhou J, Pecqueur L, Aučynaitė A, Fuchs J, Rutkienė R, Vaitekūnas J, Meškys R, Boll M, Fontecave M, Urbonavičius J, Golinelli-Pimpaneau B. 2021. Structural evidence for a [4Fe‐5S] intermediate in the non‐redox desulfuration of thiouracil. Angew Chem Weinheim Bergstr Ger 133:428–435.

51. Xiao Y, Chen H, Wang H, Zhang M, Chen X, Berk JM, Zhang L, Wei Y, Li W, Cui W, Wang F, Wang Q, Cui C, Li T, Chen C, Ye S, Zhang L, Ji X, Huang J, Wang W, Wang Z, Hochstrasser M, Yang H. 2021. Structural and mechanistic insights into the complexes formed by Wolbachia cytoplasmic incompatibility factors. Proc Natl Acad Sci U S A 118:e2107699118.

52. Wang H, Xiao Y, Chen X, Zhang M, Sun G, Wang F, Wang L, Zhang H, Zhang X, Yang X, Li W, Wei Y, Yao D, Zhang B, Li J, Cui W, Wang F, Chen C, Shen W, Su D, Bai F, Huang J, Ye S, Zhang L, Ji X, Wang W, Wang Z, Hochstrasser M, Yang H. 2022. Crystal structures of Wolbachia CidA and CidB reveal determinants of bacteria-induced cytoplasmic incompatibility and rescue. Nat Commun 13:1608.

53. Peixeiro N, Keller J, Collinet B, Leulliot N, Campanacci V, Cortez D, Cambillau C, Nitta KR, Vincentelli R, Forterre P, Prangishvili D, Sezonov G, van Tilbeurgh H. 2013. Structure and function of AvtR, a novel transcriptional regulator from a hyperthermophilic archaeal lipothrixvirus. J Virol 87:124–136.

54. Wang G, Klein MG, Tokonzaba E, Zhang Y, Holden LG, Chen XS. 2008. The structure of a DnaB-family replicative helicase and its interactions with primase. Nat Struct Mol

Biol 15:94–100.

55. Ramirez BE, Voloshin ON, Camerini-Otero RD, Bax A. 2000. Solution structure of DinI provides insight into its mode of RecA inactivation. Protein Sci 9:2161–2169.

56. Payne K, Sun Q, Sacchettini J, Hatfull GF. 2009. Mycobacteriophage Lysin B is a novel mycolylarabinogalactan esterase. Mol Microbiol 73:367–381.

57. Messing SAJ, Ton-Hoang B, Hickman AB, McCubbin AJ, Peaslee GF, Ghirlando R, Chandler M, Dyda F. 2012. The processing of repetitive extragenic palindromes: the structure of a repetitive extragenic palindrome bound to its associated nuclease. Nucleic Acids Res 40:9964–9979.

58. Mejdani M, Pawluk A, Maxwell KL, Davidson AR. 2021. Anti-CRISPR AcrIE2 binds the type I-E CRISPR-Cas complex but does not block DNA binding. J Mol Biol 433:166759.

59. Oliva MA, Martin-Galiano AJ, Sakaguchi Y, Andreu JM. 2012. Tubulin homolog TubZ in a phage-encoded partition system. Proc Natl Acad Sci U S A 109:7711–7716.

60. Singleton MR, Scaife S, Wigley DB. 2001. Structural analysis of DNA replication fork reversal by RecG. Cell 107:79–89.

61. Pereira CT, Roesler C, Faria JN, Fessel MR, Balan A. 2017. Sulfate-Binding Protein (Sbp) from Xanthomonas citri: Structure and Functional Insights. Mol Plant Microbe Interact 30:578–588.

62. Sheikh MA, Potter JA, Johnson KA, Sim RB, Boyd EF, Taylor GL. 2008. Crystal structure of VC1805, a conserved hypothetical protein from a Vibrio cholerae pathogenicity island, reveals homology to human p32. Proteins 71:1563–1571.

63. Ye K, Serganov A, Hu W, Garber M, Patel DJ. 2002. Ribosome-associated factor Y adopts a fold resembling a double-stranded RNA binding domain scaffold. Eur J Biochem 269:5182–5191.

64. Sato A, Watanabe T, Maki Y, Ueta M, Yoshida H, Ito Y, Wada A, Mishima M. 2009. Solution structure of the E. coli ribosome hibernation promoting factor HPF: Implications for the relationship between structure and function. Biochem Biophys Res Commun 389:580–585.

65. Paetzel M, Dalbey RE, Strynadka NCJ. 1998. Erratum: Crystal structure of a bacterial signal peptidase in complex with a β-lactam inhibitor. Nature 396:707–707.

66. Paetzel M, Dalbey RE, Strynadka NCJ. 2002. Crystal structure of a bacterial signal peptidase apoenzyme: implications for signal peptide binding and the Ser-Lys dyad mechanism. J Biol Chem 277:9512–9519.

67. Smith PA, Koehler MFT, Girgis HS, Yan D, Chen Y, Chen Y, Crawford JJ, Durk MR, Higuchi RI, Kang J, Murray J, Paraselli P, Park S, Phung W, Quinn JG, Roberts TC, Rougé L, Schwarz JB, Skippington E, Wai J, Xu M, Yu Z, Zhang H, Tan M-W, Heise CE. 2018. Optimized arylomycins are a new class of Gram-negative antibiotics. Nature 561:189–194.

68. Park JS, Kim HS, Park SH, Park MS, Kang S-M, Kim H-J, Han BW. 2019. Structural analyses of Helicobacter pylori FolC conducting glutamation in folate metabolism. Crystals (Basel) 9:429.

69. Smith CA, Cross JA, Bognar AL, Sun X. 2006. Mutation of Gly51 to serine in the P-loop of Lactobacillus casei folylpolyglutamate synthetase abolishes activity by altering the conformation of two adjacent loops. Acta Crystallogr D Biol Crystallogr 62:548–558.

70. Sun X, Cross JA, Bognar AL, Baker EN, Smith CA. 2001. Folate-binding triggers the activation of folylpolyglutamate synthetase. J Mol Biol 310:1067–1078.

71. Lim L, Molla G, Guinn N, Ghisla S, Pollegioni L, Vrielink A. 2006. Structural and kinetic analyses of the H121A mutant of cholesterol oxidase. Biochem J 400:13–22.

72. Sagermann M, Ohtaki A, Newton K, Doukyu N. 2010. Structural characterization of the organic solvent-stable cholesterol oxidase from Chromobacterium sp. DS-1. J Struct Biol 170:32–40.

73. Coulombe R, Yue KQ, Ghisla S, Vrielink A. 2001. Oxygen access to the active site of cholesterol oxidase through a narrow channel is gated by an Arg-Glu pair. J Biol Chem 276:30435–30441.

74. Cámara B, Liu M, Reynolds J, Shadrin A, Liu B, Kwok K, Simpson P, Weinzierl R, Severinov K, Cota E, Matthews S, Wigneshweraraj SR. 2010. T7 phage protein Gp2 inhibits the Escherichia coli RNA polymerase by antagonizing stable DNA strand separation near the transcription start site. Proc Natl Acad Sci U S A 107:2247–2252.

75. Shi K, Bohl TE, Park J, Zasada A, Malik S, Banerjee S, Tran V, Li N, Yin Z, Kurniawan F, Orellana K, Aihara H. 2018. T4 DNA ligase structure reveals a prototypical ATP-dependent ligase with a unique mode of sliding clamp interaction. Nucleic Acids Res 46:10474–10488.

76. Butt A, Higman VA, Williams C, Crump MP, Hemsley CM, Harmer N, Titball RW. 2014. The HicA toxin from Burkholderia pseudomallei has a role in persister cell formation. Biochem J 459:333–344.

77. Hickman AB, Li Y, Mathew SV, May EW, Craig NL, Dyda F. 2000. Unexpected structural diversity in DNA recombination. Mol Cell 5:1025–1034.

78. Crystal structure of the hydrogenase maturating endopeptidase HYBD from Escherichia coli1.

79. Smith N, Roitberg AE, Rivera E, Howard A, Holden MJ, Mayhew M, Kaistha S, Gallagher DT. 2006. Structural analysis of ligand binding and catalysis in chorismate lyase. Arch Biochem Biophys 445:72–80.

80. Gallagher DT, Mayhew M, Holden MJ, Howard A, Kim KJ, Vilker VL. 2001. The crystal structure of chorismate lyase shows a new fold and a tightly retained product. Proteins 44:304–311.

81. Sari N, He Y, Doseeva V, Surabian K, Ramprakash J, Schwarz F, Herzberg O, Orban J. 2007. Solution structure of HI1506, a novel two-domain protein from Haemophilus influenzae. Protein Sci 16:977–982.

82. Zhang R-G, Duke N, Laskowski R, Evdokimova E, Skarina T, Edwards A, Joachimiak A, Savchenko A. 2003. Conserved protein YecM from Escherichia coli shows structural homology to metal-binding isomerases and oxygenases. Proteins 51:311–314.

83. Cheng X, Zhang X, Pflugrath JW, Studier FW. 1994. The structure of bacteriophage T7 lysozyme, a zinc amidase and an inhibitor of T7 RNA polymerase. Proc Natl Acad Sci U S A 91:4034–4038.

84. Nguyen H-A, El Khoury T, Guiral S, Laaberki M-H, Candusso M-P, Galisson F, Foucher A-E, Kesraoui S, Ballut L, Vallet S, Orelle C, Zucchini L, Martin J, Page A, Attieh J, Aghajari N, Grangeasse C, Jault J-M. 2017. Expanding the kinome world: A new protein kinase family widely conserved in bacteria. J Mol Biol 429:3056–3074.

85. Teplyakov A, Obmolova G, Tordova M, Thanki N, Bonander N, Eisenstein E, Howard AJ, Gilliland GL. 2002. Crystal structure of the YjeE protein from Haemophilus influenzae: a putative Atpase involved in cell wall synthesis. Proteins 48:220–226.

86. Saridakis E, Giastas P, Efthymiou G, Thoma V, Moulis J-M, Kyritsis P, Mavridis IM. 2009. Insight into the protein and solvent contributions to the reduction potentials of [4Fe-4S]2+/+ clusters: crystal structures of the Allochromatium vinosum ferredoxin variants C57A and V13G and the homologous Escherichia coli ferredoxin. J Biol Inorg Chem 14:783–799.

87. Sun S, Geng L, Shamoo Y. 2006. Structure and enzymatic properties of a chimeric

bacteriophage RB69 DNA polymerase and single-stranded DNA binding protein with increased processivity. Proteins 65:231–238.

88. Shamoo Y, Friedman AM, Parsons MR, Konigsberg WH, Steitz TA. 1995. Crystal structure of a replication fork single-stranded DNA binding protein (T4 gp32) complexed to DNA. Nature 376:362–366.

89. Iwasaki T, Yamashita E, Nakagawa A, Enomoto A, Tomihara M, Takeda S. 2018. Three-dimensional structures of bacteriophage neck subunits are shared in Podoviridae, Siphoviridae and Myoviridae. Genes Cells 23:528–536.

90. Thiyagarajan N, Pham TTK, Stinson B, Sundriyal A, Tumbale P, Lizotte-Waniewski M, Brew K, Acharya KR. 2012. Structure of a metal-independent bacterial glycosyltransferase that catalyzes the synthesis of histo-blood group A antigen. Sci Rep 2:940.

91. Pan C, Zimmer A, Shah M, Huynh MS, Lai CC-L, Sit B, Hooda Y, Curran DM, Moraes TF. 2021. Actinobacillus utilizes a binding protein-dependent ABC transporter to acquire the active form of vitamin B6. J Biol Chem 297:101046.

92. Saha I, Ghosh B, Dasgupta J. 2024. Structural insights in to the atypical type-I ABC Glucose-6-phosphate importer VCA0625-27 of Vibrio cholerae. Biochem Biophys Res Commun 716:150030.

93. Naffin-Olivos JL, Daab A, White A, Goldfarb NE, Milne AC, Liu D, Baikovitz J, Dunn BM, Rengarajan J, Petsko GA, Ringe D. 2017. Structure Determination of Mycobacterium tuberculosis Serine Protease Hip1 (Rv2224c). Biochemistry 56:2304–2314.

94. Brooks CL, Ostrov DA, Schumann NC, Kakkad S, Li D, Peña K, Williams BP, Goldfarb NE. 2022. 2.1 Å crystal structure of the Mycobacterium tuberculosis serine hydrolase, Hip1, in its anhydro-form (Anhydrohip1). Biochem Biophys Res Commun 630:57–63.

95. Bray RC, Adams B, Smith AT, Bennett B, Bailey S. 2000. Reversible dissociation of thiolate ligands from molybdenum in an enzyme of the dimethyl sulfoxide reductase family·. Biochemistry 39:11258–11269.

96. Struwe MA, Kalimuthu P, Luo Z, Zhong Q, Ellis D, Yang J, Khadanand KC, Harmer JR, Kirk ML, McEwan AG, Clement B, Bernhardt PV, Kobe B, Kappler U. 2021. Active site architecture reveals coordination sphere flexibility and specificity determinants in a group of closely related molybdoenzymes. J Biol Chem 296:100672.

97. Bray RC, Adams B, Smith AT, Richards RL, Lowe DJ, Bailey S. 2001. Reactions of dimethylsulfoxide reductase in the presence of dimethyl sulfide and the structure of the dimethyl sulfide-modified enzyme. Biochemistry 40:9810–9820.

98. Panwar A, Martins BM, Sommer F, Schroda M, Dobbek H, Iobbi-Nivol C, Jourlin-Castelli C, Leimkühler S. 2024. Purification and electron transfer from soluble c-type cytochrome TorC to TorA for trimethylamine N-oxide reduction. Int J Mol Sci 25.

99. McAlpine AS, McEwan AG, Bailey S. 1998. The high resolution crystal structure of DMSO reductase in complex with DMSO 1 1Edited by D. C. Rees. J Mol Biol 275:613–623.

100. Prahlad J, Yuan Y, Lin J, Chang C-W, Iwata-Reuyl D, Liu Y, de Crécy-Lagard V, Wilson MA. 2020. The DUF328 family member YaaA is a DNA-binding protein with a novel fold. J Biol Chem 295:14236–14247.

101. Dey S, Biswas C, Sengupta J. 2018. The universally conserved GTPase HflX is an RNA helicase that restores heat-damaged Escherichia coli ribosomes. J Cell Biol 217:2519–2529.

102. Verma P, Ho R, Chambers SA, Cegelski L, Zimmer J. 2024. Insights into phosphoethanolamine cellulose synthesis and secretion across the Gram-negative cell envelope. Nat Commun 15:7798.

103. Golan G, Zharkov DO, Grollman AP, Dodson ML, McCullough AK, Lloyd RS, Shoham G. 2006. Structure of T4 pyrimidine dimer glycosylase in a reduced imine covalent complex with abasic site-containing DNA. J Mol Biol 362:241–258.

104. Morikawa K, Ariyoshi M, Vassylyev DG, Matsumoto O, Katayanagi K, Ohtsuka E. 1995. Crystal structure of a pyrimidine dimer-specific excision repair enzyme from bacteriophage T4: Refinement at 1.45 Å and X-ray analysis of the three active site mutants. J Mol Biol 249:360–375.

105. Vassylyev DG, Kashiwagi T, Mikami Y, Ariyoshi M, Iwai S, Ohtsuka E, Morikawa K. 1995. Atomic model of a pyrimidine dimer excision repair enzyme complexed with a DNA substrate: structural basis for damaged DNA recognition. Cell 83:773–782.

106. Kim I, Jeong M, Ka D, Han M, Kim N-K, Bae E, Suh J-Y. 2018. Solution structure and dynamics of anti-CRISPR AcrIIA4, the Cas9 inhibitor. Sci Rep 8:3883.

107. Cartmell A, Lowe EC, Baslé A, Firbank SJ, Ndeh DA, Murray H, Terrapon N, Lombard V, Henrissat B, Turnbull JE, Czjzek M, Gilbert HJ, Bolam DN. 2017. How members of the human gut microbiota overcome the sulfation problem posed by glycosaminoglycans. Proc Natl Acad Sci U S A 114:7037–7042.

108. Robb CS, Hobbs JK, Pluvinage B, Reintjes G, Klassen L, Monteith S, Giljan G, Amundsen C, Vickers C, Hettle AG, Hills R, Nitin, Xing X, Montina T, Zandberg WF, Abbott DW, Boraston AB. 2022. Metabolism of a hybrid algal galactan by members of the human gut microbiome. Nat Chem Biol 18:501–510.

109. Luis AS, Baslé A, Byrne DP, Wright GSA, London JA, Jin C, Karlsson NG, Hansson GC, Eyers PA, Czjzek M, Barbeyron T, Yates EA, Martens EC, Cartmell A. 2022. Author Correction: Sulfated glycan recognition by carbohydrate sulfatases of the human gut microbiota. Nat Chem Biol 18:1032.

110. Ségurel L, Ulaganathan TS, Mathieu S, Touvrey M, Poulet L, Drouillard S, Cygler M,

Helbert W. 2023. The porphyran degradation system is complete, phylogenetically and geographically diverse across the gut microbiota of East Asian populations. bioRxiv.

111. Berntsson RP-A, Odegrip R, Sehlén W, Skaar K, Svensson LM, Massad T, Högbom M, Haggård-Ljungquist E, Stenmark P. 2014. Structural insight into DNA binding and oligomerization of the multifunctional Cox protein of bacteriophage P2. Nucleic Acids Res 42:2725–2735.

112. Kwan JJ, Smirnova E, Khazai S, Evanics F, Maxwell KL, Donaldson LW. 2013. The solution structures of two prophage homologues of the bacteriophage λ Ea8.5 protein reveal a newly discovered hybrid homeodomain/zinc-finger fold. Biochemistry 52:3612–3614.

113. Oke M, Carter LG, Johnson KA, Liu H, McMahon SA, Yan X, Kerou M, Weikart ND, Kadi N, Sheikh MA, Schmelz S, Dorward M, Zawadzki M, Cozens C, Falconer H, Powers H, Overton IM, van Niekerk CAJ, Peng X, Patel P, Garrett RA, Prangishvili D, Botting CH, Coote PJ, Dryden DTF, Barton GJ, Schwarz-Linek U, Challis GL, Taylor GL, White MF, Naismith JH. 2010. The Scottish Structural Proteomics Facility: targets, methods and outputs. J Struct Funct Genomics 11:167–180.

114. Klink BU, Barden S, Heidler TV, Borchers C, Ladwein M, Stradal TEB, Rottner K, Heinz DW. 2010. Structure of Shigella IpgB2 in complex with human RhoA. J Biol Chem 285:17197–17208.

115. Kanamaru S, Uchida K, Nemoto M, Fraser A, Arisaka F, Leiman PG. 2020. Structure and function of the T4 Spackle protein Gp61.3. Viruses 12:1070.

116. Li Q, Zallot R, MacTavish BS, Montoya A, Payan DJ, Hu Y, Gerlt JA, Angerhofer A, de Crécy-Lagard V, Bruner SD. 2021. Epoxyqueuosine reductase QueH in the biosynthetic pathway to tRNA queuosine is a unique metalloenzyme. Biochemistry 60:3152–3161.

117. Hu Y, Jaroch M, Sun G, Dedon PC, de Crécy-Lagard V, Bruner SD. 2025. Mechanism of catalysis and substrate binding of epoxyqueuosine reductase in the biosynthetic pathway to queuosine-modified tRNA. Biochemistry 64:458–467.

118. Sintchak MD, Arjara G, Kellogg BA, Stubbe J, Drennan CL. 2002. The crystal structure of class II ribonucleotide reductase reveals how an allosterically regulated monomer mimics a dimer. Nat Struct Biol 9:293–300.

119. Niu Y, Yang L, Gao T, Dong C, Zhang B, Yin P, Hopp A-K, Li D, Gan R, Wang H, Liu X, Cao X, Xie Y, Meng X, Deng H, Zhang X, Ren J, Hottiger MO, Chen Z, Zhang Y, Liu X, Feng Y. 2020. A type I-F anti-CRISPR protein inhibits the CRISPR-Cas surveillance complex by ADP-ribosylation. Mol Cell 80:512–524.e5.

120. Djordjevic S, Goudreau PN, Xu Q, Stock AM, West AH. 1998. Structural basis for methylesterase CheB regulation by a phosphorylation-activated domain. Proc Natl Acad Sci U S A 95:1381–1386.

121. Velando F, Gavira JA, Rico-Jiménez M, Matilla MA, Krell T. 2020. Evidence for pentapeptide-dependent and independent CheB methylesterases. Int J Mol Sci 21:8459.

122. González-Montes L, Del Campo I, Garcillán-Barcia MP, de la Cruz F, Moncalián G. 2020. ArdC, a ssDNA-binding protein with a metalloprotease domain, overpasses the recipient hsdRMS restriction system broadening conjugation host range. PLoS Genet 16:e1008750.

123. Schnuchel A, Wiltscheck R, Eichinger L, Schleicher M, Holak TA. 1995. Structure of severin domain 2 in solution. J Mol Biol 247:21–27.

124. Bunting KA, Roe SM, Headley A, Brown T, Savva R, Pearl LH. 2003. Crystal structure of the Escherichia coli dcm very-short-patch DNA repair endonuclease bound to its reaction product-site in a DNA superhelix. Nucleic Acids Res 31:1633–1639.

125. Kim S, Kim TG, Byon HR, Shin H-J, Ban C, Choi HC. 2009. Recognition of single mismatched DNA using MutS-immobilized carbon nanotube field effect transistor devices. J Phys Chem B 113:12164–12168.

126. Jiang Y, Li F, Wu J, Shi Y, Gong Q. 2017. Structural insights into substrate selectivity of ribosomal RNA methyltransferase RlmCD. PLoS One 12:e0185226.

127. Jiang Y, Yu H, Li F, Cheng L, Zhu L, Shi Y, Gong Q. 2018. Unveiling the structural features that determine the dual methyltransferase activities of Streptococcus pneumoniae RlmCD. PLoS Pathog 14:e1007379.

128. Saha S, Kanaujia SP. 2024. Decoding substrate selectivity of an Archaeal RlmCD-like methyltransferase through its salient traits. Biochemistry 63:2477–2492.

129. Nuclear Magnetic Resonance Solution Structure of the Escherichia coli DNA Polymerase III θ Subunit.

130. Derose E, Kirby T, Mueller G, Chikova A, Schaaper R, London R. 2004. Phage like it HOTSolution structure of the bacteriophage P1-encoded HOT protein, a homolog of the θ subunit of E. coli DNA polymerase III. Structure 12:2221–2231.

131. Prokhorov DA, Mikoulinskaia GV, Molochkov NV, Uversky VN, Kutyshenko VP. 2015. High-resolution NMR structure of a Zn2+-containing form of the bacteriophage T5l-alanyl-d-glutamate peptidase. RSC Adv 5:41041–41049.

132. Vasina DV, Antonova NP, Gushchin VA, Aleshkin AV, Fursov MV, Fursova AD, Gancheva PG, Grigoriev IV, Grinkevich P, Kondratev AV, Kostarnoy AV, Lendel AM, Makarov VV, Nikiforova MA, Pochtovyi AA, Prudnikova T, Remizov TA, Shevlyagina NV, Siniavin AE, Smirnova NS, Terechov AA, Tkachuk AP, Usachev EV, Vorobev AM, Yakimakha VS, Yudin SM, Zackharova AA, Zhukhovitsky VG, Logunov DY, Gintsburg AL. 2024. Development of novel antimicrobials with engineered endolysin LysECD7-SMAP to combat Gram-negative bacterial infections. J Biomed Sci 31:75.

133. Owen RA, Fyfe PK, Lodge A, Biboy J, Vollmer W, Hunter WN, Sargent F. 2018. Structure and activity of ChiX: a peptidoglycan hydrolase required for chitinase secretion by Serratia marcescens. Biochem J 475:415–428.

134. El Omari K, Ren J, Bird LE, Bona MK, Klarmann G, LeGrice SFJ, Stammers DK. 2006. Molecular architecture and ligand recognition determinants for T4 RNA ligase. J Biol Chem 281:1573–1579.

135. Unciuleac M-C, Goldgur Y, Shuman S. 2017. Two-metal versus one-metal mechanisms of lysine adenylylation by ATP-dependent and NAD+-dependent polynucleotide ligases. Proc Natl Acad Sci U S A 114:2592–2597.

136. Fukai S, Nureki O, Sekine S-I, Shimada A, Tao J, Vassylyev DG, Yokoyama S. 2000. Structural basis for double-sieve discrimination of L-Valine from L-isoleucine and L-threonine by the complex of tRNAVal and valyl-tRNA synthetase. Cell 103:793–803.

137. Fukai S, Nureki O, Sekine S-I, Shimada A, Vassylyev DG, Yokoyama S. 2003. Mechanism of molecular interactions for tRNAVal recognition by valyl-tRNA synthetase. RNA 9:100–111.

138. Crystal Structure of LysB4, an Endolysin fromBacillus cereus-Targeting Bacteriophage B4.

139. Flayhan A, Vellieux FMD, Lurz R, Maury O, Contreras-Martel C, Girard E, Boulanger P, Breyton C. 2014. Crystal structure of pb9, the distal tail protein of bacteriophage T5: a conserved structural motif among all siphophages. J Virol 88:820–828.

140. Engilberge S, Riobé F, Di Pietro S, Lassalle L, Coquelle N, Arnaud C-A, Pitrat D, Mulatier J-C, Madern D, Breyton C, Maury O, Girard E. 2017. Crystallophore: a versatile lanthanide complex for protein crystallography combining nucleating effects, phasing properties, and luminescence. Chem Sci 8:5909–5917.

141. Engilberge S, Riobé F, Wagner T, Di Pietro S, Breyton C, Franzetti B, Shima S, Girard E, Dumont E, Maury O. 2018. Cover feature: Unveiling the binding modes of the crystallophore, a terbium-based nucleating and phasing molecular agent for protein crystallography (chem. Eur. J. 39/2018). Chemistry 24:9701–9701.

142. Vermersch PS, Tesmer JJ, Lemon DD, Quiocho FA. 1990. A Pro to Gly mutation in the hinge of the arabinose-binding protein enhances binding and alters specificity. Sugar-binding and crystallographic studies. J Biol Chem 265:16592–16603.

143. Schreier B, Stumpp C, Wiesner S, Höcker B. 2009. Computational design of ligand binding is not a solved problem. Proc Natl Acad Sci U S A 106:18491–18496.

144. Chitrakar I, Iuliano JN, He Y, Woroniecka HA, Tolentino Collado J, Wint JM, Walker SG, Tonge PJ, French JB. 2020. Structural basis for the regulation of biofilm formation and iron uptake in A. baumannii by the blue-light-using photoreceptor, BlsA. ACS Infect Dis 6:2592–2603.

145. Love MJ, Coombes D, Ismail S, Billington C, Dobson RCJ. 2022. The structure and function of modular Escherichia coli O157:H7 bacteriophage FTBEc1 endolysin, LysT84: defining a new endolysin catalytic subfamily. Biochem J 479:207–223.

146. Gullett JM, Cuypers MG, Grace CR, Pant S, Subramanian C, Tajkhorshid E, Rock CO, White SW. 2022. Identification of structural transitions in bacterial fatty acid binding proteins that permit ligand entry and exit at membranes. J Biol Chem 298:101676.

147. Cuypers MG, Subramanian C, Gullett JM, Frank MW, White SW, Rock CO. 2019. Acyl-chain selectivity and physiological roles of Staphylococcus aureus fatty acid-binding proteins. J Biol Chem 294:38–49.

148. Zhou Y, Bushweller JH. 2018. Solution structure and elevator mechanism of the membrane electron transporter CcdA. Nat Struct Mol Biol 25:163–169.

149. Pan H, Ho JD, Stroud RM, Finer-Moore J. 2007. The crystal structure of E. coli rRNA pseudouridine synthase RluE. J Mol Biol 367:1459–1470.

150. Nováček J, Šiborová M, Benešík M, Pantůček R, Doškař J, Plevka P. 2016. Structure and genome release of Twort-like Myoviridae phage with a double-layered baseplate. Proc Natl Acad Sci U S A 113:9351–9356.

151. Rutten L, Mannie J-PBA, Stead CM, Raetz CRH, Reynolds CM, Bonvin AMJJ, Tommassen JP, Egmond MR, Trent MS, Gros P. 2009. Active-site architecture and catalytic mechanism of the lipid A deacylase LpxR of Salmonella typhimurium. Proc Natl Acad Sci U S A 106:1960–1964.

152. Ramasamy S, Abrol R, Suloway CJM, Clemons WM Jr. 2013. The glove-like structure of the conserved membrane protein TatC provides insight into signal sequence recognition in twin-arginine translocation. Structure 21:777–788.

153. Rollauer SE, Tarry MJ, Graham JE, Jääskeläinen M, Jäger F, Johnson S, Krehenbrink M, Liu S-M, Lukey MJ, Marcoux J, McDowell MA, Rodriguez F, Roversi P, Stansfeld PJ, Robinson CV, Sansom MSP, Palmer T, Högbom M, Berks BC, Lea SM. 2012. Structure of the TatC core of the twin-arginine protein transport system. Nature 492:210–214.

154. McCaughey LC, Grinter R, Josts I, Roszak AW, Waløen KI, Cogdell RJ, Milner J, Evans T, Kelly S, Tucker NP, Byron O, Smith B, Walker D. 2014. Lectin-like bacteriocins from Pseudomonas spp. utilise D-rhamnose containing lipopolysaccharide as a cellular receptor. PLoS Pathog 10:e1003898.

155. Ghequire MGK, Garcia-Pino A, Lebbe EKM, Spaepen S, Loris R, De Mot R. 2013. Structural determinants for activity and specificity of the bacterial toxin LlpA. PLoS Pathog 9:e1003199.

156. Sborgi L, Verma A, Muñoz V, de Alba E. 2011. Revisiting the NMR structure of the

ultrafast downhill folding protein gpW from bacteriophage λ. PLoS One 6:e26409.

157. Pell LG, Gasmi-Seabrook GMC, Morais M, Neudecker P, Kanelis V, Bona D, Donaldson LW, Edwards AM, Howell PL, Davidson AR, Maxwell KL. 2010. The solution structure of the C-terminal Ig-like domain of the bacteriophage λ tail tube protein. J Mol Biol 403:468–479.

158. Pellegrino S, Radzimanowski J, de Sanctis D, Boeri Erba E, McSweeney S, Timmins J. 2012. Structural and functional characterization of an SMC-like protein RecN: new insights into double-strand break repair. Structure 20:2076–2089.

159. Newcomer RL, Schrad JR, Gilcrease EB, Casjens SR, Feig M, Teschke CM, Alexandrescu AT, Parent KN. 2019. The phage L capsid decoration protein has a novel OB-fold and an unusual capsid binding strategy. Elife 8.

160. Xie Y, Zhang L, Gao Z, Yin P, Wang H, Li H, Chen Z, Zhang Y, Yang M, Feng Y. 2022. AcrIF5 specifically targets DNA-bound CRISPR-Cas surveillance complex for inhibition. Nat Chem Biol 18:670–677.

161. Lundgren CAK, Sjöstrand D, Biner O, Bennett M, Rudling A, Johansson A-L, Brzezinski P, Carlsson J, von Ballmoos C, Högbom M. 2018. Scavenging of superoxide by a membrane-bound superoxide oxidase. Nat Chem Biol 14:788–793.

162. An SY, Ka D, Kim I, Kim E-H, Kim N-K, Bae E, Suh J-Y. 2020. Intrinsic disorder is essential for Cas9 inhibition of anti-CRISPR AcrIIA5. Nucleic Acids Res 48:7584–7594.

163. Moe E, Leiros I, Riise EK, Olufsen M, Lanes O, Smalås A, Willassen NP. 2004. Optimisation of the surface electrostatics as a strategy for cold adaptation of uracil-DNA N-glycosylase (UNG) from Atlantic cod (Gadus morhua). J Mol Biol 343:1221–1230.

164. Wang H-C, Hsu K-C, Yang J-M, Wu M-L, Ko T-P, Lin S-R, Wang AH-J. 2014. Staphylococcus aureus protein SAUGI acts as a uracil-DNA glycosylase inhibitor.

Nucleic Acids Res 42:1354–1364.

165. Kesharwani S, Raj P, Paul A, Roy K, Bhanot A, Mehta A, Gopal A, Varshney U, Gopal B, Sundriyal S. 2023. Crystal structures of non-uracil ring fragments in complex with Mycobacterium tuberculosis uracil DNA glycosylase (MtUng) as a starting point for novel inhibitor design: A case study with the barbituric acid fragment. Eur J Med Chem 258:115604.

166. Earl C, Bagnéris C, Zeman K, Cole A, Barrett T, Savva R. 2018. A structurally conserved motif in γ-herpesvirus uracil-DNA glycosylases elicits duplex nucleotide-flipping. Nucleic Acids Res 46:4286–4300.

167. Parker JB, Bianchet MA, Krosky DJ, Friedman JI, Amzel LM, Stivers JT. 2007. Enzymatic capture of an extrahelical thymine in the search for uracil in DNA. Nature 449:433–437.

168. Baños-Sanz JI, Mojardín L, Sanz-Aparicio J, Lázaro JM, Villar L, Serrano-Heras G, González B, Salas M. 2013. Crystal structure and functional insights into uracil-DNA glycosylase inhibition by phage Φ29 DNA mimic protein p56. Nucleic Acids Res 41:6761–6773.

169. NMR Structure of the 18 kDa Protein CC1736 From Caulobacter crescentus Identifies a Member of the “START” Domain Superfamily and Suggests Residues Mediating Substrate Specificity.

170. Pazos M, Peters K, Boes A, Safaei Y, Kenward C, Caveney NA, Laguri C, Breukink E, Strynadka NCJ, Simorre J-P, Terrak M, Vollmer W. 2020. SPOR proteins are required for functionality of class A penicillin-binding proteins in Escherichia coli. MBio 11.

171. Mueser TC, Jones CE, Nossal NG, Hyde CC. 2000. Bacteriophage T4 gene 59 helicase assembly protein binds replication fork DNA. The 1.45 Å resolution crystal structure reveals a novel α-helical two-domain fold 1 1Edited by P. E. Wright. J Mol Biol

296:597–612.

172. Inniss NL, Kochan TJ, Minasov G, Wawrzak Z, Chang C, Tan K, Shuvalova L, Kiryukhina O, Pshenychnyi S, Wu R, Dubrovska I, Babnigg G, Endres M, Anderson WF, Hauser AR, Joachimiak A, Satchell KJF. 2023. A structural systems biology approach to high-risk CG23 Klebsiella pneumoniae. Microbiol Resour Announc 12:e0101322.

173. Parsons LM, Yeh DC, Orban J. 2004. Solution structure of the highly acidic protein HI1450 from Haemophilus influenzae, a putative double-stranded DNA mimic. Proteins 54:375–383.

174. Yang C-S, Ko T-P, Chen C-J, Hou M-H, Wang Y-C, Chen Y. 2023. Crystal structure and functional implications of cyclic di-pyrimidine-synthesizing cGAS/DncV-like nucleotidyltransferases. Nat Commun 14:5078.

175. Whiteley AT, Eaglesham JB, de Oliveira Mann CC, Morehouse BR, Lowey B, Nieminen EA, Danilchanka O, King DS, Lee ASY, Mekalanos JJ, Kranzusch PJ. 2019. Bacterial cGAS-like enzymes synthesize diverse nucleotide signals. Nature 567:194–199.

176. Dönhöfer A, Franckenberg S, Wickles S, Berninghausen O, Beckmann R, Wilson DN. 2012. Structural basis for TetM-mediated tetracycline resistance. Proc Natl Acad Sci U S A 109:16900–16905.

177. Strohmeier M, Raschle T, Mazurkiewicz J, Rippe K, Sinning I, Fitzpatrick TB, Tews I. 2006. Structure of a bacterial pyridoxal 5'-phosphate synthase complex. Proc Natl Acad Sci U S A 103:19284–19289.

178. Bauer JA, Bennett EM, Begley TP, Ealick SE. 2004. Three-dimensional structure of YaaE from Bacillus subtilis, a glutaminase implicated in pyridoxal-5'-phosphate biosynthesis. J Biol Chem 279:2704–2711.

179. Liang L, Zhao H, An B, Tang L. 2018. High-resolution structure of podovirus tail adaptor suggests repositioning of an octad motif that mediates the sequential tail assembly. Proc Natl Acad Sci U S A 115:313–318.

180. Whelan F, Lafita A, Griffiths SC, Cooper REM, Whittingham JL, Turkenburg JP, Manfield IW, St John AN, Paci E, Bateman A, Potts JR. 2019. Defining the remarkable structural malleability of a bacterial surface protein Rib domain implicated in infection. Proc Natl Acad Sci U S A 116:26540–26548.

181. Leysen S, Vanderkelen L, Van Asten K, Vanheuverzwijn S, Theuwis V, Michiels CW, Strelkov SV. 2012. Structural characterization of the PliG lysozyme inhibitor family. J Struct Biol 180:235–242.

182. Leysen S, Vanderkelen L, Weeks SD, Michiels CW, Strelkov SV. 2013. Structural basis of bacterial defense against g-type lysozyme-based innate immunity. Cell Mol Life Sci 70:1113–1122.

183. Volpon L, Young CR, Matte A, Gehring K. 2006. NMR structure of the enzyme GatB of the galactitol-specific phosphoenolpyruvate-dependent phosphotransferase system and its interaction with GatA. Protein Sci 15:2435–2441.

184. Vo JL, Ortiz GCM, Totsika M, Lo AW, Hancock SJ, Whitten AE, Hor L, Peters KM, Ageorges V, Caccia N, Desvaux M, Schembri MA, Paxman JJ, Heras B. 2022. Variation of Antigen 43 self-association modulates bacterial compacting within aggregates and biofilms. NPJ Biofilms Microbiomes 8:20.

185. Aylett CHS, Izoré T, Amos LA, Löwe J. 2013. Structure of the tubulin/FtsZ-like protein TubZ from Pseudomonas bacteriophage ΦKZ. J Mol Biol 425:2164–2173.

186. Sterckx YGJ, Volkov AN, Vranken WF, Kragelj J, Jensen MR, Buts L, Garcia-Pino A, Jové T, Van Melderen L, Blackledge M, van Nuland NAJ, Loris R. 2014. Small-angle X-ray scattering- and nuclear magnetic resonance-derived conformational ensemble of

the highly flexible antitoxin PaaA2. Structure 22:854–865.

187. Kavanagh KL, Klimacek M, Nidetzky B, Wilson DK. 2002. Crystal structure of Pseudomonas fluorescens mannitol 2-dehydrogenase binary and ternary complexes. J Biol Chem 277:43433–43442.

188. Kim T-S, Patel SKS, Selvaraj C, Jung W-S, Pan C-H, Kang YC, Lee J-K. 2016. A highly efficient sorbitol dehydrogenase from Gluconobacter oxydans G624 and improvement of its stability through immobilization. Sci Rep 6:33438.

189. Habazettl J, Allan MG, Jenal U, Grzesiek S. 2011. Solution structure of the PilZ domain protein PA4608 complex with cyclic di-GMP identifies charge clustering as molecular readout. J Biol Chem 286:14304–14314.

190. Solution NMR structure of Pseudomonas Aeruginosa protein PA4608. Northeast Structural Genomics target PaT7.

191. Røhr AK, Hersleth H-P, Andersson KK. 2010. Tracking flavin conformations in protein crystal structures with Raman spectroscopy and QM/MM calculations. Angew Chem Int Ed Engl 49:2324–2327.

192. Williams C, Galyov EE, Bagby S. 2004. solution structure, backbone dynamics, and interaction with Cdc42 of Salmonella guanine nucleotide exchange factor SopE2. Biochemistry 43:11998–12008.

193. Liu B, Shadrin A, Sheppard C, Mekler V, Xu Y, Severinov K, Matthews S, Wigneshweraraj S. 2014. A bacteriophage transcription regulator inhibits bacterial transcription initiation by σ-factor displacement. Nucleic Acids Res 42:4294–4305.

194. Kudhair BK, Hounslow AM, Rolfe MD, Crack JC, Hunt DM, Buxton RS, Smith LJ, Le Brun NE, Williamson MP, Green J. 2017. Structure of a Wbl protein and implications for NO sensing by M. tuberculosis. Nat Commun 8:2280.

195. Schirmer T, de Beer TAP, Tamegger S, Harms A, Dietz N, Dranow DM, Edwards TE, Myler PJ, Phan I, Dehio C. 2021. Evolutionary diversification of host-targeted Bartonella effectors proteins derived from a conserved FicTA toxin-antitoxin module. Microorganisms 9:1645.

196. Hontz JS, Villar-Lecumberri MT, Potter BM, Yoder MD, Dreyfus LA, Laity JH. 2006. Differences in crystal and solution structures of the cytolethal distending toxin B subunit: Relevance to nuclear translocation and functional activation. J Biol Chem 281:25365–25372.

197. 1997. Corrigendum: A low energy short hydrogen bond in very high resolution structures of protein receptor–phosphate complexes. Nat Struct Biol 4:840–840.

198. Yao N, Ledvina PS, Choudhary A, Quiocho FA. 1996. Modulation of a salt link does not affect binding of phosphate to its specific active transport receptor. Biochemistry 35:2079–2085.

199. Ledvina PS, Tsai AL, Wang Z, Koehl E, Quiocho FA. 1998. Dominant role of local dipolar interactions in phosphate binding to a receptor cleft with an electronegative charge surface: equilibrium, kinetic, and crystallographic studies. Protein Sci 7:2550–2559.

200. Boorman J, Zeng X, Lin J, van den Akker F. 2024. Structural insights into peptidoglycan glycosidase EtgA binding to the inner rod protein EscI of the type III secretion system via a designed EscI-EtgA fusion protein. Protein Sci 33:e4930.

201. Bellinzoni M, Haouz A, Miras I, Magnet S, André-Leroux G, Mukherjee R, Shepard W, Cole ST, Alzari PM. 2014. Structural studies suggest a peptidoglycan hydrolase function for the Mycobacterium tuberculosis Tat-secreted protein Rv2525c. J Struct Biol 188:156–164.

202. Schumacher MA, Min J, Link TM, Guan Z, Xu W, Ahn Y-H, Soderblom EJ, Kurie JM,

Evdokimov A, Moseley MA, Lewis K, Brennan RG. 2012. Role of unusual P loop ejection and autophosphorylation in HipA-mediated persistence and multidrug tolerance. Cell Rep 2:518–525.

203. Schumacher MA, Piro KM, Xu W, Hansen S, Lewis K, Brennan RG. 2009. Molecular mechanisms of HipA-mediated multidrug tolerance and its neutralization by HipB. Science 323:396–401.

204. Rutten L, Geurtsen J, Lambert W, Smolenaers JJM, Bonvin AM, de Haan A, van der Ley P, Egmond MR, Gros P, Tommassen J. 2006. Crystal structure and catalytic mechanism of the LPS 3-O-deacylase PagL from Pseudomonas aeruginosa. Proc Natl Acad Sci U S A 103:7071–7076.

205. Allard P, Rak AV, Wimberly BT, Clemons WM Jr, Kalinin A, Helgstrand M, Garber MB, Ramakrishnan V, Härd T. 2000. Another piece of the ribosome: solution structure of S16 and its location in the 30S subunit. Structure 8:875–882.

206. Sauvage E, Powell AJ, Heilemann J, Josephine HR, Charlier P, Davies C, Pratt RF. 2008. Crystal structures of complexes of bacterial DD-peptidases with peptidoglycan-mimetic ligands: the substrate specificity puzzle. J Mol Biol 381:383–393.

207. Nicholas RA, Krings S, Tomberg J, Nicola G, Davies C. 2003. Crystal Structure of Wild-type Penicillin-binding Protein 5 from Escherichia coli. J Biol Chem 278:52826–52833.

208. Chen Y, Zhang W, Shi Q, Hesek D, Lee M, Mobashery S, Shoichet BK. 2009. Crystal structures of penicillin-binding protein 6 from Escherichia coli. J Am Chem Soc 131:14345–14354.

209. Brem J, Cain R, Cahill S, McDonough MA, Clifton IJ, Jiménez-Castellanos J-C, Avison MB, Spencer J, Fishwick CWG, Schofield CJ. 2016. Structural basis of metallo-β-lactamase, serine-β-lactamase and penicillin-binding protein inhibition by

cyclic boronates. Nat Commun 7:12406.

210. Nicola G, Tomberg J, Pratt RF, Nicholas RA, Davies C. 2010. Crystal structures of covalent complexes of β-lactam antibiotics with Escherichia coli penicillin-binding protein 5: toward an understanding of antibiotic specificity. Biochemistry 49:8094–8104.

211. Wang C, Xiao Q, Duan H, Li J, Zhang J, Wang Q, Guo L, Hu J, Sun B, Deng D. 2021. Molecular basis for substrate recognition by the bacterial nucleoside transporter NupG. J Biol Chem 296:100479.

212. Shin DH, Yokota H, Kim R, Kim S-H. 2002. Crystal structure of conserved hypothetical protein Aq1575 from Aquifex aeolicus. Proc Natl Acad Sci U S A 99:7980–7985.

213. Ghetu AF, Gubbins MJ, Frost LS, Glover JN. 2000. Crystal structure of the bacterial conjugation repressor finO. Nat Struct Biol 7:565–569.

214. Rooijakkers SHM, Milder FJ, Bardoel BW, Ruyken M, van Strijp JAG, Gros P. 2007. Staphylococcal complement inhibitor: structure and active sites. J Immunol 179:2989–2998.

215. Wang Z, Shen H, He B, Teng M, Guo Q, Li X. 2021. The structural mechanism for the nucleoside tri- and diphosphate hydrolysis activity of Ntdp from Staphylococcus aureus. FEBS J 288:6019–6034.

216. Yao W, Shi L, Liang D-C. 2007. Crystal structure of scaffolding protein CheW from thermoanaerobacter tengcongensis. Biochem Biophys Res Commun 361:1027–1032.

217. McGinnis RJ, Brambley CA, Stamey B, Green WC, Gragg KN, Cafferty ER, Terwilliger TC, Hammel M, Hollis TJ, Miller JM, Gainey MD, Wallen JR. 2022. A monomeric mycobacteriophage immunity repressor utilizes two domains to recognize an asymmetric DNA sequence. Nat Commun 13:4105.

218. Xiang Y, Morais MC, Cohen DN, Bowman VD, Anderson DL, Rossmann MG. 2008. Crystal and cryoEM structural studies of a cell wall degrading enzyme in the bacteriophage phi29 tail. Proc Natl Acad Sci U S A 105:9552–9557.

219. De Vitis V, Nakhnoukh C, Pinto A, Contente ML, Barbiroli A, Milani M, Bolognesi M, Molinari F, Gourlay LJ, Romano D. 2018. A stereospecific carboxyl esterase from Bacillus coagulans hosting nonlipase activity within a lipase-like fold. FEBS J 285:903–914.

220. Liu G, Shen Y, Atreya HS, Parish D, Shao Y, Sukumaran DK, Xiao R, Yee A, Lemak A, Bhattacharya A, Acton TA, Arrowsmith CH, Montelione GT, Szyperski T. 2005. NMR data collection and analysis protocol for high-throughput protein structure determination. Proc Natl Acad Sci U S A 102:10487–10492.

221. Liu G, Shen Y, Xiao R, Acton T, Ma LC, Joachimiak A, Montelione GT, Szyperski T. 2006. NMR structure of protein yqbG from Bacillus subtilis reveals a novel alpha-helical protein fold. Proteins 62:288–291.

222. Franke B, Veses-Garcia M, Diederichs K, Allison H, Rigden DJ, Mayans O. 2020. Structural annotation of the conserved carbohydrate esterase vb_24B_21 from Shiga toxin-encoding bacteriophage Φ24B. J Struct Biol 212:107596.

223. Unno M, Ishikawa-Suto K, Kusaka K, Tamada T, Hagiwara Y, Sugishima M, Wada K, Yamada T, Tomoyori K, Hosoya T, Tanaka I, Niimura N, Kuroki R, Inaka K, Ishihara M, Fukuyama K. 2015. Insights into the proton transfer mechanism of a bilin reductase PcyA following neutron crystallography. J Am Chem Soc 137:5452–5460.

224. Wada K, Hagiwara Y, Fukuyama K. 2011. One residue substitution in PcyA leads to unexpected changes in tetrapyrrole substrate binding. Acta Crystallogr A 67:C794–C795.

225. Hagiwara Y, Wada K, Irikawa T, Sato H, Unno M, Yamamoto K, Fukuyama K,

Sugishima M. 2016. Atomic-resolution structure of the phycocyanobilin:ferredoxin oxidoreductase I86D mutant in complex with fully protonated biliverdin. FEBS Lett 590:3425–3434.

226. Joutsuka T, Nanasawa R, Igarashi K, Horie K, Sugishima M, Hagiwara Y, Wada K, Fukuyama K, Yano N, Mori S, Ostermann A, Kusaka K, Unno M. 2023. Neutron crystallography and quantum chemical analysis of bilin reductase PcyA mutants reveal substrate and catalytic residue protonation states. J Biol Chem 299:102763.

227. Hagiwara Y, Sugishima M, Takahashi Y, Fukuyama K. 2006. Crystal structure of phycocyanobilin:ferredoxin oxidoreductase in complex with biliverdin IXalpha, a key enzyme in the biosynthesis of phycocyanobilin. Proc Natl Acad Sci U S A 103:27–32.

228. Insight into the Radical Mechanism of Phycocyanobilin−Ferredoxin Oxidoreductase (PcyA) Revealed by X-ray Crystallography and Biochemical Measurements. Insight into the Radical Mechanism of Phycocyanobilin−Ferredoxin Oxidoreductase (PcyA) Revealed by X-ray Crystallography and Biochemical Measurements.

229. Kohler AC, Gae DD, Richley MA, Stoll S, Gunn A, Lim S, Martin SS, Doukov TI, Britt RD, Ames JB, Lagarias JC, Fisher AJ. 2010. Structural basis for hydration dynamics in radical stabilization of bilin reductase mutants. Biochemistry 49:6206–6218.

230. Bowman SEJ, Backman LRF, Bjork RE, Andorfer MC, Yori S, Caruso A, Stultz CM, Drennan CL. 2019. Solution structure and biochemical characterization of a spare part protein that restores activity to an oxygen-damaged glycyl radical enzyme. J Biol Inorg Chem 24:817–829.

231. Rychlik MP, Chon H, Cerritelli SM, Klimek P, Crouch RJ, Nowotny M. 2010. Crystal structures of RNase H2 in complex with nucleic acid reveal the mechanism of RNA-DNA junction recognition and cleavage. Mol Cell 40:658–670.

232. Muroya A, Tsuchiya D, Ishikawa M, Haruki M, Morikawa M, Kanaya S, Morikawa K.

2001. Catalytic center of an archaeal type 2 ribonuclease H as revealed by X-ray crystallographic and mutational analyses. Protein Sci 10:707–714.

233. Chon H, Sparks JL, Rychlik M, Nowotny M, Burgers PM, Crouch RJ, Cerritelli SM. 2013. RNase H2 roles in genome integrity revealed by unlinking its activities. Nucleic Acids Res 41:3130–3143.

234. Takano K, Katagiri Y, Mukaiyama A, Chon H, Matsumura H, Koga Y, Kanaya S. 2007. Conformational contagion in a protein: structural properties of a chameleon sequence. Proteins 68:617–625.

235. Sivaraman J, Iannuzzi P, Cygler M, Matte A. 2004. Crystal structure of the RluD pseudouridine synthase catalytic module, an enzyme that modifies 23S rRNA and is essential for normal cell growth of Escherichia coli. J Mol Biol 335:87–101.

236. Yu F, Tanaka Y, Yamashita K, Suzuki T, Nakamura A, Hirano N, Suzuki T, Yao M, Tanaka I. 2011. Molecular basis of dihydrouridine formation on tRNA. Proc Natl Acad Sci U S A 108:19593–19598.

237. Cort JR, Koonin EV, Bash PA, Kennedy MA. 1999. A phylogenetic approach to target selection for structural genomics: solution structure of YciH. Nucleic Acids Res 27:4018–4027.

238. Gruenig MC, Lu D, Won SJ, Dulberger CL, Manlick AJ, Keck JL, Cox MM. 2011. Creating directed double-strand breaks with the ref protein. J Biol Chem 286:8240–8251.

239. Ben Bdira F, Qing L, Volkov AN, Erkelens M, Dame RT. 2021. Novel anti-repression mechanism of H-NS proteins by phage's "early proteins." Biophys J 120:18a.

240. van den Berg B. 2010. Crystal structure of a full-length autotransporter. J Mol Biol 396:627–633.

241. Chen K, Bonagura CA, Tilley GJ, McEvoy JP, Jung Y-S, Armstrong FA, Stout CD, Burgess BK. 2002. Crystal structures of ferredoxin variants exhibiting large changes in [Fe-S] reduction potential. Nat Struct Biol 9:188–192.

242. Schipke CG, Goodin DB, McRee DE, Stout CD. 1999. Oxidized and reduced Azotobacter vinelandii ferredoxin I at 1.4 Å resolution: Conformational change of surface residues without significant change in the [3Fe-4S]+/0 cluster. Biochemistry 38:8228–8239.

243. Meuwly M, Karplus M. 2004. Theoretical investigations on Azotobacter vinelandii ferredoxin I: effects of electron transfer on protein dynamics. Biophys J 86:1987–2007.

244. Xu Q, Göhler A-K, Kosfeld A, Carlton D, Chiu H-J, Klock HE, Knuth MW, Miller MD, Elsliger M-A, Deacon AM, Godzik A, Lesley SA, Jahreis K, Wilson IA. 2012. The structure of Mlc titration factor A (MtfA/YeeI) reveals a prototypical zinc metallopeptidase related to anthrax lethal factor. J Bacteriol 194:2987–2999.

245. Schureck MA, Dunkle JA, Maehigashi T, Miles SJ, Dunham CM. 2015. Defining the mRNA recognition signature of a bacterial toxin protein. Proc Natl Acad Sci U S A 112:13862–13867.

246. Chabbra A, Alzamora C, Ghali H, Handal M, Kathiria M, Martinek A, Nguyen J, Patel C, Patel S, Pham A, Lavin ES. 2019. Developing a physical model of HigB toxin and its endonuclease cleavage mechanism. FASEB J 33.

247. Benešík M, Nováček J, Janda L, Dopitová R, Pernisová M, Melková K, Tišáková L, Doškař J, Žídek L, Hejátko J, Pantůček R. 2018. Role of SH3b binding domain in a natural deletion mutant of Kayvirus endolysin LysF1 with a broad range of lytic activity. Virus Genes 54:130–139.

248. Lu JZ, Fujiwara T, Komatsuzawa H, Sugai M, Sakon J. 2006. Cell wall-targeting domain of glycylglycine endopeptidase distinguishes among peptidoglycan

cross-bridges. J Biol Chem 281:549–558.

249. Gonzalez-Delgado LS, Walters-Morgan H, Salamaga B, Robertson AJ, Hounslow AM, Jagielska E, Sabała I, Williamson MP, Lovering AL, Mesnage S. 2020. Two-site recognition of Staphylococcus aureus peptidoglycan by lysostaphin SH3b. Nat Chem Biol 16:24–30.

250. Frazão C, McVey CE, Amblar M, Barbas A, Vonrhein C, Arraiano CM, Carrondo MA. 2006. Unravelling the dynamics of RNA degradation by ribonuclease II and its RNA-bound complex. Nature 443:110–114.

251. Nagata R, Nishiyama M, Kuzuyama T. 2023. Substrate recognition mechanism of a trichostatin A-forming hydroxyamidotransferase. Biochemistry 62:1833–1837.

252. Pederick JL, Klose J, Jovcevski B, Pukala TL, Bruning JB. 2023. Escherichia coli YgiC and YjfC possess Peptide─Spermidine ligase activity. Biochemistry 62:899–911.

253. Zhang B, Lewis JA, Vermerris W, Sattler SE, Kang C. 2023. A sorghum ascorbate peroxidase with four binding sites has activity against ascorbate and phenylpropanoids. Plant Physiol 192:102–118.

254. Zhang T, Tamman H, Coppieters 't Wallant K, Kurata T, LeRoux M, Srikant S, Brodiazhenko T, Cepauskas A, Talavera A, Martens C, Atkinson GC, Hauryliuk V, Garcia-Pino A, Laub MT. 2022. Direct activation of a bacterial innate immune system by a viral capsid protein. Nature 612:132–140.

255. Sasnauskas G, Tamulaitiene G, Druteika G, Carabias A, Silanskas A, Kazlauskas D, Venclovas Č, Montoya G, Karvelis T, Siksnys V. 2023. TnpB structure reveals minimal functional core of Cas12 nuclease family. Nature 616:384–389.

256. d'Acapito A, Roret T, Zarkadas E, Mocaër P-Y, Lelchat F, Baudoux A-C, Schoehn G, Neumann E. 2023. Structural Study of the Cobetia marina Bacteriophage 1 (Carin-1) by

Cryo-EM. J Virol 97:e0024823.

257. Li D, Xiao Y, Fedorova I, Xiong W, Wang Y, Liu X, Huiting E, Ren J, Gao Z, Zhao X, Cao X, Zhang Y, Bondy-Denomy J, Feng Y. 2024. Single phage proteins sequester signals from TIR and cGAS-like enzymes. Nature 635:719–727.

258. Sigal N, Lichtenstein-Wolfheim R, Schlussel S, Azulay G, Borovok I, Holdengraber V, Elad N, Wolf SG, Zalk R, Zarivach R, Frank GA, Herskovits AA. 2024. Specialized Listeria monocytogenes produce tailocins to provide a population-level competitive growth advantage. Nat Microbiol 9:2727–2737.

259. Kejzar N, Laanto E, Rissanen I, Abrishami V, Selvaraj M, Moineau S, Ravantti J, Sundberg L-R, Huiskonen JT. 2022. Cryo-EM structure of ssDNA bacteriophage ΦCjT23 provides insight into early virus evolution. Nat Commun 13:7478.

260. Bad Phages in Good Bacteria: Role of the Mysterious orf63 of lambda and Shiga Toxin-Converting Phi 24 B Bacteriophages.

261. Zhou J, Wang L, Xiao H, Chen W, Liu Z, Song J, Zheng J, Liu H. 2025. In situ structures of the contractile nanomachine myophage Mu in both its extended and contracted states. J Virol 99:e0205624.

262. Peng Y, Tang H, Xiao H, Chen W, Song J, Zheng J, Liu H. 2024. Structures of mature and urea-treated empty bacteriophage T5: Insights into siphophage infection and DNA ejection. Int J Mol Sci 25:8479.

263. Lokareddy RK, Hou C-FD, Doll SG, Li F, Gillilan RE, Forti F, Horner DS, Briani F, Cingolani G. 2022. Terminase subunits from the Pseudomonas-phage E217. J Mol Biol 434:167799.

264. Debiasi-Anders G, Qiao C, Salim A, Li N, Mir-Sanchis I. 2025. Phage parasites targeting phage homologous recombinases provide antiviral immunity. Nat Commun

16:1889.

265. Jenson JM, Li T, Du F, Ea C-K, Chen ZJ. 2023. Ubiquitin-like conjugation by bacterial cGAS enhances anti-phage defence. Nature 616:326–331.

266. Valentová L, Füzik T, Nováček J, Hlavenková Z, Pospíšil J, Plevka P. 2024. Structure and replication of Pseudomonas aeruginosa phage JBD30. EMBO J 43:4384–4405.
